# Salvianolic acids are natural senolytics and increase lifespan in old age

**DOI:** 10.64898/2026.04.29.721790

**Authors:** Hongwei Zhang, Zhirui Jiang, Qiang Fu, Qixia Xu, Halidan Wufuer, Shuangbai Zhou, Qingfeng Li, Guilong Zhang, James L. Kirkland, Yu Sun

**Affiliations:** Center for Single-Cell Omics, School of Public Health, Shanghai Jiao Tong University School of Medicine, Shanghai, 200025, China; CAS Key Laboratory of Tissue Microenvironment and Tumor, Shanghai Institute of Nutrition and Health, Chinese Academy of Sciences, Shanghai 200031, China; School of Pharmacy, Institute of Aging Medicine, Shandong Medical and Pharmacological University, Yantai, Shandong 264003, China; Department of Plastic & Reconstructive Surgery, Shanghai Ninth People’s Hospital, Shanghai; Jiao Tong University School of Medicine, Shanghai 200011, China; Center for Advanced Gerotherapeutics, Cedars-Sinai Medical Center, Pacific Design Center, West Hollywood, CA 90069, USA; Division of Endocrinology, Diabetes and Metabolism, Cedars-Sinai Medical Center, Los Angeles, CA 90048, USA

## Abstract

Cellular senescence plays a critical role in chronological aging and is implicated in the onset and progression of multiple age-related disorders. Targeting senescent cells represents a promising strategy to reduce disease burden and improve healthspan. Here we report the senotherapeutic properties of salvianolic acid (SA) family members, specifically A (SAA), B (SAB) and E (SAE). From a natural medicinal agent library, we identified selective cytotoxicity of these SA compounds against senescent cells across diverse senescence types and cell lineages. Mechanistically, SAA, SAB and SAE enhance ROS production and target glutathione S-transferase Pi1 (GSTP1), a redox homeostasis modulator, inducing both apoptosis and ferroptosis in senescent cells. Incorporation of SAs into chemotherapeutic regimens enhanced anticancer efficacy and prolonged post-treatment survival. Intermittent SA administration improved physical function and increased healthspan and lifespan in aged mice. Collectively, our study establishes SAs as an emerging class of natural senolytics (phenolic acids) with the capacity to delay aging and alleviate age-related pathologies in advanced life.

## Introduction

Significant improvements in living conditions, development of public health and advances in modern medicine have fostered an unprecedented increase in median life expectancy worldwide. As a major risk factor for human health, aging represents a time-dependent and systemic decline of organ function that underlies the rise in age-related conditions, including cardiovascular diseases, metabolic disorders, neurodegenerative symptoms and various malignancies, a geriatric syndrome termed multimorbidity ^1, 2^. More than 30% of adults suffer from multimorbidity, with the prevalence rising among elderly populations, particularly in low- and middle-income countries ^3^. Identification of mechanisms shared by multimorbidities and development of evidence-based interventions is becoming the mainstay of aging-related studies. It is increasingly evident that addressing fundamental aging processes as a whole, rather than targeting individual pathologies, may be a more effective strategy for reducing age-related multimorbidities as a group to improve the healthspan in aging populations ^4, 5^.

Cellular senescence, a hallmark of aging, is a cell state characterized by stable cell cycle arrest first described in 1961 by Hayflick and Moorehead ^6^. Senescent cells arise in response to various inherent and/or environmental stimuli including telomeric attrition or dysfunction, DNA damage, oncogene activation, metabolic stress, cytotoxic chemotherapy and ionizing radiation ^7, 8^. Senescence is beneficial in a few pathophysiological cases, such as embryogenic patterning, tissue repair, wound healing and tumorigenesis prevention ^2^. However, senescent cells lingering in the tissue microenvironment of an aged host are mostly harmful through their senescence-associated secretory phenotype (SASP), which contributes to multiple pathological conditions throughout the lifespan and elevates age-related morbidity and mortality ^9^.

Senescent cells are intrinsically resistant to programmed cell death, although subject to *in vivo* surveillance and removal by immune cells ^10^. However, in the case of a dysfunctional immune system, senescent cells can accumulate and promote geriatric syndromes including frailty, cognitive dysfunction and immobility, and cause reduced physical resilience such as impaired recovery after infection or trauma ^11^. Therapeutic agents that selectively remove senescent cells, termed as senolytics, and drugs that attenuate the SASP, namely senomorphics, are being actively developed as a strategy to delay, prevent or alleviate age-related disorders. Activation of senescent cell anti-apoptotic pathways (SCAPs) has been recognized as a major mechanism regulating inherent resistance of senescent cells to death and an effective target for eliminating these cells ^12, 13^. To date, a handful of senolytic agents have been identified or developed with the potential to mitigate geriatric diseases ^11, 14^. Results from pilot clinical trials involving senolytics indicate the safety, tolerability and feasibility of reducing senescent cell burden in human patients ^15–18^, although much more remains to be done to validate the use of senolytics.

Naturally derived senolytics, specifically phytochemical compounds displaying senolytic activity, are being pursued, mainly due to their relatively high safety and low toxicity, but are scarce in nature. *Salvia miltiorrhiza* (*S. miltiorrhiza*), a perennial plant belonging to the genus *Salvia*, is a natural source of herbal medicine. Chemical constituents of *S. miltiorrhiza* have been identified, including more than 30 lipophilic compounds with diterpene quinone structures such as tanshinone I (Tan I) and tanshinone IIA (Tan IIA), as well as more than 50 hydrophilic compounds with phenolic acid structures such as salvianolic acid A (SAA) and rosmarinic acid (RA) ^19^. Salvianolic acids are among the most important bioactive components of *S. miltiorrhiza*, as they display antioxidant, anti-inflammatory, anticancer and other beneficial activities ^20^. Upon a thorough screening of a natural medicinal agent (NMA) library, we identified a group of *S. miltiorrhiza*-derived compounds specifically SAA, salvianolic acid B (SAB) and salvianolic acid E (SAE), as candidates with the capacity to selectively induce programmed death of senescent cells. In animal models, these SA agents remarkably improved chemotherapeutic efficacy and prolonged post-treatment survival of tumor-bearing mice. More importantly, SAs improved physical function of animals by depleting senescent cells from multiple solid organs. We propose natural compounds SAA, SAB and SAE as an emerging subgroup of senolytics, which are phenolic acids in nature and induce a dual death modality of senescent cells, thus holding the potential to intervene in age-related diseases.

## Results

### The SA family members A, B and E display senolytic activity upon preliminary screening of a NMA library

To identify small molecule chemicals with the potential to modulate senescent cells, an unbiased drug screen was performed with a library composed of 55 naturally-derived medicinal agents (NMA library) (Supplementary Table 1). We employed a primary normal human prostate stromal cell line, PSC27, as a cell-based model for *in vitro* screening. Composed predominantly of fibroblasts but with a minor percentage of non-fibroblast cell lineages such as smooth muscle cells and endothelial cells, PSC27 cells develop a SASP pattern upon exposure to exogenous stimuli including genotoxic chemotherapy and ionizing radiation ^21–24^. To induce senescence, we treated PSC27 cells with a pre-optimized sub-lethal dose of the genotoxic agent bleomycin (BLEO) and observed typical cellular senescence, as evidenced by increased staining positivity of senescence-associated β-galactosidase (SA-β-gal), decreased 5-bromo-2’-dexoyuridine (BrdU) incorporation and elevated DNA damage repair (DDR) foci several days afterwards (Supplementary Fig. 1a-c). There was a significant upregulation of the SASP and senescence markers after BLEO treatment (Supplementary Fig. 1d). We then developed a technical strategy to streamline the screening procedure and determine the effects individual agents generated on the survival and/or expression status of human senescent cells (Fig. 1a).

**Fig. 1.**
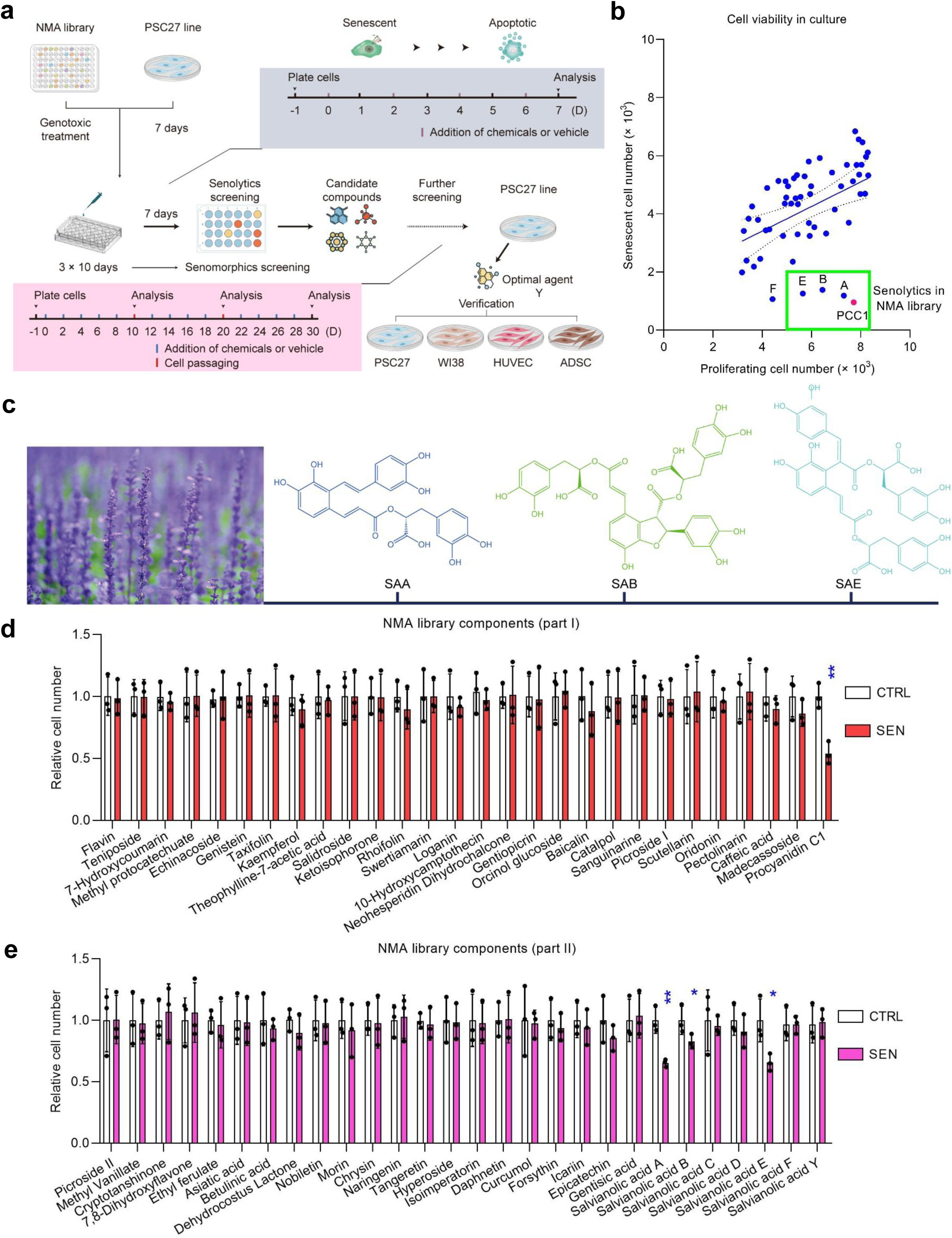
Workflow for *in vitro* screening of a NMA library to identify potential senotherapeutic candidates. (**a**) Schematic representation of the technical procedure for human cell model-based screening of a NMA library comprising 55 naturally derived agents. Screening was preliminarily performed for all natural products to evaluate their individual senotherapeutic activity, with the efficacy of potential candidates further validated in human stromal cell lines including PSC27, WI38, HUVECs and ADSCs. (**b**) Senolytic screening outputs after drug treatments in culture. Natural compounds were examined by incubation for 3 days at a concentration of 3 μg/ml/agent (PCC1 at 50 μM) with 5.0ⅹ10^3^ cells. Each blue dot indicates a hit based on effectiveness in selectively killing SEN (TIS) but not CTRL stromal cells, and represents the mean of 3 biological replicates. The green-edged rectangular region shows the resulting output of senolytic candidate screening. Note, the positive control, PCC1 (red dot), fell in this region. (**c**) A representative botanical image of *Salvia miltiorrhiza* and chemical structures of 3 senolytic Salvianolic acid molecules, namely SAA, SAB and SAE. Image source of *Salvia miltiorrhiza*, Pixabay.com (no permission or acknowledgement required). (**d**,**e**) Appraisal of the efficacy of natural agents as potential senolytics in SEN and their CTRL counterparts. (**d**), group part I. (**e**), group part II. NMA, natural medicinal agent. CTRL, control. SEN, senescent. TIS, therapy-induced senescence. PCC1, procyanidin C1. Data in **b**, **d** and **e** are shown as mean ± SD and representative of 3 independent biological replicates. *P* values were calculated by Student’s *t*-tests. ^, *P* > 0.05; *, *P* < 0.05; **, *P* < 0.01.

A major advantage of senolytic agents is their prominent capacity for selectively inducing programmed death of senescent cells, as exemplified by ABT-263 (navitoclax), ABT-737, combined use of dasatinib and quercetin (‘D + Q’) and the natural flavonoid procyanidin C1 (PCC1), the latter a senolytic flavonoid we already reported ^25–28^. To establish the potential of PSC27 as a valid cell model for senolytic screening, we first tested the efficacy of these geroprotective agents against senescent cells. Preliminary data indicated that each of these senolytics was able to effectively deplete senescent cells, but not their proliferating counterparts, thus validating the technical feasibility of using PSC27 cells for subsequent assays (Supplementary Fig. 1e). After screening the NMA library by experimentally scrutinizing individual components, we noticed that a subset of compounds was able to selectively kill senescent cells in culture (Fig. 1b; Supplementary Fig. 1f).

Notably, the naturally derived agents showing a capacity as potential senolytics all belonged to the salvianolic acid (SA) family. Similar to PCC1, a natural senolytic agent used as a positive control in these assays, several SA members including A, B, E and F (SAA, SAB, SAE and SAF, respectively, hereafter) displayed a strong capacity for reducing the number of senescent cells (Fig. 1c,d). Among them, however, SAF was excluded from further studies, mainly due to its cytotoxicity against both proliferating and senescent cells, implying potential adverse effects *in vivo* (Fig. 1b). In this study, we chose to focus on SAA, SAB and SAE, a subset of phytochemical agents that had been largely underexplored for senotherapeutic potential, although they had previously been reported to have various pharmacological activities including anti-atherosclerotic, cardioprotective, neuroprotective and anticancer effects ^29, 30^.

Despite the outstanding potential of a few agents in the NMA library to serve as effective senomorphics, as evidenced by their capacity in downregulating the expression of CXCL8, a key SASP factor (Supplementary Fig. 1g), we chose to focus on NMA components that exhibited senolytic activity, a highly valuable property that hitherto has been found only for a limited number of compounds, among which naturally-derived agents are even rarer in the current geroprotective arsenal.

### SA compounds are senolytic against senescent cells induced by various stimuli and across multiple cell lineages

The polyphenolic and hydrophilic compounds of SA family have been reported with various bioactivities, whereas their potential for targeting senescent cells appears to have remained unknown to date. We first assessed the competency and selectivity of SAA, SAB and SAE in eliminating senescent cells in culture conditions. Both SAA and SAB exhibited senolytic activity against senescent stromal cells once the concentration has reached 100 μM, while SAE displayed senolytic capacity starting even at the concentration of 50 μM. In contrast, proliferating cells remained largely unaffected under these conditions (Fig. 2a-c, Extended Data Fig. 1a-c and Extended Data Fig. 2a-c).

**Fig. 2.**
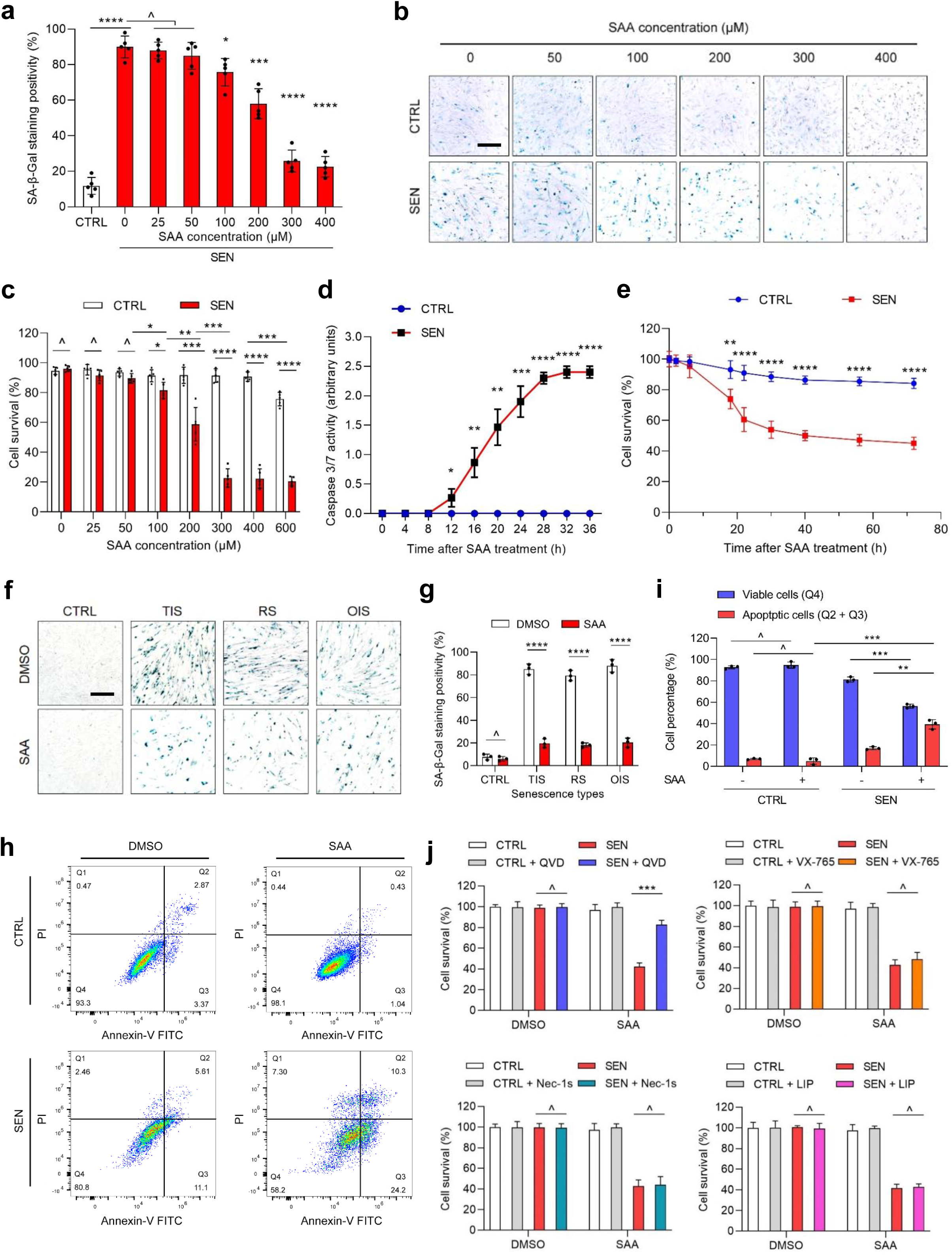
Characterization of the senolytic potential of SAA. (**a**) Measurement of senescent PSC27 survival by SA-β-gal staining. SAA was applied at increasing concentrations as indicated. CTRL, control. SEN, senescent. (**b**) Representative images from SA-β-gal staining assays to profile survival of cells including CTRL cells and their SEN counterparts induced by BLEO and treated with increasing concentrations of SAA. Scale bar, 50 μm. (**c**) Cell counting-based survival assay of CTRL and SEN cells exposed to increasing concentrations of SAA. (**d**) Cell apoptosis by caspase 3/7 activity measurement. SAA, 200 μM. (**e**) Cell viability in a time course to assess *in vitro* survival after SAA treatment (200 μM). (**f**) Representative images from SA-β-gal staining. TIS, therapy-induced senescence (by BLEO). RS, replicative senescence (by passive passaging). OIS, oncogene-induced senescence (by oncogenic *HRas^G12V^*). Scale bar, 50 μm. (**g**) Comparative statistics for SA-β-gal staining positivity as described in (**f**). (**h**) FACS sorting with an annexin V-FITC/PI staining to assay cell apoptosis. (**i**) Quantitative profiling of individual percentages of viable (Q4: PI^−^/annexin V^−^) and apoptotic (Q2 and Q3: PI^+^/annexin V^+^ and PI^−^/annexin V^+^, respectively) cells after treatment with vehicle or SAA for 3 days. (**j**) Survival analysis of CTRL and SEN cells after exposure to various agents targeting apoptosis, pyroptosis, necroptosis and ferroptosis. QVD, VX-765, Nec-1s and LIP are chemical inhibitors of apoptosis, pyroptosis, necroptosis and ferroptosis, respectively. QVD, QVD-Oph. VX-765, Belnacasan. Nec-1s, Necrostatin 2 racemate. LIP, lipoxstatin-1. Data in **a**, **c**, **d**, **e**, **g**, **i** and **j** are shown as mean ± SD and representative of 3 independent biological replicates with *P* values calculated by Student’s *t-*tests. ^, *P* > 0.05; *, *P* < 0.05; **, *P* < 0.01; ***, *P* < 0.001; ****, *P* < 0.0001.

Although higher concentrations of these compounds resulted in a relatively lower survival rate of senescent cells, with a threshold at approximately 300 μM, SAA and SAB exhibited cytotoxicity against proliferating cells only when used at 600 μM or higher (Fig. 2c and Extended Data Fig. 1c). The upper threshold of SAE for senolytic activity againist senescent cells was 200 μM, with toxicity against control cells starting from 300 μM (Extended Data Fig. 2c). Data from time-course assays of caspase 3/7 activity indicated that SAA, SAB and SAE caused apoptotic responses within 12 h, with the effect reaching a plateau 28 h after treatment (Fig. 2d, Extended Data Fig. 2d and Extended Data Fig. 3d). These results are generally consistent with data acquired from viability measurements (Fig. 2e, Extended Data Fig. 1e and Extended Data Fig. 2e). The senolytic nature of these SA compounds was subsequently confirmed in cells that were induced to become senescent by replicative exhaustion (RS) or oncogene *HRAS^G12V^* overexpression (OIS), senescence induction modalities that are alternative to TIS but share phenotypic similarities (Fig. 2f,g, Extended Data Fig. 1f,g, Extended Data Fig. 2f,g and Supplementary Fig. 2a-l) (Supplementary Table 2). Our data together suggest that SAA, SAB and SAE can selectively clear senescent human stromal cells induced by various stimuli in a dose-dependent manner, while non-senescent or proliferating cells remain largely unaffected when treated at the same concentrations of SA agents as proved to be effective against their senescent counterparts.

Given the preeminent capacity of SA compounds in selectively targeting PSC27 cells, which are of human prostate origin, we also tested human fetal lung fibroblasts (IMR90), primary human umbilical vein endothelial cells (HUVECs) and human adipose-derived mesenchymal stem cells (AT-MSC) with these phytochemicals. The results showed that senescent cells of all of these lineages had similar susceptibility to selective ablation by SA compounds, whereas non-senescent controls remained largely unaffected (Supplementary Fig. 3a-i) (Supplementary Table 3). We further confirmed by flow cytometry induction of apoptosis in senescent cells in response to SA agents, whereas proliferating cells remained basically viable (Fig. 2h,i, Extended Data Fig. 1h,i and Extended Data Fig. 2h,i). Taken together, these data suggest SA compounds have potential for selectively targeting senescent cells across various cell types and regardless of senescence inducers, by causing programmed cell death through apoptosis.

Apoptosis, necroptosis, ferroptosis and pyroptosis represent distinct programmed cell death mechanisms ^31^. To determine whether senescent cell clearance mediated by these SA compounds is exclusively through induction of apoptosis, or collaterally *via* other forms of programmed cell death, we chose to first treat cells with the pan-caspase apoptosis inhibitor quinolyl-valyl-O-methylaspartyl-[-2,6-difluorophenoxy]-methylketone (QVD-OPh). The capacity of SAA in killing senescent cells was markedly abrogated upon QVD-OPh treatment, suggesting that SAA executes its senolytic activity markedly through activation of an apoptotic response in senescent cells (Fig. 2j). To substantiate this, we performed further experiments. Data from assays involving relevant chemical inhibitors basically excluded the possibility of SAA-inducing cell death *via* pyroptosis or necroptosis. However, senescent cells were partially rescued when exposed to the ferroptosis-specific inhibitor Liproxstatin-1 (LIP), suggesting the involvement of ferroptotic pathway upon senescent cell death induced by SAA (Fig. 2j).

A similar pattern was observed in the case of SAB- or SAE-mediated cell death, which was mainly through apoptosis and ferroptosis of senescent cells (Extended Data Fig. 1j and Extended Data Fig. 2j). Thus, SA compounds, specifically SAA, SAB and SAE, a subset of phytochemical constituents derived from *S. miltiorrhiza*, hold the potential to induce senescent cell death *via* engagement of both apoptosis and ferroptosis. Although some natural agents such as ginsenoside RK1 can cause ferroptosis in cancer cells and inhibit tumor progression ^32^, the potential of targeting senescent cells with a natural compound *via* induction of ferroptosis either along or together with other forms of programmed cell death has been so far underexplored.

### SAs downregulate the SASP while engaging pro-apoptotic and pro-ferroptotic pathways in senescent cells

Given the efficacy of SA compounds for selectively inducing senescent cell death, we next queried the functional mechanism(s) supporting these actions. SAs and their precursors, including RA, caffeic acid ester derivatives, represent a group of water-soluble phenolic acids of *S. miltiorrhiza* bunge ^29, 33^. As the major bioactive compounds of *S. miltiorrhiza*, SAs display the capacity to inhibit cancer cell proliferation, metastasis and angiogenesis, while inducing their apoptosis and autophagy ^34^. Of note, SAs possess anti-inflammatory and antioxidant stress properties, and are effective in preventing lipid accumulation, mitochondrial dysfunction and ferroptosis ^35^. For example, SAA can inhibit lipid accumulation, reduce oxidative stress, and attenuate chronic inflammation and hepatic steatosis, thereby alleviating non-alcoholic fatty liver disease (NAFLD) in diabetic apoE^−/−^ mice fed a Western diet, mainly through activation of the AMPK and IGFBP-1 pathways ^35^. However, the molecular targets of SAs in senescent cells remain largely unclear. As a plant-derived secondary metabolite, SAA represents an important component of the hydrophilic constituents isolated from roots of *S. miltiorrhiza*, and displays structural features distinct from other SA compounds such as SAB and SAE (Fig. 3a, Extended Data Fig. 3a and Extended Data Fig. 4a).

**Fig. 3.**
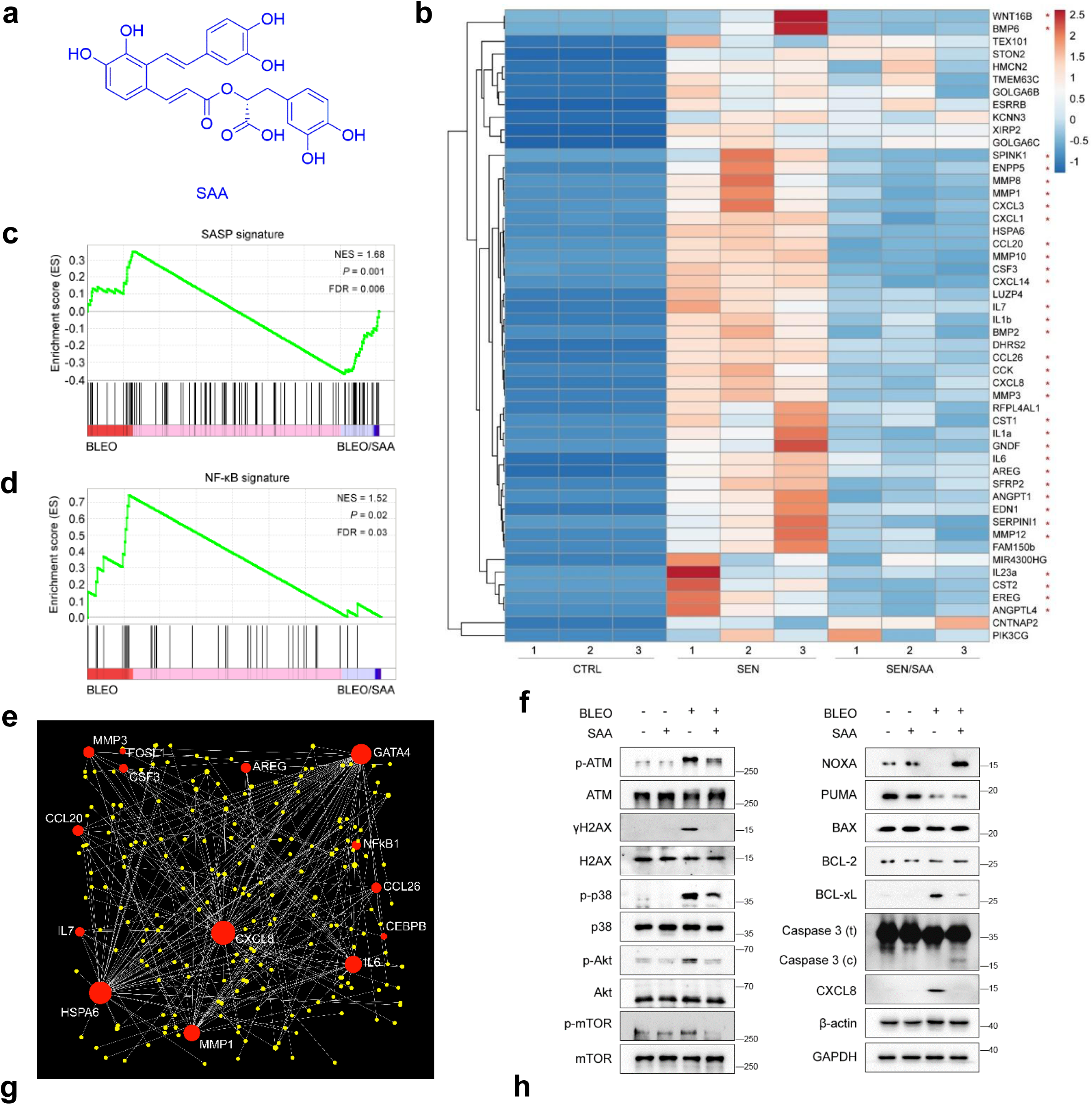
SAA induces senescent cell apoptosis by engaging pro-apoptotic pathways. (**a**) Chemical structure of SAA (CAS no. 96574-01-5). (**b**) Heatmap depicting top genes (50, by fold change) significantly upregulated in senescent PSC27 cells but downregulated by SAA treatment (50 μM). Red stars, SASP factors. (**c**) GSEA plot analysis of a significant gene set of the SASP spectrum upon treatment by BLEO *vs.* BLEO/SAA. (**d**) GSEA plot assessment of a significant gene set associated with NF-κB-mediated signaling upon treatment by BLEO *vs.* BLEO/SAA. (**e**) NetworkAnalyst map profiling protein-protein interactions of SASP-associated factors significantly upregulated in senescent cells but downregulated by SAA. (**f**) Immunoblots of PSC27 cells treated under different conditions involving BLEO and/or SAA. Expression of DDR signaling molecules, SASP regulatory factors, and pro-apoptotic and anti-apoptotic proteins was examined. Caspase 3 (t), total caspase 3; caspase 3 (c), cleaved caspase 3; p, phosphorylated. β-actin and GAPDH, loading controls. (**g**) Immunoblots of PSC27 cells exposed to different conditions involving BLEO and/or SAB. Examination of target molecules as described in (**f**) was performed. (**h**) Immunoblots of PSC27 cells exposed to different conditions involving BLEO and/or SAE. Examination of target molecules as described in (**f**) was conducted. Data are representative of 3 independent experiments. ^, *P* > 0.05; *, *P* < 0.05; **, *P* < 0.01; ***, *P* < 0.001.

We first determined the impact of SAA on the transcriptomic profile of senescent cells. Bioinformatics showed that 2089 genes were significantly downregulated, accompanied by upregulation of 1270 genes in senescent stromal cells upon SAA treatment (Supplementary Fig. 4a). Among these genes, we noticed a large array of SASP factors, which were upregulated upon cellular senescence, a tendency largely reversed by SAA (Fig. 3b). Network analysis indicated intricate mutual interactions or functional links between these ‘senescence-up and SAA-down’ molecules, which showed up on the top list of differentially expressed genes, many of which were secreted factors (Fig. 3c). Genome-wide GSEA mapping suggested downregulation of intracellular activities such as TNFα signaling, hypoxia, inflammatory response and DNA repair (Fig. 3d). Geneset-specific GSEA profiling indicated pronounced inhibition of both the SASP and NF-κB signatures by SAA treatment (Fig. 3e,f).

After mapping significantly changed transcripts to a gene ontology (GO) database comprising HPRD, Entrez Gene and UniProt accession identifiers ^36–38^, we found that the most prevalent biological processes of upregulated genes (top 200 selected as representatives) include cell communication, signal transduction, immune response, cell growth and/or maintenance (Supplementary Fig. 4b). The most evident cellular components of bioactive proteins encoded by these genes were those associated with plasma membrane, integral to plasma or organelle membrane, nucleus and endoplasmic reticulum (Supplementary Fig. 4c). Additionally, major molecular functions correlated with the most altered genes involve activities of G-protein coupled receptors, regular receptors, structural molecules and cell adhesion molecules (Supplementary Fig. 4d). Together, our data imply that SAA limits the expression of genes inherently associated with pro-inflammatory responses and secretory activity of senescent cells. The gene expression modulation pattern, SASP and NF-κB signature change, intermolecular function network and GO database profiling we observed in SAA-treated senescent cells were largely reproduced by datasets derived from bioinformatics analyses of SAB- and SAE-exposed senescent cells (Extended Data Fig. 3b-h and Extended Data Fig. 4b-h), suggesting a range of overlapping functions between these SA compounds, although they do not seem to share a pronounced similarity in chemical structure.

We then employed GSVA, a strategy providing increased power to detect subtle pathway activity changes over sample populations. There was a significant inactivation of pathway activities including antigen processing and presentation, cytokine-cytokine receptor interaction and inflammatory response (Fig. 3g), largely consistent with GSEA outputs. We also noticed increased activities of apoptosis and ferroptosis in SAA-treated senescent cells, suggesting a distinct cell death fate (Fig. 3g). To further dissect mechanisms underlying senescent cell death, we performed immunoblot assays and found that the DDR signaling appeared suppressed in senescent cells, as indicated by reduced levels of p-ATM and γH2AX (Fig. 3h and Supplementary Fig. 5a,b). Activation of the cellular stress sensor, p38 mitogen-activated protein kinase (MAPK), was observed in senescent cells, a pattern subject to reversal by SAA or reduction by SAB and SAE. The signaling pathway mediated by phosphorylation of Akt and mTOR was downregulated, implying SASP inhibition in these cells. We noticed increased expression of NOXA and/or PUMA, which are BCL-2 homology domain 3 (BH3)-only pro-apoptotic family members that functionally mediate programmed cell death (Fig. 3f-h). Although expression of BCL-2 remained unchanged in all cases, expression of BAX was elevated upon exposure of senescent cells to SAE, BCL-xL was decreased in cells treated by SAA or SAB, changes that potentially impair survival of senescent cells. We observed increased self-cleavage of caspase 3 in all groups, suggesting the engagement of mitochondria-mediated apoptosis, which usually involves cascade signaling by caspases. Of note, expression of glutathione peroxidase 4 (GPX4) and GTP cyclohydrolase 1 (GCH1) was markedly upregulated upon cellular senescence, but subject to reversal when cells were treated with SAs (Fig. 3g). This finding is consistent with data derived from our cell survival assays (Fig. 2j, Extended Data Fig. 1j and Extended Data Fig. 2j). As both GPX4 and GCH1 functionally antagonize ferroptosis, a non-apoptotic form of cell death typically characterized by ferrous ion (Fe^2+^)-dependent accumulation of lipid peroxides, our data suggest a distinct mechanism involving both apoptosis and ferroptosis correlated with reduced viability of senescent cells when treated by SAs, a subset of polyphenolic compounds isolated from *S. miltiorrhiza*.

GSVA-based proteomic data profiling further supported the induction of both apoptosis and ferroptosis of senescent cells after exposure to SA compounds (Extended Data Fig. 5b-d). This is consistent with the datasets of transcript-associated RNA-seq profiling and protein level analysis with immunoblot assays.

### Proteomics reveal GSTP1 as a major molecular target of SAs to mediate downstream cellular responses

Disruption of cellular homeostatic mechanisms essential for senescent cell survival seems to be a function shared by all three SA compounds. To establish how SAs induce programmed death of senescent cells, we chose to first screen for potential molecules that SAA, SAB and SAE commonly target. To this end, we generated a set of biotin-labeled compounds (SAA-bio, SAB-bio and SAE-bio) (Supplementary Fig. 6a-c) and measured the binding capacity of these complexes to recombinant proteins coated on 20K HuProt^TM^ human protein microarrays. We reasoned that proteins binding to these bio-compounds may represent direct targets of SA molecules and functionally mediate their biological activities. We applied bio-compounds to 20K HuProt^TM^ human protein microarrays (Supplementary Fig. 6a), and found the presence of 154 co-identified targets between the outputs of these bio-compounds (Supplementary Fig. 6b) (Supplementary Table 4). Pathway and process enrichment analysis indicated that the most outstanding GO items of these co-identified targets are cellular oxidant detoxification, cellular catabolic and carboxylic acid metabolic processes, activities inherently correlated with anti-oxidative and metabolic regulation (Supplementary Fig. 6c). It is noteworthy that among these proteins, 46 were top-ranked by statistical significance and linked to cellular catabolic process and cellular oxidant detoxification, activities that are hierarchically clustered on the enriched terms (Fig. 4a,b) (Supplementary Table 5).

**Fig. 4.**
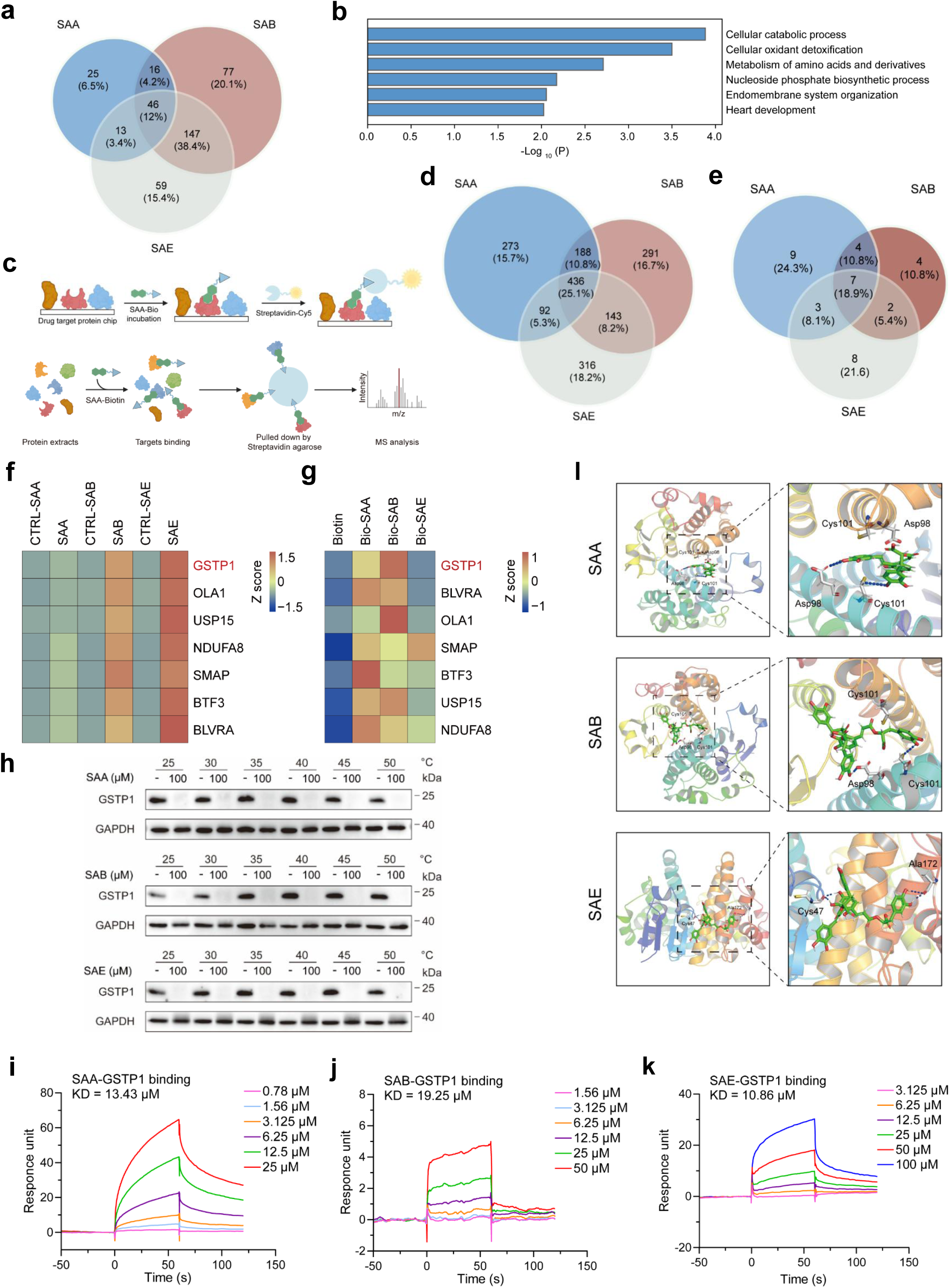
Proteomics profiling reveals potential factor(s) mediating the senolytic actions of SA compounds in senescent cells. (**a**) Venn diagram outlining the shared potential targets (46) of SAA, SAB and SAE identified by 20K HuProt^TM^ human protein microarrays as being significant (*P* < 0.05). (**b**) GO profiling of the 46 co-targets of SA compounds (SAA, SAB and SAE) identified by protein microarrays as significant. (**c**) A schematic illustration of technical procedure for protein microarray analysis (upper) and affinity-based MS that applied a SAA-Bio probe for target enrichment, followed by pulldown assays mediated by Cy5-Streptavidin-agarose (lower). Both strategies enable discovery of target(s) of a chemical using protein lysates from senescent cells. (**d**) Venn diagram depicting the potential co-targets (436) of SA compounds disclosed by pulldown assays. (**e**) Venn diagram depicting the potential co-targets (7) of SA compounds identified by both protein microarrays and pulldown assays. (**f**-**g**) A full list of the 7 co-targeted proteins of SA compounds as revealed in (**e**), after probing by protein microarrays (**f**) and pulldown assays (**g**). Note, GSTP1 (red highlighted) was inferred to be the major co-target of the 3 SA compounds. (**h**) CETSA assay evaluation of the thermal stabilization of GSTP1 by incubation with SA compounds over a gradient of temperatures from 25 to 50°C in protein lysates of PSC27 cells. (**i**-**k**) Demonstration that SA compounds physically bind to GSTP1. In the SPR assay, GSTP1 was treated by each of the SA compounds (**i**, SAA; **j**, SAB; **k**, SAE) over a range of concentrations as indicated in the individual plots. Kinetically derived binding affinity (KD) values are shown beside the traces. (**l**) Molecular docking that illustrates amino acid residues Cys101 and others (such as Asp98) of GSTP1 bind to SAA and SAB (S=-6.817 kcal/mol; S=-8.304 kcal/mol), or alternatively, amino acid residues Cys47 and others (such as Ala172) of GSTP1 bind to SAE (S=-6.762 kcal/mol). Reported X-ray diffraction microscopy structure (PDB ID: 2A2R) was applied to perform molecular docking, with the key amino acid residues highlighted in structure. MS, mass spectrometry. ^, *P* > 0.05; *, *P* < 0.05; **, *P* < 0.01.

Further assessment performed with biotin-based pulldown assays involving streptavidin-agarose beads followed by MS profiling showed that there are 436 proteins physically bound by all the 3 biotin-labeled compounds (Fig. 4c,d) (Supplementary Table 6). Upon performance of an overall screening of the targets identified by both human protein microarrays and pulldown/MS analysis, we noticed there were 7 targets shared by outputs of these strategies (Fig. 4e and Supplementary Fig. 6d-f). Among them was glutathione S-transferase Pi1 (GSTP1), an isozyme responsible for redox homeostasis maintenance *via* detoxification and anti-oxidation in cellular oxidative stress responses including senescence *per se* ^39, 40^ (Fig. 4f,g). Being a homodimeric protein of ∼46 kDa (Supplementary Fig. 6g), GSTP1 acts as a key regulator of redox homeostasis, while a potent inducer of cellular senescence, tryptanthrin, can directly bind GSTP1 and inhibit its enzymatic activity, causing ROS accumulation and DDR exacerbation, events that sensitize senescent cells to apoptosis induced by ABT-263 ^39^. However, whether GSTP1 is a co-target of SA compounds and mediates senolysis remains basically unknown.

We then queried the biophysical properties associated with interactions of SA compounds with GSTP1. Cellular thermal shift assay (CETSA), which assesses protein thermal stabilization after ligand binding at the molecular level, was performed over a temperature range from 25-50 °C. Interestingly, a substantial decrease of the stability of recombinant human GSTP1 (rhGSTP1) upon incubation with each SA compound was observed, in sharp contrast to the vehicle-treated (Fig. 4h). Although GSTP1 expression remained largely unchanged at transcription level, the overall activity significantly reduced upon senescence, and further declined upon treatment by SAs (Fig. 4h,i). The decline of GSTP1 protein signal in the presence of SA compounds is consistent with its activity loss, suggesting that physical interaction with these agents may lead to reduced stability, potentiated by degradation of GSTP1 during *in vitro* incubation. Alternatively, results from surface plasmon resonance (SPR) assay, a label-free direct optical biosensor strategy, indicated a potent binding of GSTP1 in the presence of SA compounds, yielding a predicted dissociation constant (KD) for each agent (1.88 µM, 1.15 µM and 1.32 µM for SAA, SAB and SAE, respectively) (Fig. 4k-m).

To identify the residue(s) of GSTP1 responsible for its physical interaction with SA compounds, we performed *in silico* simulation modelling by molecular docking. The results suggest that both SAA and SAB have direct hydrogen bond connections with GSTP1 *via* certain residues, specifically Cys101 (Fig. 4n, upper and middle illustrations). Alternatively, we noticed that SAE has direct hydrogen bond connections with GSTP1 *via* its residue Cys47, which we speculate serves as a major site (Fig. 4n, lower illustration). Previous studies indicate that GSTP1 has two solvent accessible cysteine residues, which can affect its catalytic activity once genetically modified. Among them, Cys47 is located near the binding site for reduced glutathione (G-site) and is critical for maintaining conformation and stability of the G-site, while Cys101 is located at the dimer interface and forms a disulfide bridge with Cys47, necessitating a conformational change of the active site to inactivate the enzyme ^41^.

As molecular docking simulation indicated proximity of Cys101 to SAA and SAB, which bind within the enzymatic pocket of GSTP1, we queried whether this residue participates in SAA or SAB binding. We further hypothesized that SAA and SAB might competitively inhibit glutathione (GSH) binding to GSTP1, potentially suppressing its enzymatic activity, or alternatively, cause GSTP1 degradation. We performed site-directed mutagenesis to generate a Cys101Ser (C101S) variant, with the recombinant GSTP1 protein linked to a GFP tag. Our study indicated that mutation of C101S compromised binding of GSTP1 with SAA (Supplementary Fig. 8a), suggesting that Cys101 mediates the interaction between GSTP1 and SAA. Notably, a similar pattern was observed in the case of GSTP1 binding by SAB (Supplementary Fig. 8b). To probe the GSTP1-SAE interaction, we examined whether the Cys47Ser (C47S) mutation significantly alters the binding capacity of the enzyme with SAE. Our data suggest that Cys47 is essential for SAE to bind GSTP1, as evidenced by elevated KD or reduced binding affinity between these molecules (Supplementary Fig. 8c).

Recent studies reported a GPX4/FSP1-independent cellular defense mechanism against ferroptosis, while SMURF2-mediated GSTP1 ubiquitination and degradation, or chemical inhibition of GSTP1’s catalytic activity can sensitize cancer cells to ferroptosis ^42^. However, whether directly targeting GSTP1 can induce programmed death of senescent cells remains unclear. To address this, we used LAS17, a potent and selective tyrosine-directed irreversible inhibitor of GSTP1 activity ^43^, to treat senescent cells. Upon exposure to LAS17 in culture, senescent cells displayed significantly reduced survival rate, in contrast to parallel controls (Extended Data Fig. 5a). Of note, this tendency was significantly, albeit incompletely reversed in the presence of QVD or LIP, suggesting the involvement of both apoptosis and ferroptosis during senescent cell death induced by GSTP1-specific targeting. These results were essentially reproduced by data from treatment with Ezatiostat, a tripeptide analog of glutathione to act as prodrug inhibitor for GSTP1 ^44^. Collectively, the data are basically consistent with our finding that SA compounds directly bind to the antioxidant enzyme GSTP1 and affect its stability and/or function in senescent cells, eventually causing their increased death rate. Therefore, GTSP1 is an antioxidant enzyme in senescent cells and has an essential role in maintaining their survival, although its relationship to other molecules in the SCAP network of senescent cells merits further investigation.

### SAs cause mitochondrial dysfunction and induce apoptosis and ferroptosis in senescent cells

Since SA family members generally enhance cell viability, restrain ROS production and reduce oxidative stress in mammalian cells ^29, 45^, we next asked whether similar or antioxidant effects can be observed in senescent cells exposed to SAs. We first examined the influence of SA compounds on intracellular ROS levels or ROS production capacity of PSC27 sublines. Upon senescence induction by BLEO, we observed elevated ROS biosynthesis, as evidenced by increased signals of 2’,7’-dichlorodihydrofluorescein diacetate (DCFH-DA), a fluorescent probe used to monitor free radical reactions (Fig. 5a,b). Of note, treatment by each of SAA, SAB and SAE further augmented DCFH-DA signal intensity, rather than decreasing it. This is in contrast to previously reported effects of SA molecules, which can dampen intracellular oxidation, counteract oxidative stress and alleviate apoptosis ^46, 47^. More importantly, we noticed increased activity of caspase 3 cleavage in senescent cells upon exposure to SAs, as exemplified by the case of SAA treatment (Fig. 5c). Elevated nuclear translocation of p53 was observed in senescent cells, a tendency that was reversed upon exposure to SAA, consistent with previous studies demonstrating nuclear exclusion of p53 as a key step in senescent cell apoptosis ^48^.

**Fig. 5.**
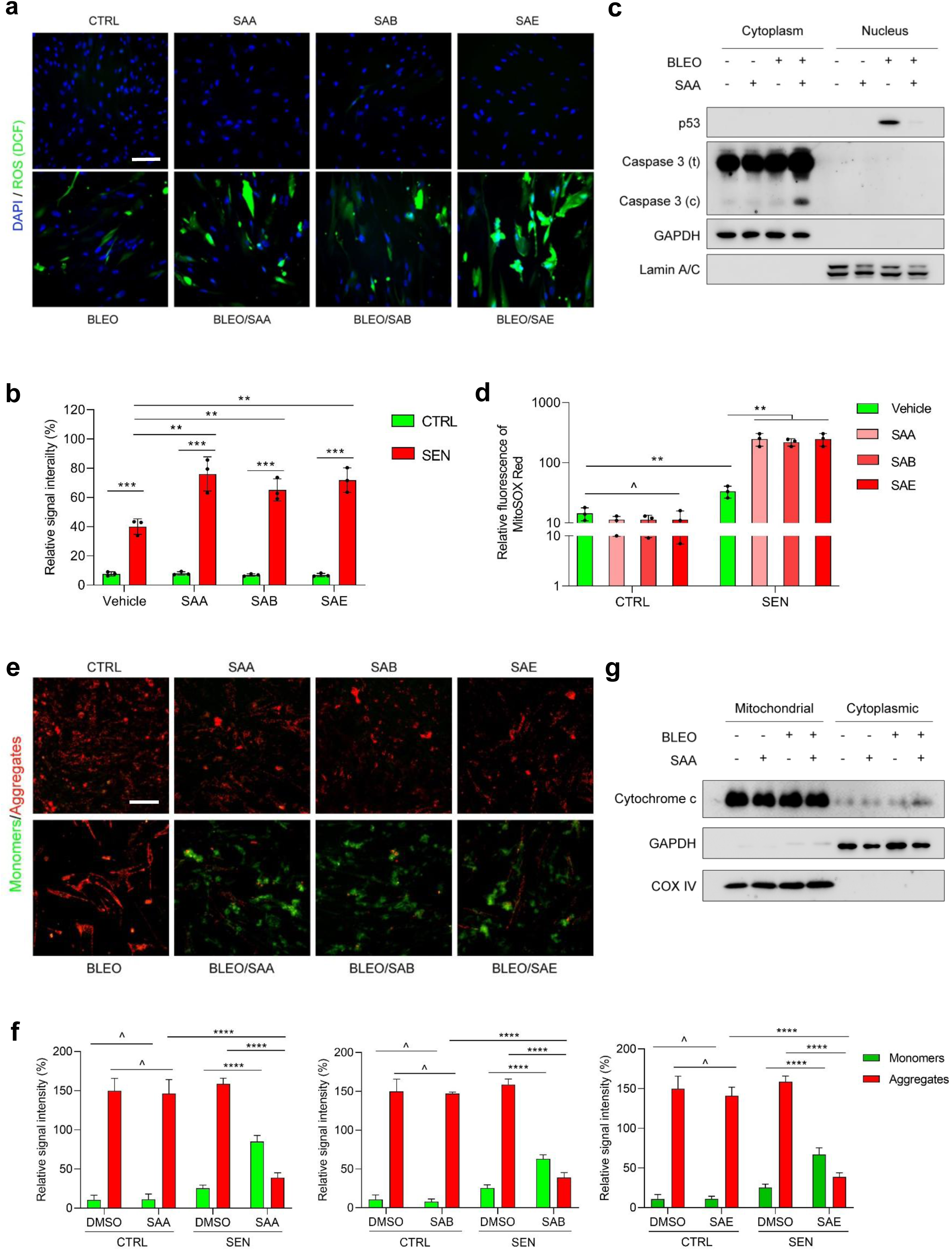
SA compounds eliminate senescent cells by enhancing intracellular ROS production and disturbing mitochondrial membrane potential. (**a**) Representative images during measurement of ROS production with 2′,7′ dichlorofluorescein diacetate (DCFH-DA), a cell-permeable fluorescent probe that labels intracellular ROS molecules. Experiments were performed 1 d after treatment with each of the SA compounds. DCF, dichlorodihydrofluorescein. Scale bar, 20 μm. (**b**) Comparative statistics of the relative signal intensity *per* treatment group as described in **a**. (**c**) Immunoblot analysis of cytoplasmic and nuclear lysates of PSC27 cells treated under different conditions involving BLEO and/or SAA. GAPDH and Lamin A/C, loading controls for cytoplasmic and nuclear samples, respectively. (**d**) Fluorescence signal measurement of MitoSOX Red, a mitochondrial superoxide indicator, in PSC27 cells under different conditions. (**e**) Representative images from assessment of mitochondrial membrane potential (Δψm) by staining with a fluorescent dye JC-1. Signals were measured over 3 d. Green fluorescence indicates JC-1 monomers, which appear in cytosol after mitochondrial membrane depolarization and indicate early-stage apoptosis. Red fluorescence indicates JC-1 aggregates, which reside on intact mitochondria. Scale bar, 20 μm. (**f**) Comparative statistics of the relative signal intensity *per* treatment group as described in **e**. SAA (left), SAB (middle) and SAE (right) were employed at 100 μM, 100 μM and 50 μM in relevant assays, respectively. (**g**) Immunoblot analysis of mitochondrial and cytoplasmic lysates of PSC27 cells treated under different conditions involving BLEO and/or SAA. GAPDH and COX IV, loading controls for cytoplasmic and mitochondrial samples, respectively. Data in **b**, **d** and **f** are shown as mean ± SD and representative of 3 independent biological replicates, with *P* values calculated by Student’s *t*-tests. ^, *P* > 0.05; *, *P* < 0.05; **, *P* < 0.01; ***, *P* < 0.001; ****, *P* < 0.0001.

Senescent cells have mitochondrial dysfunction and metabolic reprogramming, events responsible for oxidative stress and production of reactive oxygen species (ROS) including superoxide ^49^. We employed MitoSOX Red, a cell permeable probe to measure the production capacity of mitochondrial ROS especially superoxide, and observed a significantly increased level of mitochondrial ROS in senescent cells, but not their proliferating counterparts, upon treatment by SA compounds (Fig. 5d). These data suggest compromised ability to maintain homeostasis in senescent cells treated with SAs, a case distinct from that of non-senescent cells and reminiscent of the effect of other senolytic agents, such as the natural flavonoid PCC1 ^28^.

Mitochondrial dysfunction is inherently linked with cellular senescence and aging phenotypes, as evidenced by compromised mitochondrial membrane potential (Δψm) and elevated mitochondrial ROS levels ^50^. Specifically, reduction of Δψm represents an intracellular physiological dysfunction that can trigger apoptosis *via* the mitochondrial-mediated intrinsic pathway ^51^. A significant decrease of Δψm was observed in senescent cells, although control cells remained largely unaffected upon exposure to SAs, as evidenced by data from JC-1 assays (Fig. 5e, f). However, the pattern of Δψm decline in senescent cells appeared further exacerbated when cells were exposed to SAs, as suggested by an increased number of monomers but fewer aggregates (Fig. 5e,f).

To validate these apoptosis-related changes, we examined extent of cytochrome c release, an event associated with apoptosis and that can act as a direct apoptotic driver ^51^. Immunoblots with lysates SA-treated cells indicated that exposure of senescent cells to SAs such as SAA enhanced cytochrome c release from mitochondria to their surrounding cytoplasmic space (Fig. 5g). Indeed, the release of cytochrome c from mitochondria is basically in line with caspase activation in SAA-treated senescent cells (Fig. 2h-j). Therefore, SA compounds augment ROS production, cause Δψm loss and promote cytochrome c release from mitochondria in senescent cells, events linked with mitochondrial disability and programmed cell death at least *via* apoptosis.

As our earlier data suggest that SAs may induce both apoptosis and ferroptosis in senescent cells, we next examined the underlying mechanisms. Induction of senescence or treatment by SAs did not alter the protein level of GSTP1, as evidenced by immunoblot (Extended Data Fig. 5e). However, the p-JNK/p-c-Jun pathway, which can functionally mediate ferroptosis ^52, 53^, was activated upon senescence induction, with the activation further enhanced after SA treatment. In contrast, GPX4 expression showed a different alteration pattern, with the signal remarkably elevated in senescent cells but reduced upon SA treatment. Further data suggest that GPX4 is subject to normal intracellular turnover and excluded the ubiquitin/proteasome- or ribosome/autophagy-dependent degradation of GPX4, suggesting that the transcription-based regulation plays a major role in GPX4 expression in senescent cells (Extended Data Fig. 5f-h). Senescent cells exhibited a marked decline of reduced glutathione/oxidized glutathione (GSH/GSSG) ratio, which further decreased upon SA treatment (Extended Data Fig. 5i). Immunoprecipitation assay indicated a direct physical interaction of GSTP1 with JNK in senescent cells, which was weakened upon SA treatment (Extended Data Fig. 5j). This is consistent with activation of p-c-Jun signaling in SA-treated senescent cells (Extended Data Fig. 5e) and effect of ferroptosis-inducing compounds on phosphorylation of c-Jun, a core component of the activating protein-1 (AP-1) transcription factor complex, which regulates cellular processes including proliferation, survival and death ^54, 55^. As AP-1 can compete with certain transcription factors at the enhancer/promoter region of genes, we observed downregulated GPX4 expression in senescent cells treated by SA compounds. Data from cloning of GPX4 enhancer/promoter into a luciferase reporter vector supported that GPX4 is negatively regulated at transcriptional level by JNK/c-Jun pathway (Extended Data Fig. 5k).

We noticed that senescent cells displayed higher levels of intracellular Fe^2+^ (Supplementary Fig. 9a), which contributes to lipid oxidation. The signal intensity of intracellular Fe^2+^ was not changed upon treatment with SA compounds such as SAA, erastin or Lip1 (activator and inhibitor of ferroptosis, respectively), but was reduced by an iron chelator deferoxamine mesylate (**DFO**), suggesting the senescent cell-related vulnerability to ferroptosis was ferrous ion-dependent. We next used 4,4-difluoro-4-bora-3a,4a-diaza-s-indacene (BODIPY-C11), a sensor dye that monitors lipid peroxidation in ferroptotic cells ^56^, and found increased BODIPY oxidation/reduction ratio in senescent cells upon SA treatment (Supplementary Fig. 9b). However, this tendency was largely reversed by GSTP1, JNK or ferroptosis inhibitors, further substantiating the pivotal role of GSTP1/JNK/c-Jun axis in mediating ferroptosis of senescent cells.

It is reported that both apoptosis and ferroptosis are closely related to mitochondrial dysfunction, a subcellular change that can induce oxidative stress damage and in turn further promotes programmed cell death activities ^57, 58^. To profile mitochondrial dynamics, we employed transmission electron microscopy (TEM) for morphological characterization. Senescent cells treated by SA compounds showed ruptured mitochondria, abnormally shaped cristae, which are surrounded by an increased number of dark-stained fragments as compared with control cells, the latter generally exhibiting round-shaped mitochondria with well-developed cristae (Fig. 5h). Thus, TEM imaging provides ultramicro evidence for the subcellular structure-damaging effect of SAs in senescent cells, which were induced to undergo a dual death modality characterized by both apoptosis and ferroptosis.

### SAs promote tumor regression and restrain chemoresistance conferred by treatment-damaged TME

Since SAs have capacity and selectivity for eliminating senescent cells *in vitro*, we next queried whether these agents could be exploited to intervene in age-related pathologies *in vivo*. Drug resistance can affect the efficacy of anticancer treatments and senescent cells can contribute to therapeutic resistance through an *in vivo* SASP in the treatment-damaged tumor microenvironment (TME) ^21, 22, 59^. Targeted elimination of therapy-induced senescent cells reduces side-effects of chemotherapy and prevents cancer relapse in animals ^28, 60^. Despite these advances, the feasibility of SA-mediated clearance of senescent cells from primary tumors to increase efficacy of anticancer treatments needs further study.

To this end, we chose to generate tissue recombinants by combining PSC27 cells with PC3 cells, the latter a malignant prostate cancer (PCa) cell line, at a pre-optimized ratio under *in vitro* condition (1:4) ^22^. Admixed cells were subcutaneously implanted to the hind flank of mice with non-obese diabetes and severe combined immunodeficiency (M-NSG). The resulting tumors were measured at the end of an 8-week period, with xenograft tissues gathered for pathological assessment. Compared with tumors comprising naive PSC27 stromal cells and PC3 cancer cells, xenografts composed of senescent PSC27 cells and PC3 cells had increased volumes, confirming the tumor-promoting effects of senescent cells *in vivo* (Extended Data Fig. 6a).

We designed a preclinical protocol to incorporate genotoxic therapeutics and/or senolytics to imitate clinical scenarios (Fig. 6a). Upon stable establishment of cancer grafts in the transplanted animals, which usually takes 2 weeks after subcutaneous implantation, a single dose of placebo or mitoxantrone (MIT, a chemotherapeutic agent used in clinical oncology) was delivered to mice on the 1st day of the 3rd, 5th and 7th weeks until the end of the 8-week regimen (Extended Data Fig. 6b). In contrast to placebo group, administration of MIT delayed tumor growth, thus validating the efficacy of MIT as a chemotherapeutic drug (46.1% reduction in tumor size) (Fig. 6b). Although administration of SA compounds, as exemplified by the case of SAA intervention, did not cause tumor shrinkage, MIT treatment followed by SAA delivery (20 mg/kg *via* intraperitoneal (*i.p.*) injection 2 weeks post the 1st MIT dose and then delivered biweekly) promoted tumor regression (49.5% reduction of tumor volume compared with MIT treatment alone; 72.8% reduction compared with the placebo treatment) (Fig. 6b).

**Fig. 6.**
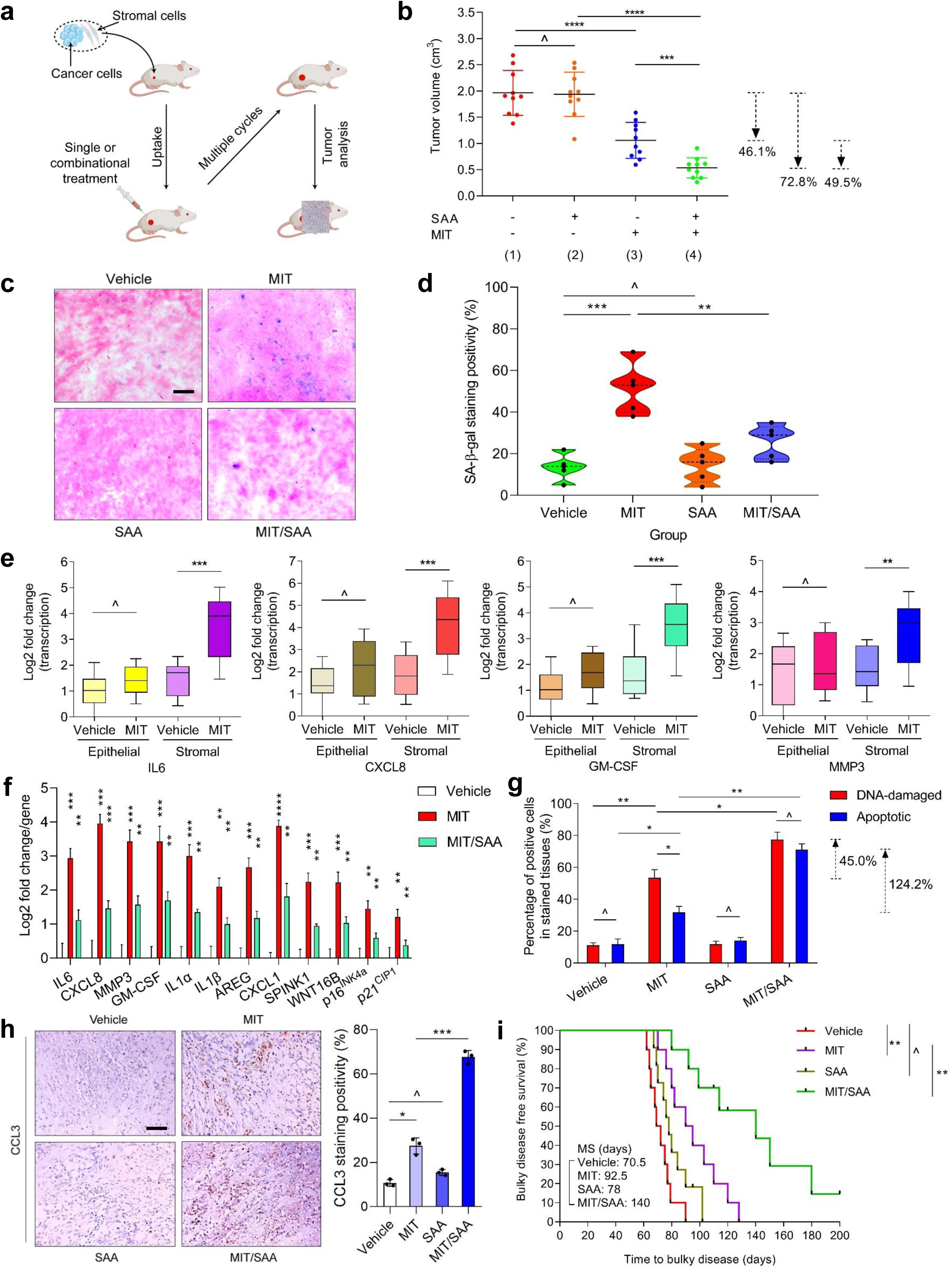
Senolysis by SA compounds reduces cancer resistance acquired from the treatment-damaged tumor microenvironment. (**a**) A schematic workflow illustrating the preclinical therapeutic procedure. Two weeks after subcutaneous inoculation and *in vivo* uptake of PC3/PSC27 recombinants, severe combined immunodeficient (M-NSG) mice were subject to either single agent or combinatorial treatment in a metronomic schedule of several cycles. The illustration was created in BioRender. (**b**) Comparative statistics of tumor end volumes. PC3 cancer cells were inoculated either alone or combined with PSC27 stromal cells before being implanted subcutaneously to the hind flank of M-NSG animals, which were administered with MIT and SAA, either alone or in combination. (**c**) Representative images of *in vivo* senescence in tumor foci by SA-β-gal staining. Scale bar, 50 μm. (**d**) Comparative statistics of tumor senescence as described in (**c**). (**e**) Transcriptional analysis of a subset of SASP factors expressed in epithelial *vs.* stromal cells acquired from tumor foci after laser capture microdissection (LCM) to isolate stromal and cancer cells, respectively. Signals were normalized to that of the sample with lowest value in the placebo group. (**f**) Transcriptional analysis of SASP factors and two canonical senescence biomarkers p16^INK4a^ and p21^CIP1^. (**g**) Statistical assessment of DDR and cellular apoptosis in tumors. Values are shown as the percentage of cells positively stained by IF or IHC specific to γH2AX or cleaved caspase 3 (CCL3), respectively. (**h**) Representative images of IHC staining for CCL3 at the completion of treatment regimens. Scale bar, 50 μm. (**i**) Comparative survival of animals sacrificed upon development of advanced bulky disease. Survival duration was calculated from tissue recombinant injection until death. *P* values were calculated by a two-sided log-rank (Mantel-Cox) test. DDR, DNA damage response. IF, immunofluorescence. IHC, immunohistochemistry. Data in **b**, **d**, **e**, **f** and **g** are shown as mean ± SD and representative of 3 independent biological replicates, with *P* values calculated by Student’s *t*-tests. ^, *P* > 0.05; *, *P* < 0.05; **, *P* < 0.01; ***, *P* < 0.001; ****, *P* < 0.0001.

We next tested if cellular senescence occurred in tumor foci. Unsurprisingly, MIT administration caused appearance of many senescent cells in tumor foci. Of note, SAA delivery to chemotherapy-treated animals eliminated the majority of senescent cells (Fig. 6c,d). We performed laser capture microdissection (LCM), a technique that allows selective acquirement of targeted cell lineages from tissues, which was then coupled to transcriptomics-based assays. The data indicated significantly enhanced expression of SASP factors such as IL6, CXCL8, GM-CSF, MMP3, IL1α and SPINK1, a change accompanied by upregulation of genes encoding the senescence markers p16^INK4a^ and p21^CIP1^ in chemotherapy-challenged animals (Fig. 6e and Extended Data Fig. 6c). Interestingly, this tendency was predominantly observed in stromal cells rather than their neighboring cancer cell counterparts, implying the possible repopulation of residual cancer cells, which can develop gain-of-functions, particularly drug resistance acquired from the treatment-damaged TME. Nevertheless, upon SAA administration to animals, the SASP-intensive expression pattern was attenuated, as evidenced by transcript assays and RNA-seq data (Fig. 6f and Extended Data Fig. 6d).

To delineate the mechanisms supporting SASP upregulation in MIT-treated mice, we dissected tumors from animals challenged with these two agents 7 d after the 1st dose of SAA delivery, a time point prior to the development of resistant colonies in tumor foci. Compared with placebo treatment, MIT administration caused DNA damage and apoptosis, whereas SAA treatment alone did not (Fig. 6g). However, when MIT-treated mice were co-administered SAA, the extent of DNA damage and apoptosis appeared to be increased, implying enhanced cytotoxicity in animals that received treatment by both chemotherapy and the senolytic agent. Of note, we observed increased caspase 3 cleavage, a hallmark of cellular apoptosis, when SAA was administered together with MIT (Fig. 6h). These data suggest there is an enhanced apoptotic effect by combining of SAA with a chemotherapeutic agent.

Since SAA appears to improve short term anticancer therapeutic outcomes, posttreatment survival of the different animal groups was investigated. Tumor growth was monitored in the preclinical cohort, by assessing if tumor burden exceeded a threshold (a size of 2,000 mm^3^), a criterion used by others ^21, 61^. Animals receiving the combined MIT/SAA treatment exhibited the most prolonged median survival, which was at least 38.9% longer than the group treated with MIT alone (Fig. 6i, green *vs* pink). However, SAA alone only marginally increased posttreatment survival. Thus, administration of SAA *per se* neither alters tumor growth nor promotes animal survival, whereas co-administration of SAA with MIT has a synergistic effect.

The therapies performed in these studies appeared to be well tolerated by mice, as no significant perturbations in creatinine, urea, liver and kidney enzyme levels or body weight were observed (Extended Data Fig. 6e,f). Furthermore, chemotherapeutic and geroprotective agents delivered at doses used in preclinical trials did not interfere significantly with immune system integrity or homoeostasis in critical organs, even in immunocompetent mice (Supplementary Fig. 10a-c).

We then tested if effects of SAA observed in these preclinical studies are reproduced by other SA compounds with senolytic activity. Results from *in vivo* treatments with SAB or SAE largely resembled those from SAA treatment (Supplementary Fig. 11a-h and Supplementary Fig. 12a-f). Together, these data support the possibility that the SA family members extracted from *S. miltiorrhiza*, particularly SAA, SAB and SAE, phytochemical molecules of geroprotective potential, once combined with classical chemotherapy, can enhance tumor response without causing severe toxicity *in vivo*.

### Preclinical intervention with SAs improves physical function and extends survival of prematurely aged mice

Senolysis represents an emerging strategy for treatment of various age-related disorders, while selective elimination of senescent cells alleviates pathological symptoms and improves organ function, as evidenced by preclinical studies and early clinical trials ^62^. To further test establish the effect of SA compounds on organismal aging, we first used model animals with accelerated development of senescence in tissues. We treated WT mice with whole body irradiation (WBI) at a sub-lethal dose (5 Gy), an approach that induces systemic cellular senescence ^28^. These therapy-challenged mice were subsequently subjected to geroprotective intervention with SA compounds (20 mg/kg *via* i.p.) or vehicle (twice *per* week) (Fig. 7a). Mice that had undergone WBI treatment manifested abnormal body appearance, including greying hair. However, this was alleviated by SAs (Fig. 7b and Supplementary Fig. 13a,b). SA-β-gal-positive cells were detected *in vivo* in these animals, with staining in cardiac, pulmonary, and prefrontal cortex tissues (Fig. 7c-f). Intermittent administration of SAA, SAB or SAE, decreased the percent of SA-β-gal-positive cells in these tissues significantly decreased, in contrast to control samples from vehicle-treated post-WBI mice (Fig. 7c-f and Supplementary Fig. 13c).

**Fig. 7.**
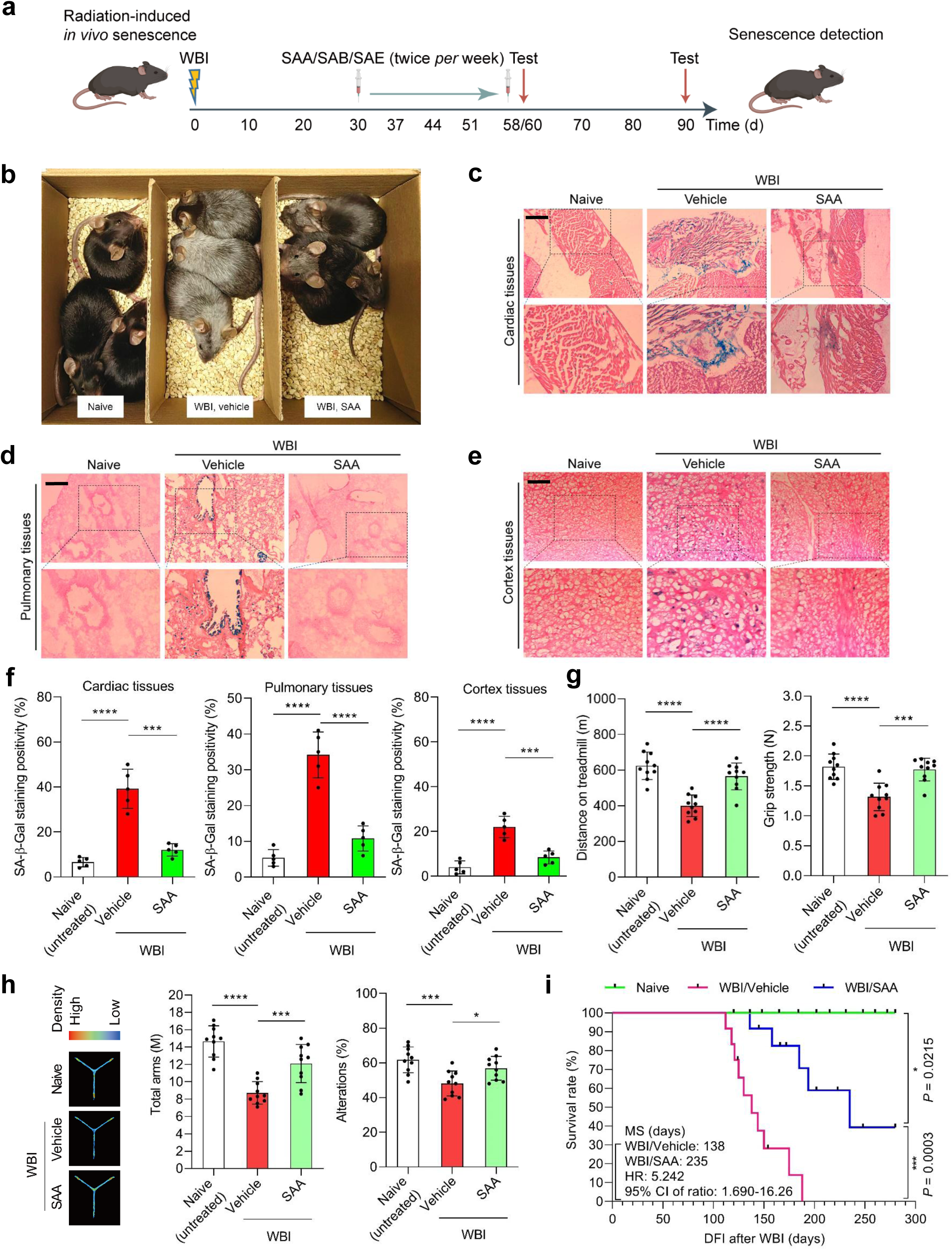
SAA treatment alleviates physical dysfunction in animals exposed to WBI to induce premature aging. (**a**) Schematic presentation of experimental procedure for mice experiencing whole body irradiation (WBI) and physical function tests. The illustration was created in BioRender. (**b**) Whole body snapshoot comparison of C57BL/6J males that were naïve, WBI-exposed followed by vehicle-treatment and WBI-exposed followed by PCC1-treatment, respectively. (**c**) Representative images of SA-β-gal staining of cardiac tissues of untreated (naive) and WBI-exposed mice subject to vehicle or PCC1 treatment. Scale bar, 200 μm. (**d**) Representative images of SA-β-gal staining of pulmonary tissues of mice as described in (**c**). Scale bar, 200 μm. (**e**) Representative images of SA-β-gal staining of cortical tissues of untreated (naive) and WBI-exposed mice subjected to vehicle or PCC1 treatment. Scale bar, 200 μm. (**f**) Comparative statistics of SA-β-gal staining positivity of cardiac (left), pulmonary (middle) and cortical (right) tissues of animals examined in (**c**), (**d**) and (**e**), respectively. (**g**) Measurement of running distance on treadmill (left) and grip strength (right) in experimental mice. (**h**) Mouse Y-maze test. Left, an illustrative model. Middle, comparative total arms. Right, comparative alterations. (**i**) Comparative survival of in naïve, WBI followed by treatment with either vehicle or SAA. *P* values were calculated by a two-sided log-rank (Mantel-Cox) test. Data in **f**-**h** are shown as mean ± SD and representative of 3 independent biological replicates, with *P* values calculated by Student’s *t*-tests. ^, *P* > 0.05; *, *P* < 0.05; **, *P* < 0.01; ***, *P* < 0.001; ****, *P* < 0.0001.

We assessed the impact of SA treatment on mouse physical function. Our data suggest that WBI *per se* compromised exercise capacity as well as muscle strength as determined by treadmill walking and grip strength assays in the vehicle group compared to naïve counterpart animals (Fig. 7g and Supplementary Fig. 13d). Of note, administering SAA, SAB, or SAE alleviated physical dysfunction.

In mammals, cognitive plasticity and short-term memory are executive functions mediated by the prefrontal cortex, which can become progressively impaired with aging ^63^. Although WBI caused a severe decline in cognitive function and memory, there was alleviation of short-term spatial working memory in the prematurely aged mice that had received SA based on Y-maze tests (Fig. 7h and Supplementary Fig. 13e). SA compounds, albeit administered in an intermittent schedule, prolonged posttreatment survival relative to the vehicle group, with animal body weight remaining largely unchanged (Fig. 7i and Supplementary Fig. 13f-h). These data suggest administering SA compounds can alleviate physical dysfunction and reduce cognitive impairment in the case of premature aging, without overtly affecting physiology.

### Senescent cell clearance by SA compounds alleviates physical dysfunction associated with natural aging

Recent studies demonstrated that even a small number of senescent cells can have effects on distant organs and transplanting senescent cells into young animals can induce physical dysfunction ^64, 65^. We queried whether SAs that selectively eliminate senescent cells *in vivo* can decrease physical dysfunction in chronologically aged animals. We treated 20-month-old WT animals with vehicle or SAA (20 mg/kg *i.p.*) once every 2 weeks for 4 months (Fig. 8a). There was a higher percent of SA-β-gal-positivity in lung, liver, kidney, prostate, heart, spleen, cortex, and pancreas of aged animals and this was decreased by SAA treatment (Fig. 8b,c, Supplementary Fig. 14a, Extended Data Fig. 7a-h and Extended Data Fig. 8a-f). Extent of senescent cell clearance by SAA approached that of PCC1, a flavonoid senolytic agent. SAA enhanced maximal walking speed, hanging endurance, treadmill endurance, grip strength, daily activity, and beam balance performance compared to animals treated with vehicle (Fig. 8d-i), although body weight and food intake changed little (Extended Data Fig. 8g,h). Effects observed with SAA were similar to SAB or SAE (Supplementary Fig. 14b-e).

**Fig. 8.**
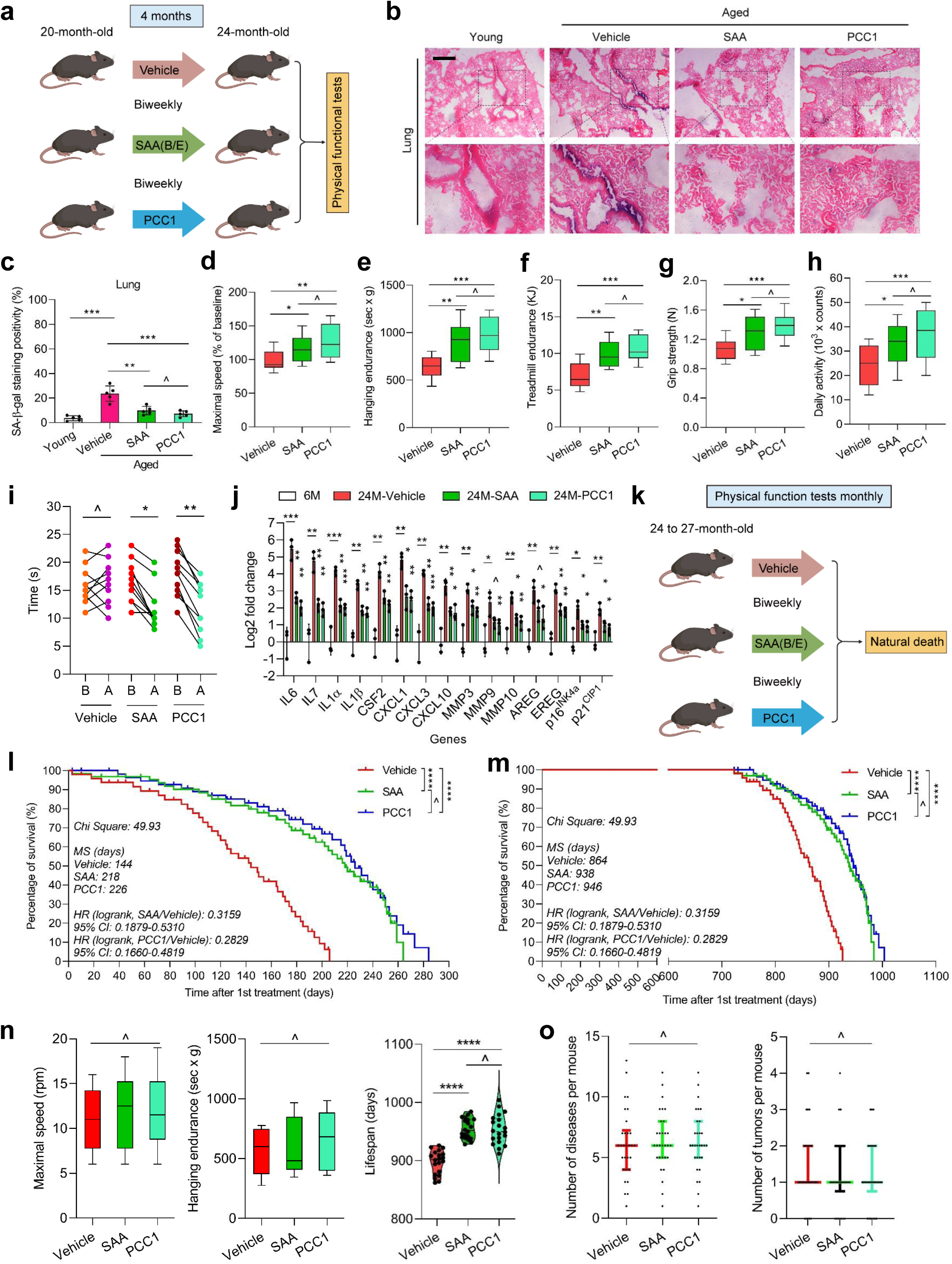
Intermittent SAA administration improves physical function and increases lifespan of naturally aged mice. (**a**) Schematic illustration of a preclinical procedure for 20-month-old C57BL/6J animals treated with SAA (SAB or SAE) once every 2 weeks (biweekly) for 4 months. After completion of the regimen, mice underwent physical function testing. PCC1 was administered as a natural senolytic control. The illustration was created in BioRender. (**b**) Representative images of SA-β-gal staining of lung tissues from young and aged mice treated with vehicle SAA. Scale bar, 100 μm. As a positive control for senolytics, PCC1 was administered in parallel. (**c**) Comparative statistics of SA-β-gal staining positivity in lung tissues of mice. (**d**-**h)** Quantification of maximal walking speed (relative to baseline) (**d**), hanging endurance (**e**), treadmill endurance (**f**), grip strength (**g**) and daily activity (**h**) of 20-month-old *C57BL/6J* males after a 4-month intervention with SAA. (**i**) Quantification of the time to cross a balance beam. Data before and after treatment of each animal were connected to allow direct comparison of treatments. (**j**) Quantitative transcript profiling of SASP expression in lung tissues collected from 6-month-old untreated (6M), 24-month-old vehicle-treated (24M-Vehicle) and 24-month-old SAA-treated mice (24M-SAA). The 24-month-old PCC1-treated mice (24M-PCC1) served as positive control. Data are shown as mean ± SD and derive from 3 biological replicates (*n* = 3 independent assays). (**k**) Schematic presentation of lifespan analysis of mice at 24-to-27 months of age. The illustration was created in BioRender. (**l**-**m**) Post-treatment survival (**l**) and whole-life survival (**m**) curves of *C57BL/6J* animals treated biweekly with SAA (*n* = 38; 20 males, 18 females) or vehicle (*n* = 39; 20 males, 19 females) starting at 24-27 months of age. In each case, PCC1 was administered as a positive control (*n* = 32; 16 males, 16 females). (**n**) Maximal walking speed and hanging endurance averaged over the last 2 months of life (*n* = 10/group), and lifespan for the longest living mice (top 20) in both groups. (**o**) Disease burden and tumor burden at death of mice. In the case of disease burden, for both sexes, *n* (animal number) = 30 *per* arm. For males (♂), *n* = 15 for V, *n* = 16 for SAA, *n* = 14 for PCC1. For females (♀), *n* = 15 for V, *n* = 14 for SAA, *n* = 16 for PCC1. (**p**) Schematic representation of key findings related with senolysis induced by SA compounds. Senescent cells exhibit inherent resistance to apoptosis and ferroptosis, a property that is mediated by the SCAP network involving bioactive molecules such as GSTP1. Mechanistically, SA compounds can target GSTP1 and enhance programmed death of senescent cells by activating a dual death modality characterized by both apoptotic and ferroptotic responses, potentiating their geroprotective role in counteracting natural aging and preventing age-related dysfunctions. SCAP, senescent cell anti-apoptotic pathway. For **c**-**h**, **j**, *n* = 3 biologically independent assays. Data in **d**-**h** are displayed as box-and-whisker plots, where a box extends from the 25th to 75th percentile with the median shown as a line in the middle, and whiskers indicating smallest and largest values (**d**-**h**, **n**), or as mean ± SD (**o**). Unpaired two-tailed Student’s *t*-tests (**c**-**j**, **n**, **o**) and Cox proportional hazard regression model (**l**, **m**). ^, *P* > 0.05; *, *P* < 0.05; **, *P* < 0.01; ***, *P* < 0.001; ****, *P* < 0.0001.

The SASP was attenuated in lungs of aged mice treated with SAA compared to vehicle-treated animals (Fig. 8j and Supplementary Fig. 14f). In lung transcriptome-wide expression profiles of animals treated with SAA, a number of intracellular factors were affected, many of which were SASP components (Supplementary Fig. 15a), generally in line with downregulated SASP factors in human stromal tissues after SAA intervention (Fig. 6f). In most of these studies, *in vivo* effects of SAA were comparable to those generated by PCC1, although effects of PCC1 were more pronounced. Effects of SAB or SAE were similar to those of SAA (Supplementary Fig. 14f). These results suggest that these small molecule compounds share geroprotective effects.

To further test the potential of senescent cell clearance to extend remaining lifespan, we treated with SAA from older ages (Fig. 8k). WT mice receiving SAA intervention (once *per* 2 weeks or biweekly) beginning at 24-27 months of age (approximately equivalent to an age of 75-90 years in humans) had a 51.4% longer median post-treatment lifespan (or 10.9% longer overall lifespan) and lower mortality hazard (68.4%, *P* < 0.0001) than the vehicle-treated group (Fig. 8l, m). These data suggest that SAA can reduce risk of age-associated mortality in old animals. We also treated aged animals with SAB or SAE. SAB did not fully recapitulate effects of SAA, but SAE approached the efficacy of SAA (Supplementary Fig. 15b-e). Hence, therapeutic potential differs among SA compounds administered *in vivo*.

We tested if reduced mortality in aged animals came at a cost of increased late-life morbidity. Physical function in mice treated with SAA or vehicle was tested monthly until death. Despite the longer remaining lifespan in SAA-treated mice, physical function during the last 2 months of life did not appear substantially lower than that in vehicle-treated mice (Fig. 8n and Extended Data Fig. 9 a,b). At autopsy, incidence of some age-related disorders, tumor burdens and cause of death differed little between SAA-treated and vehicle-treated animals (Fig. 8o). Similar findings were made in mice treated with SAB or SAE (Supplementary Fig. 15f,g). The SASP was attenuated in liver and lung, together with declines in peripheral blood protein levels of IL6, CXCL1 and CCL2 (Extended Data Fig. 9c-e). Oxidative stress appeared to be lower in liver of SAA-treated mice, with decreases in adducts of the lipid peroxidation product, 4-hydroxynonenal (HNE), and an increases in the ratio of reduced to oxidized glutathione (Extended Data Fig. 9f,g), consistent with the antioxidant properties of SA molecules, which scavenge free radicals and activate the antioxidant defense system ^66, 67^.

Altogether, we identified several senolytic members of the SA family, specifically SAA, SAB and SAE, a subset of hydrophilic phytochemicals mainly derived from *S. miltiorrhiza* Bunge. These compounds can reduce the burden of senescent cells and possibly other cells developing a pro-inflammatory phenotype and largely dependent on pro-survival senescence-associated anti-apoptotic pathways, with a notable advantage of extending lifespan but without causing increased morbidity in experimental mice. Through a series of preclinical trials, we demonstrated that even when administered in advanced life, therapeutic regimens involving SA compounds have the potential to significantly postpone age-related physical dysfunction, restrain age-related pathologies and improve health conditions. Therefore, our study provides a feasible and effective therapeutic paradigm to both improve health and increase lifespan by taking advantage of the senolytic potential of phenolic acids, an emerging subset of naturally derived senolytics.

## Discussion

Exploiting bioactive agents to target senescent cells with senotherapeutics, holds the potential to prolong healthspan and reduce the incidence and severity of chronic pathologies ^2^. Beneficial effects of dietary phytochemicals, agents mainly derived from fruits and vegetables, have generally been attributed to their anti-inflammatory and antioxidant properties ^68^. Given the promising efficacy of senolytics in helping achieve healthy aging with few side-effects, there is considerable interest in discovering novel senolytics, particularly those of natural origin. Naturally occurring senolytics may be less pharmacologically potent, but might cause lower toxicity than synthetic analogs. The first generation of senolytics, such as dasatinib, quercetin, fisetin and ABT-263, emerged in mid-2010s and were derived from targeted bioinformatics focusing on pathways that protect senescent cells against apoptosis ^26, 27, 69^. In contrast, several subsequently reported senolytics were identified through high-throughput screens and mechanistic studies. However, many of these senolytics have side-effects or intrinsic limitations to be translated into clinical settings. For instance, fisetin has limited bioavailability, while ABT-263 can cause thrombocytopenia and neutropenia in humans ^70^. To date, discovery of novel senolytic compounds particularly those of polyphenolic origin remains highly attractive.

Polyphenols comprise a superfamily of plant-derived secondary metabolites, usually characterized by the presence of more than one phenolic group in the molecule, and can exist as either monomers or polymers ^71^. Depending on their chemical structure, polyphenols can be classified as flavonoids, phenolic acids, stilbenes, lignans, curcuminoids, or tannins, several subgroups of phytochemicals that share common and key chemical features, partially explaining why they appear to generate similar favorable effects despite differences in chemical composition ^72, 73^. Flavonoids are broadly distributed across fruits and vegetables, and represent a major class of natural senolytic agents, including quercetin, curcumin, fisetin, luteolin and PCC1 ^71, 74^. The senolytic potential of other polyphenol subclasses has remained relatively underexplored. In this study, we found senolytic potential of several phenolic acid molecules, specifically SAA, SAB and SAE, which are mainly from the roots of *S. miltiorrhiza*, and validated the feasibility of using these small molecule compounds to target senescent cells. Administration with these phenolic acids appeared to effectively delay or alleviate aging-related conditions. Although a single biochemical process may not fully explain their pleiotropic pharmacological effects, we unraveled a functional mechanism that at least partially explains how SAs eliminate senescent cells (Fig. 8p). Our pre-clinical studies suggest translational potential of these senolytic SA compounds.

Ferroptosis is a form of programmed cell death that is different from apoptosis, pyroptosis and necrosis, and usually occurs in an iron-dependent manner with distinct morphologic features and molecular mechanisms ^75^. Loss of redox homeostasis causes intracellular accumulation of lipid peroxides, promoting ferroptosis, in which lipid peroxidation is a major regulator ^76^. Metabolic pathways triggering increased redox-reactive molecules including iron, enzymatic lipid peroxidation, ROS/reactive nitrogen species (RNS), and failure of endogenous antioxidant systems, mainly GPX4/glutathione (GSH) and coenzyme Q, can fuel ferroptosis ^77^. Linked to iron accumulation, ferroptosis elicits dysfunction of antioxidant systems, favoring lipid peroxide production and cell membrane damage, and ultimately, cell death. The signaling pathways eliciting ferroptosis are paradoxically associated with those protecting cells against iron excess and/or lipid-derived ROS. In this study, we found senescent cells have increased expression of GPX4, an antioxidant factor that inhibits ferroptosis, enabling them to have anti-ferroptotic potential. This is consistent with a recent study reporting that senescent cells rely on GPX4 to avoid ferroptosis and that GPX4-specific inhibitors kill senescent cells ^78^. Upon treatment with SAs, accelerated senescent cell death was observed, a process that was mediated partially by interfering with the GSTP1-GPX4 pathway to result in increased ferroptosis. In contrast to a recently reported prodrug nanoplatform, which was developed as a laser-free photodynamic therapy to degrade ferritin and trigger ferroptosis in senescent cells ^79^, SAs have the advantage of serving as naturally derived senolytic agents with relatively low cost, high safety and bioavailability.

Glutathione *S*-transferases (GSTs) catalyze conjugation of electrophilic compounds to endogenous nucleophilic glutathione substrates, a reaction that eliminates endogenous and exogenous toxic compounds, especially electrophiles ^80^. Although these enzymes generally interact with various kinases to modulate both cell proliferation and programmed cell death processes such as apoptosis, GSTP1 specifically catalyzes conjugation of electrophiles with glutathione during detoxification and plays an important role in regulating oxidative stress ^81^. Loss of GSTP1 can increase oxidative stress, alter mitogen-activated protein (MAP) kinase and NF-κB signaling, enhance caspase-3 cleavage and induce apoptosis ^82^. Genetic modulation of GSTP1-related axis or pharmacological inhibition of GSTP1’s catalytic activity sensitizes cancer cells to Food and Drug Administration (FDA)-approved ferroptosis-inducing agents ^42^. Here, we identified GSTP1 as a target of SAA, SAB and SAE, as it mediates apoptosis/ferroptosis-inducing effects of these SA compounds. Inhibitor-mediated GSTP1 targeting (such as by LAS17 or Ezatiostat) enhanced death of senescent cells, resembling the pattern observed upon exposure of these cells to SA compounds, implying GSTP1 plays an anti-apoptotic and anti-ferroptotic role in senescent cells and supports their long-term survival. Moreover, expression of GSTP1 can also inhibit cellular ferroptosis by decreasing acyl-CoA synthetase long chain family member 4 (ACSL4) activity and increasing GSH content ^83^. Consistent with this, senescent cells exhibited ferroptotic activity when GTSP1 was selectively targeted, suggesting anti-ferroptotic effects of this enzyme during cellular senescence that differs from its anti-apoptotic potential. These findings are in line with our data from SA-associated assays, indicating that SAs promote programmed death of senescent cells *via* induction of both apoptosis and ferroptosis.

Our RNA-seq and immunoblot data suggest that both ACSL4 and GPX4 are upregulated in senescent cells, allowing the formation of a pro-ferroptotic and anti-ferroptotic balance, at least on the ferroptosis arm. Disruption of this balance, for example, by SA molecules or GSTP1 inhibitors, can compromise cell survival potential *via* programmed cell death pathways including ferroptosis and apoptosis. Of note, we demonstrated the mechanism allowing SA compounds to induce ferroptosis in senescent cells, which is mediated by disengagement of the GSTP1/JNK/c-Jun pathway to downregulate GPX4 and disrupt the inherent anti-ferroptotic balance of senescent cells. Indeed, the interaction between GSTP1 and JNK/c-Jun can also be observed in the case of galangin-alleviated doxorubicin-induced cardiotoxicity, wherein JNK/c-Jun pathway was deactivated and ferroptosis was inhibited ^44^. GSTP1-mediated resistance to both apoptosis and ferroptosis of senescent cells is somehow reminiscent of p21, a master regulator that orchestrates a multi-faceted defense network to suppress ferroptosis and cuproptosis by governing redox and metal ion homeostasis, a case reported for resistant cancers ^84^. In senescent cells, engagement of both anti-ferroptotic and anti-apoptotic survival activities suggests a complex and intricate regulatory mechanism supporting the SCAP network, which merits further investigation across different cell lineages and various species including rodents and primates.

While our study provides valuable insights into the targets of SA compounds and sets a cornerstone for future research, it is essential to recognize the limitations of molecular modeling and the necessity for extended validation by medicinal chemistry, such as using computational chemistry methods that encompass quantum mechanics and molecular dynamics ^85^. SA compounds herein emerge as a subset of promising natural candidates for effective senolysis, and further investigation may unlock their full potential for improving health and enhancing longevity. Of note, most senolytics reported so far kill senescent cells by engaging the apoptotic pathway *via* interference with signal transduction in the SCAP network. Our study indicates that senescent cells have both anti-apoptotic and anti-ferroptotic defenses and point to new approaches for targeting senescent cells. A recent study defined an iron-triggered, ACSL4-governed and lipid peroxidation-driven program that arises with age across diverse cell types and multiple organs in non-human primates ^86^. Termed as ferro-aging, this process is triggered by iron (particularly ferrous) overload and lipid peroxidation, operates in a chronic and low-grade mode, mechanistically coupling iron dysregulation to progressive cellular senescence and organismal aging. It is noteworthy that ferro-aging distinguishes from acute ferroptosis, one of the major death modalities of senescent cells discussed in this study.

Agents with senolytic effects in murine models have advanced into early phase clinical trials for senescence-related pathologies. Among these are natural products and re-purposed drugs. The most comprehensively studied senolytics include “D + Q”, fisetin, piperlongumine, UBX0101, curcumin and analogues such as EF24 ^2, 11^. Early preclinical studies have yielded promising results, although much further work remains to be done to test the geroprotective potential of these senolytic agents ^15, 18^. Compared to vehicle-treated control, intermittent administration of senolytics can alleviate physical dysfunction of aged or diseased animals. Intermittent “D + Q” administration appears to have acceptable safety profiles in clinical trials for several senescence-related conditions ^87^. The preclinical SA regimens we tested appear to be feasible, safe, and effective and can improve overall health in 20-month-old and 24-27-month-old mice. SA compounds show promise in preventing cognitive dysfunction in prematurely aged animals. These preclinical data support moving towards phase I clinical trials to explore safety and feasibility SA compounds and potentially future clinical trials of target engagement, efficacy, and effectiveness for senescence-related disorders and diseases.

Although median lifespan is increasing globally, healthspan is not ^88^. Hence, geroprotective interventions that may close this disability gap by enhancing healthspan are needed. Research involving screening, design and development of senolytic drugs to selectively eliminate senescent cells is becoming a biomedical science priority. However, much work is needed to facilitate clinical use of senolytic agents and increase knowledge of their molecular targets and associated pathways. The current study provides proof-of-concept evidence to support targeting senescent cells with an emerging group of natural polyphenols, namely phenolic acids, to improve overall health in old age. It appears SA compounds exert senolytic activity by binding to GSTP1, altering its protein stability and compromising its antioxidant capacity. Despite potential challenges for establishing the safety and effectiveness of these agents in future clinical trials, SA compounds add to the reservoir of geroprotective drugs and may one day contribute to personalized solutions for extending healthspan of humans. We propose that SA compounds, which appear to be a novel category of natural compound senolytic agents, hold promise for future translational clinic trials, particularly those in the elderly to increase remaining healthspan.

## Methods

### Cell culture

The human prostate primary normal stromal cell line PSC27 was a gift from Dr. Peter Nelson (Fred Hutchinson Cancer Center) and maintained in stromal complete medium as described previously ^22^. The human fetal lung normal fibroblast line IMR90 and human primary normal umbilical vein endothelial cell line, HUVECs were from ATCC and cultured with EMEM medium supplemented with 10% FBS or vascular cell basal media supplemented with endothelial cell growth kit components with addition of human recombinant VEGF, respectively. The human adipose tissue-derived mesenchymal stem cell (AT-MSC) line was from ATCC and cultured in MSC basal medium (2% FBS) with 10 ng/ml bFGF and 10 μg/ml recombinant human insulin as reported ^89^. The human prostate epithelial cancer cell line, PC3, was from ATCC and cultured with F-12K media (10% FBS). All lines were tested for *Mycoplasma* contamination and authenticated with STR assays.

### Cell treatments

Stromal cells were grown until 80% confluent (CTRL) and treated with 50 μg/ml bleomycin (BLEO). Cells were rinsed briefly with PBS and maintained for 8-10 d prior to performing experiments. For drug screening of potential senolytics, natural candidates (totally 55 in the NMA library) (TargetMol anti-aging compound library L8200 TCM subset) were tested, each at 3 μg/ml for survival of 5.0 ☓ 10^3^ control and senescent cells for 3 d. Subsequent evaluation of effects of these candidate agents (each applied at 1 μg/ml) was conducted to test the extent of SASP suppression (as potential senomorphics). For agents showing promise for being effective and safe senolytics, further assays were performed. The phytochemical substances SAA, SAB and SAE were primarily examined over the range of 1 to 400 μM, with 100 μM, 100 μM and 50 μM determined to be the lowest effective concentrations for senolytic activity, respectively. The positive control senolytics ABT-263 (navitoclax) and PCC1 were applied at 1.25 μM and 50 μM, respectively. GSTP1 inhibitors LAS17 and Ezatiostat were used at 10 μM. Cell death suppressors QVD, VX-765, Nec-1s and LIP were applied at 10 μM, 20 nM, 20 nM and 200 nM, respectively. MG-132, chloroquine, bafilomycin, SP600125, JNK-IN-8, Ferrostatin-1, Liproxstatin-1 and LAS17 were used at 10 μM, 10 μM, 10 nM, 20 μM, 2 μM, 1 μM, 200 nM and 10 μM, respectively.

### Bulk RNA-seq and bioinformatics

Total RNA samples were prepared from senescent PSC27 cells cultured with regular media or media containing SA compounds (10 μM) for 3 consecutive days. Sample quality was validated by Bioanalyzer 2100 (Agilent) and RNA was subject to sequencing by Illumina NovaSeq 6000, with gene expression levels quantified by the software package RSEM (https://deweylab.github.io/RSEM/). Briefly, rRNAs in the RNA samples were eliminated using the RiboMinus Eukaryote kit (Qiagen), and strand-specific RNA-seq libraries were generated using the TruSeq Stranded Total RNA preparation kit (Illumina) according to the manufacturer’s instructions before deep sequencing.

Pair-end transcriptomic reads were mapped to the reference genome (GRCh38.p14) (Genome Reference Consortium Human Build 38; INSDC Assembly GCA_000001405.28, 12.2013) (http://asia.ensembl.org/Homo_sapiens/Info/Index) (ensembl_105) with reference annotation from GENCODE v42 using the Bowtie tool. Duplicate reads were identified using picard tools (1.98) script mark duplicates (https://github.com/broadinstitute/picard) and only non-duplicate reads were retained. Reference splice junctions are provided by a reference transcriptome (Ensembl build 73) ^90^. FPKM values were calculated using Cufflinks, with differential gene expression called by the Cuffdiff maximum-likelihood estimate function ^91^. Genes of significantly changed expression were defined by a false discovery rate (FDR)-corrected *P* value < 0.05. Only ensembl genes 73 of status “known” and biotype “coding” were used for downstream analysis.

Reads were trimmed using Trim Galore (v0.6.1) (http://www.bioinformatics.babraham.ac.uk/projects/trim_galore/) and quality assessed using FastQC (v0.10.0) (http://www.bioinformatics.bbsrc.ac.uk/projects/fastqc/). Differentially expressed genes (DEGs) were subsequently analyzed for enrichment of biological themes using the DAVID bioinformatics platform (https://david.ncifcrf.gov/) and the Ingenuity Pathways Analysis program (http://www.ingenuity.com/index.html). Raw data from bulk RNA-seq were deposited in the NCBI Gene Expression Omnibus (GEO) database under the accession codes GSE216218, GSE279345, GSE275945.

### Venn diagrams

Venn diagrams and associated empirical *P*-values were generated using the USeq (v7.1.2) tool IntersectLists ^92^. The *t*-value used was 22,008, as the total number of genes of status “known” and biotype “coding” in ensembl genes 73. The number of iterations used was 1,000.

### RNA-seq heatmaps

For each gene, the FPKM value was calculated based on aligned reads, using Cufflinks ^91^. Z-scores were generated from FPKMs. Hierarchical clustering was performed using the R package heatmap.2 and the distfun = “pearson” and hclustfun = “average”.

### Pulldown of SA compound-bound proteins

PSC27 cells were lysed in IP lysate buffer on ice with protease and phosphatase inhibitors, then centrifuged at 4°C for 10 min and the supernatant was collected and subjected to protein quantification by BCA method. The supernatant was divided into 4 equal parts, treated with biotin or biotin-SAA (100 μM) with unlabeled SAA in IP buffer overnight at 4°C (other SA compounds treated in a similar manner). The next day, 30 μl streptavidin-agarose beads (Sigma-Aldrich, USA) were added for binding for 2 h at 4°C. After 3 rinses with IP buffer, proteins in beads were eluted, separated by SDS-PAGE and stained with silver or directly subjected to sample separation for MS analysis. The resultant proteomic data of MS were deposited to the ProteomeXchange Consortium (http://proteomecentral.proteomexchange.org) through the iProX partner repository with a unique dataset identifier PXD056831.

### 20K HuProt™ human protein microarrays

The HuProt™ Human protein Microarray covers approximately 20,000 full-length human proteins ^93^. Proteins tagged with GST were expressed and purified using a eukaryotic expression system, with each protein repeated on the microarray. SA compounds were biotin-labeled and processed alongside the microarrays. The microarray images were analyzed using GenePix Pro v6.0 software, with signal intensities of the background, positive, and negative control spots quantified. In this experiment, D-biotin served as the positive reference at 635 nm, with BSA and buffer acting as negative controls. After normalization of the data, proteins meeting the Z-score threshold in both replicates were selected as those binding to SA compounds in microarray experiments.

### Cellular thermal shift assay (CETSA)

To assess whether SAA (or other SA compounds) physically targets GSTP1 protein, CETSA-immunoblot experiments were carried out. CETSA was used to determine the binding efficiency of a drug to its targeting protein. Briefly, PSC27 cells in culture were trypsinized and precipitated by centrifugation, lysed on ice with IP lysis buffer containing protease inhibitor cocktail and centrifuged at 14,000 × g at 4°C for 20 min. Each lysate was then equally aliquoted into several parts in Eppendorf tubes, incubated with SAA (or other SAs) at room temperature for 1 h. Samples were heated for 3 min at different temperatures (25, 30, 35, 40, 45, 50°C). The precipitated proteins were separated from the soluble fraction by centrifugation, and boiled at 95°C for 10 min. Samples were then analyzed by immunoblotting.

### Surface plasmon resonance (SPR) assays

The binding affinity of SAA (or other SA compounds) to recombinant human GSTP1 (rhGSTP1) was determined using Biacore T200 instrument equipped with CM5 sensor chips (GE Healthcare) ^94^. Briefly, rhGSTP1 protein (SinoBiological) was diluted in sodium acetate solution (pH 4.5) to a final concentration of 30 μg/ml. GSTP1 protein was immobilized on a CM5 sensor chip (GE Healthcare) by amine and allowed to couple with the final immobilization densities of 5,400 RU. Immobilized GSTP1 protein was then applied to capture SAA (or other SAs). The running buffer contained PBS-P (10 mM sodium phosphate, 150 mM NaCl, 0.05% surfactant P20, pH 7.3) and 5% DMSO. The pH was changed from 7.0 to 4.5 to adjust the binding affinity over several pH environments. Experiments were performed at 25°C, with a series of concentrations (25, 12.5, 6.25, 3.125, 1.56 and 0.78 μM) of SAA (concentrations of other SAs may vary) analyzed at a flow rate of 30 μl/min, with an injection time of 60 s and a dissociation time of 60 s in each binding cycle. A blank immobilization was performed on one channel surface of the chip to adjust the binding response. Association and dissociation constants were acquired using the Biacore T200 Evaluation software (v3.0 GE Healthcare). Data were exported to GraphPad Prism to plot the final curves.

### Nano-LC-MS/MS for proteomic analysis

Peptides were resolved in 0.1% formic acid, with peptide concentration measured with the BCA protein quantification kit. Samples were separated utilizing an analytic column (75 μm × 15 cm; 3 μm; 100Å) (C-18, Thermo Scientific) on a nanoflow HPLC Easy-nLC 1200 system (Thermo Scientific), following a 120 min LC gradient at 300 nl/min. Buffer A comprising 0.1% (v/v) formic acid in H_2_O and Buffer B comprising 0.1% (v/v) formic acid in 80% acetonitrile. The solution gradient: 1%–5% B in 5 min; 5%-26% B in 85 min; 26%-37% B in 20 min; 37%-100% B in 5 min; 90% B in 5 min. MS analyses were conducted on a Orbitrap Lumos mass spectrometer (Thermo Scientific). MS scan was performed in positive ion mode with spray voltage at 1,900 V and ion transfer tube temperature at 320℃. Xcalibur software was used to acquire profile spectrum data in a data-dependent acquisition pattern (DDA). The MS1 full scan was set at a resolution of 60,000 @ m/z 200, AGC target 4e5 and maximum IT 50 ms by Orbitrap mass analyzer (350-1600 m/z), followed by ‘top 15’ MS2 scans generated by HCD fragmentation at a resolution of 15,000 @ m/z 200, AGC target 5e4 and maximum IT 50 ms. Isolation window was set at 1.6 m/z. The normalized collision energy (NCE) was set at NCE 30%, and the dynamic exclusion time was 30 s. Precursors with charge 1, 7, 8 and > 8 were excluded for MS2 analysis.

All raw Xcalibur files acquired from MS runs were analyzed using the default settings of Proteome Discoverer 2.4 (Thermo Scientific) with minor modifications. Enzyme specification during the search was trypsin. Carbamidomethylation of cysteine was selected as a fixed modification, while oxidation of methionine and N-terminal acetylation were selected as variable modifications. Mass tolerances for precursor and fragment ions were set at 10 ppm and 0.5 Da respectively. The SEQUEST search algorithm was used to search MS/MS spectra against a composite database comprised of all the Swissprot *homo sapiens* database comprising 20,423 sequences (downloaded in August, 2023). Minimum cutoff for peptide length was set at 7 amino acids, and maximum permissible missed cleavage was set at 2. Maximal FDR for peptide spectral match, proteins and site was set to 0.01.

### Modelling of SAA/SAB/SAE binding to GSTP1 by molecular docking

The amino acid sequences of SAA/SAB/SAE (code: P09211) were acquired from UniProt. The X-ray diffraction microscopy structure of GSTP1 (PDB ID: 2A2R) was derived from RCSB Protein Data Bank (PDB, https://www.rcsb.org/). Water molecules farther than 4.5 Å from the receptor were removed. The structures of SAA/SAB/SAE were obtained from PubChem (https://pubchem.ncbi.nlm.nih.gov/), with the ligand-receptor interactions imitated by docking and analyzing with the ‘Triangle Matcher’ placement method with the molecular operating environment (MOE) software (2019.01.02) using the MMFF94 force field with the gradient convergence set to 0.1 kcal/mol. All protein structures were then prepared by the built-in MOE structure preparation and Protonate3D software tools using the default parameters illustrated with open source PyMOL software. The binding site of each protein was generated by the SiteFinder module in MOE. All conformations *per* ligand were scored using the ‘London dG’ scoring function, submitted to a refinement step based on molecular mechanics and rescored with the ‘GBVI/WSA dG’ scoring function, a force field-based scoring function that determines the binding free energy (kcal/mol) of the ligand from a given pose. Ultimately, the binding mode with the best docking score was selected for next step analysis. The docking score denotes the affinity between the protein receptor and the docking ligand. The lower docking score indicates the better affinity ^95^.

### Microscale thermophoresis (MST)

To substantiate the binding of SA compounds to wildtype and mutant GSTP1, we generated GSTP1 constructs encoding the wildtype protein and its mutant (Cys101Ser or Cys47Ser) tagged with a GFP reporter and overexpressed them in 293T cells. The cell sublines were lysed 48 h afterwards for MST analysis using Nano Temper Monolith NT Labelfree instrument (Nano Temper Technologies). A series of concentrations of SAs (800, 400, 200, 100, 50, 25, 12.5, 6.25,3.2, 1.6 and 0.8 μM) were mixed with GSTP1 protein in solution. After incubation at room temperature for 5 min, the mixtures were loaded into MO-AK002 capillaries. MST assessments were performed with 20% MST power and 20% LED power. MST datasets were processed using the NT analysis 1.5.41 software (Nano Temper).

### Immunoblot and immunofluorescence assays

Whole cell lysates were prepared using RIPA lysis buffer supplemented with protease/phosphatase inhibitor cocktail (Biomake). Nitrocellulose membranes were incubated overnight at 4°C with primary antibodies, and HRP-conjugated goat anti-mouse or -rabbit served as secondary antibodies (Vazyme). For immunofluorescence analysis, cells were fixed with 4% formaldehyde and permeabilized before incubation with primary and secondary antibodies, each for 1 h. After counterstaining with DAPI (0.5 μg/ml), samples were examined with an Imager A2. Axio (Zeiss) upright microscope to analyze specific gene expression.

### Intracellular reactive oxygen species (ROS) examination

Levels of intracellular ROS were determined using a ROS Assay Kit (Beyotime), which employs dichloro-dihydro-fluorescein diacetate (DCFH-DA) as a probe. Briefly, cells were cultured in 6-well plates for 24 h at 37°C and were then washed twice with serum-free medium. The medium containing 10 µM DCFH-DA was added. Cells were then incubated for 20 min at 37°C, with light avoided during incubation. After incubation, the cells were washed thrice with serum-free medium, then observed and photographed using a fluorescence microscope (Nikon). The fluorescence intensity was measured using ImageJ (v1.51, NIH).

### Transmission electron microscopy (TEM)

Stromal cells were subject to various treatments for 24 h before precipitation was collected in 2 ml centrifuge tubes after trypsinization. The precipitation was incubated with TEM fixative at 4 °C for 2 h and suspended to be wrapped in the heated 1 % agarose before solidification, with re-suspension and washing in 0.1 M phosphate buffer (PB) repeated for 2 more times (3 min each). Cells were post-fixed with 1 % OsO_4_ in 0.1 M PB at room temperature for 2 h. Afterward, cells were sequentially dehydrated in a graded ethanol series (30 %, 50 %, 70 %, 80 %, 95 %, 100 %, 100 %) for 20 min at each solution and two changes of acetone (15 min each). The cells were then penetrated and embedded with the sequential infiltration of the following formula: acetone: EMBed 812 = 1:1 for 2-4 h at 37 °C; acetone: EMBed 812 = 1:2 overnight at 37 °C; pure EMBed 812 for 5-8h at 37 °C. The cells were inserted into the pure EMBed812 in the embedding models and kept in 37 °C overnight before the models were moved into 65 °C to polymerize for over 48 h. Ultrathin sections (60-80 nm) were obtained on the ultramicrotome, and fished out onto the 150 meshes cuprum grids with formvar film. The slices were post-stained with 2 % uranium acetate saturated alcohol solution and 2.6 % lead citrate avoiding CO_2_ before dried overnight at room temperature. The cuprum grids were observed under TEM, with images captured on the spot.

### Lipid peroxidation activity assay

The BODIPY 581/591 C11 kit (MedChemExpress) was used to detect the lipid peroxidation activity of cell samples. First, the cell samples were digested with trypsin to make a single-cell suspension. After centrifugation, the supernatant was discarded, washed twice with PBS, and then 1 ml of BODIPY 581/591 C11 (lipid peroxidation sensor) working solution was added and incubated at room temperature for 30 min. The supernatant was then centrifuged and washed twice with PBS. Cells were resuspended in serum-free medium and analyzed by fluorescent microscopy.

### GPX4 promoter activity assay with reporter system

To clone the enhancer/promoter region of human GPX4, a 1000-bp non-encoding region immediately upstream of the transcription start site of human GPX4 gene was amplified by PCR and inserted into the multiple cloning site of pGL4.22[luc2CP Puro] firefly luciferase reporter vector (Promega, USA) between NdeI and HindIII restriction sites. The resulting construct was used to transfect PSC27 cells, together with pGL4.22-Rluc-Neo vector (Promega, USA) for signal normalization. To apply the Dual-Luciferase® Reporter Assay System (DLR) (Promega, USA), a 1:1:8 ratio of either Rluc (Renilla): Fluc (firefly):construct DNA was followed. SAA, SAB or SAE was added to cell culture, and cells were collected for lysis and luciferase signal analysis 12 h after.

### Experimental mice and chemotherapeutic studies

Animals were maintained in a specific pathogen-free (SPF) facility, with M-NSG (NOD-PrkdcscidIl2rgem1/Smoc) (Shanghai Model Organisms) mice at an age of approximately 6 weeks (∼20 g body weight) being used. Ten mice were incorporated into each group, with xenografts subcutaneously generated at the hind flank under isoflurane inhalation. Stromal cells (PSC27) were mixed with cancer cells (PC3) at a ratio of 1:4 (i.e., 250,000 stromal cells admixed with 1,000,000 cancer cells to make tissue recombinants before implantation *in vivo*). Animals were sacrificed at 2-8 weeks after implantation, according to tumor burden and/or experimental requirements. Tumor growth was monitored weekly, with tumor volume (v) measured and calculated according to tumor length (l), width (w) and height (h) by the formula: v = (π/6) × ((l+w+h)/3)^3 96^. Freshly dissected tumors were either snap-frozen or fixed to prepare FFPE samples. Resulting sections were used for IHC staining against specific antigens or subjected to hematoxylin/eosin staining.

For chemoresistance studies, animals received subcutaneous implantation of tissue recombinants as described above and were given standard laboratory diets for 2 weeks to allow tumor uptake and growth initiation. Starting from the 3^rd^ week (tumors reaching 4-8 mm in diameter), MIT (0.2 mg/kg doses), the senolytic agent SAA (20.0 mg/kg doses, 200 μl/dose) (or other SA compounds) or vehicle controls were administered by intraperitoneal injection (therapeutic agents *via* i.p.), on the 1^st^ day of 3^rd^, 5^th^ and 7^th^ week, respectively. Upon completion of the 8-week therapeutic regimen, animals were sacrificed, with tumor volumes recorded and tissues processed for histological evaluation.

At the end of chemotherapy and/or targeting treatment, animals were anaesthetized and peripheral blood was gathered *via* cardiac puncture. Blood was transferred into a 1.5 ml Eppendorf tube and kept on ice for 45 min, followed by centrifugation at 9000 x g for 10 min at 4°C. Clear supernatants containing serum were collected and transferred into a sterile 1.5 ml Eppendorf tube. All serum markers were measured using dry-slide technology on IDEXX VetTest 8008 chemistry analyzer (IDEXX). About 50 μl of the serum sample were loaded on the VetTest pipette tip before being securely fitted on the pipettor and manufacturer’s instructions were followed for further examination.

### Whole body irradiation of wildtype animals

Wildtype experimental animals were housed in a SPF facility at an ambient temperature of 22-25°C with 30% humidity, under a 12 h light/12 h dark cycle (6 am-6 pm), with free access to food and were fed ad libitum. For induction of *in vivo* senescence, *C57BL/6J* mice (males, Shanghai Lingchang Biotechnology) at 8-10 weeks of age were exposed to a sublethal dose (5 Gy) of whole body irradiation (WBI). Four weeks later, animals were injected with 20.0 mg/kg SAA (or other SA compounds) or vehicle *via i.p.* twice *per* week for eight consecutive weeks. Mice that had undergone WBI typically manifested an abnormal body appearance including markedly greyed hair, but this was partly reversed by SA compound administration.

### Y-maze test

Spontaneous alternation behavior indicates the tendency for mice to alternate their (conventionally) non-reinforced choices on successive iterations ^97^. The Y-maze spontaneous alternation was employed to assess short-time spatial recognition and working memory of mice by measurement of spontaneous alternations ^98^. The maze consisted of 3 arms positioned at a 120° angle from each other, with each arm 31 cm long, 5 cm wide or 10 cm high, respectively. Each arm had markers of different colors as distinct visual cues. After placed individually at the center of the apparatus, mice were allowed to explore freely through the maze during an 8-min session. Alternation was defined as successive entries into all 3 arms on overlapping triplet sets. The number of arm entries and alternations were recorded visually to calculate the percentage of the alternation behavior with the following formula: % Alternation = [Number of alternations/ (Total number of arm entries - 2)] x 100. Spontaneous alternation (%), defined as successive entries into 3 arms on overlapping triplet sets, is associated with short-term spatial learning memory.

### Senescent cell clearance and animal lifespan extension

For preclinical trials involving naturally aged animals, wildtype *C57BL/6J* mice (including both males and females, with sex generally not considered in study design) were maintained in a specific SPF facility at 22-25°C and 30% humidity under 12 h light/12 h dark cycle (6 am-6 pm). Mice were allowed to have free access to a standard rodent chow diet (5LOD, PicoLab), with water provided *ad libitum*. The experimental procedure was approved by the IACUC at Shanghai Institute of Nutritional and Health, Chinese Academy of Sciences or Binzhou Medical University, with all animals conducted in accordance with the guidelines for animal experiments defined by the relevant IACUC.

To determine the impact of SA compound administration on animal lifespan, 20-month-old naturally aging wildtype *C57BL/6J* mice were employed, which were generally sorted according to their body weight and randomly assigned to vehicle, a SA compound or PCC1 intervention groups. Animals were treated once every 2 weeks, in an intermittent manner for 4 months before being subjected to physical tests at 24 months of age. For preclinical trials involving lifespan extension at advanced stage, we used animals starting at 24-27 months of age (equivalent to human age of 75-90 years). Mice (both sexes) were treated once every 2 weeks (biweekly) with vehicle, each SA compound (SAA, SAB or SAE) or PCC1 by *i.p.* injection (20.0 mg/kg doses for each SA compound or PCC1, respectively) for 3 consecutive days. Some mice were moved from their original cages during the study to minimize single cage-housing stress. RotaRod (TSE system) and hanging tests were performed for monthly measurement of maximal speed and hanging endurance, respectively, as these tests are considered sensitive and noninvasive. Mice were euthanized and scored as having died if displaying more than one of the following signs: (i) incapable of drinking or eating; (ii) reluctant to move even upon exterior stimulus; (iii) rapid gross weight loss; (iv) severe balance disorder; or (v) bleeding or ulcerated skin. No animals was lost due to fighting, accidental death or dermatitis. The Cox proportional hazard model was used for survival appraisal.

All animal experiments were performed in compliance with NIH Guide for the Care and Use of Laboratory Animals (National Academies Press, 2011) and the ARRIVE guidelines, and were approved by the Institutional Animal Care and Use Committee (IACUC) of Shanghai Institute of Nutrition and Health, Chinese Academy of Sciences and Binzhou Medical University.

### Tissue SA-β-gal staining and histological examination

For SA-β-gal staining, frozen sections were dried at 37°C for 20-30 min before fixed for 15 min at room temperature. The frozen sections were washed 3 times with PBS and incubated with SA-β-Gal staining reagent (Beyotime) overnight at 37°C. After completion of SA-β-gal staining, sections were stained with eosin for 1-2 min, rinsed under running water for 1 min, differentiated in 1% acid alcohol for 10-20 s, and washed again under running water for 1 min. Sections were dehydrated in increasing concentrations of alcohol and cleared in xylene. After drying, samples were examined under a bright-field microscope.

### Histology and immunohistochemistry

Preclinical specimens from mouse tissues were fixed overnight in 10% neutral-buffered formalin and processed for paraffin embedding. For histological evaluation, standard staining with hematoxylin/eosin was performed on sections of 6-8 μm thickness cut from each specimen block. For immunohistochemistry, tumor specimens were first fixed in 4% paraformaldehyde and embedded in paraffin. For appraisal, tissue sections were cut from FFPE chunks, deparaffinized and incubated in 10 mM sodium citrate buffer (pH 6.5) at 95°C for 40 min for antigen retrieval. Peroxidase activity was quenched with 3% H_2_O_2_ and tissues were blocked in 5% bovine serum albumin for 30 min, before incubated with primary antibodies (*e.g*., against IL6 or cleaved caspase 3 [CCL3]), for SASP factor analysis or cell apoptosis evaluation, respectively) overnight at 4°C. After 3 washes with PBS, tissue sections were incubated with biotinylated secondary antibody (1:200 dilution, Vector Laboratories) for 1 h at room temperature then washed 3 times, after which streptavidin-horseradish peroxidase conjugates (Vector Laboratories) were added and the slides incubated for 45 min. DAB solution was then added and slides counterstained with haematoxylin. Alternatively, frozen tissue sections were used for immunofluorescence staining to probe target protein expression in animals receiving the various preclinical treatments, with counterstaining performed with DAPI. Images were then captured with an epifluorescence microscope.

### Appraisal of *in vivo* cytotoxicity by blood tests

For routine blood examination, 100 μl fresh blood was acquired from each animal and mixed with EDTA immediately. The blood samples were analyzed with Celltac Alpha MEK-6400 series hematology analyzers (Nihon Kohden). For serum biochemical analyses, blood samples were collected and clotted for 2 h at room temperature or overnight at 4 °C. An aliquot of approximately 50 μl serum was subject to analysis for creatinine, urea, alkaline phosphatase (ALP) and alanine transaminase (ALT) by an VetTest 8008 chemistry analyzer (IDEXX) as reported previously ^96^. For blood cell tests, 50 μl fresh blood was collected from each mouse and mixed with EDTA immediately. Circulating levels of hemoglobin, white blood cells, lymphocytes and platelets of the blood samples were analyzed by Sysmex XN-1000 series automated hematology analyzers (Sysmex, XN-1000).

All animal experiments were conducted in compliance with the NIH Guide for the Care and Use of Laboratory Animals (National Academies Press, 2011) and the ARRIVE guidelines, and were approved by the IACUC of Shanghai Institute of Nutrition and Health, Chinese Academy of Sciences. For each preclinical regimen, animals were monitored for conditions including hypersensitivity (changes in body temperature, altered breathing and ruffled fur), body weight, mortality and changes in behavior (i.e., loss of appetite and distress), and were disposed of appropriately according to the individual pathological severity as defined by relevant guidelines.

### Statistics and reproducibility

All *in vitro* experiments were performed in triplicate, while animal studies were conducted with at least 8 mice *per* group for most preclinical assays. Data are presented as mean ± SD except where otherwise indicated. GraphPad Prism (9.5.1) was used to collect and analyze data, with statistical significance determined according to individual settings. Cox proportional hazards regression model and multivariate Cox proportional hazards model analyses were performed with SPSS statistical software. Statistical significance was determined by unpaired two-tailed Student’s *t*-tests, one- or two-way ANOVA, Pearson’s correlation coefficients tests, Kruskal-Wallis, log-rank tests, Wilcoxon-Mann-Whitney tests or Fisher’s exact tests. For all statistical tests, a *P* value < 0.05 was considered significant. The sample size for each experiment was not predetermined by statistical method, and no data were excluded from analyses.

## Reporting summary

Further information on research design is available in the Nature Portfolio Reporting Summary linked to this article.

## Data availability

Experimental data supporting the plots within this article and other findings from this study are available from the corresponding author upon reasonable request. The RNA-seq data generated in the present study have been deposited in the Gene Expression Omnibus database under accession code GSE216218, GSE279345, GSE275945. The mass spectrometry proteomics data have been deposited to the ProteomeXchange Consortium *via* the PRIDE partner repository iProX ^99^ with the dataset identifier PXD056831. All data that support the findings of this study are available within the article and Supplementary Information. Source data are provided with this paper. A reporting summary for this article is available in the Supplementary Information.

## Acknowledgements

We are grateful to the members of Sun laboratory for reagents, comments and other contributions. This study was supported by grants from the Strategic Priority Research Program of Chinese Academy of Sciences (XDB39010500 to Y.S.); National Key Research and Development Program of China (2020YFC2002800 to Y.S.), National Natural Science Foundation of China (NSFC) (82130045, 82350710221 and 82571777 to Y.S.); Shanghai Municipal Science and Technology Commission Excellent Academic Leader Program (20XD1404300 to Y.S.); Taishan Scholar Project of Shandong Province (tstp20230634 Y.S.); Natural Science Foundation of Shandong Province Joint Fund Program (ZR202108130049, ZR2021LSW021 to Y.S.); the Connor Fund, Robert and Theresa Ryan, and the Noaber Foundation (to J.L.K.); Key R&D Program of Shandong Province (2023CXPT012 to G.Z.); the National Natural Science Foundation of China (22007006 to G.Z.); Taishan Scholar Project of Shandong Province (tsqn201909144 to G.Z.); Science and Technology Commission of Shanghai Municipality (22MC1940300 to Q.L.); Natural Science Foundation of Shandong Province (ZR2023MH262 to Q.F.) and Translational Medicine Association of Shandong Province (ZHA2025021 to Q.F.).

## Author contributions

Y.S. conceived this study, designed the experiments and orchestrated the project. Z.J. and H.Z. performed most of the *in vitro* assays and some of the *in vivo* experiments and wrote part of the manuscript. Q.X. and W.H. helped with cell culture, drug treatment and sample preparation. S.Z. assisted in cell culture and maintenance. H.Z. conducted proteomic profiling of human senescent cells. Q.F. assayed SASP expression *in vivo* and analyzed serum of animals. Q.L., G.Z. and J.L.K. provided constructive advice and/or supervised a specific subset of experiments. Y.S. performed data analysis, graphical presentation and finalized the manuscript. All authors critically read and commented on the final manuscript.

## Additional information

**Extended data** is available for this paper at designated link.

**Supplementary information.** The online version contains supplementary material available at the designated link.

**Correspondence and requests for materials** should be addressed to Yu Sun.

