## Supplementary Information for "Salvianolic acids are natural senolytics and increase lifespan in old age"

Supplementary Figures 1-15

Supplementary Tables 1-6

**Supplementary Fig. 1**

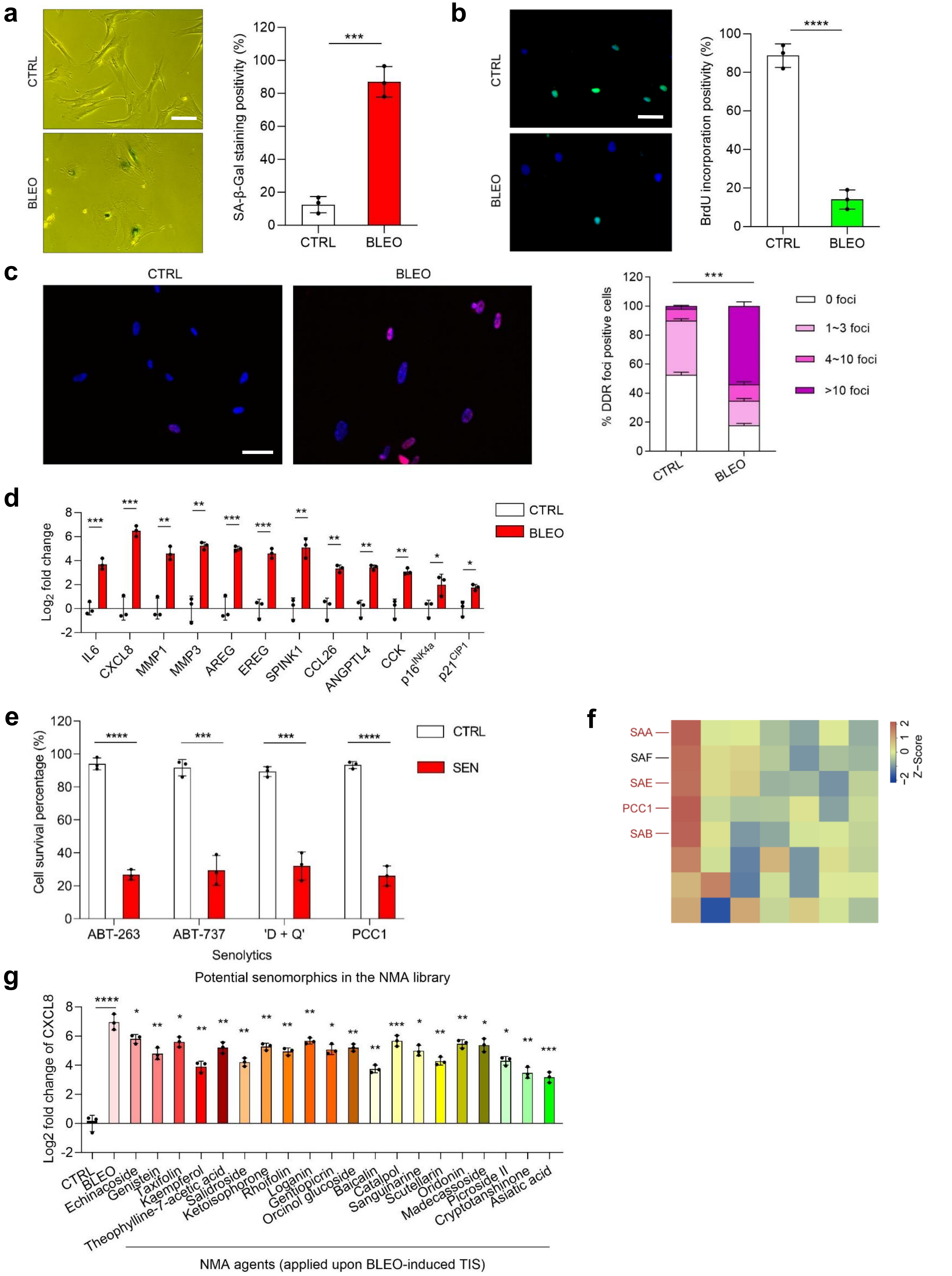

**Supplementary Fig. 2**

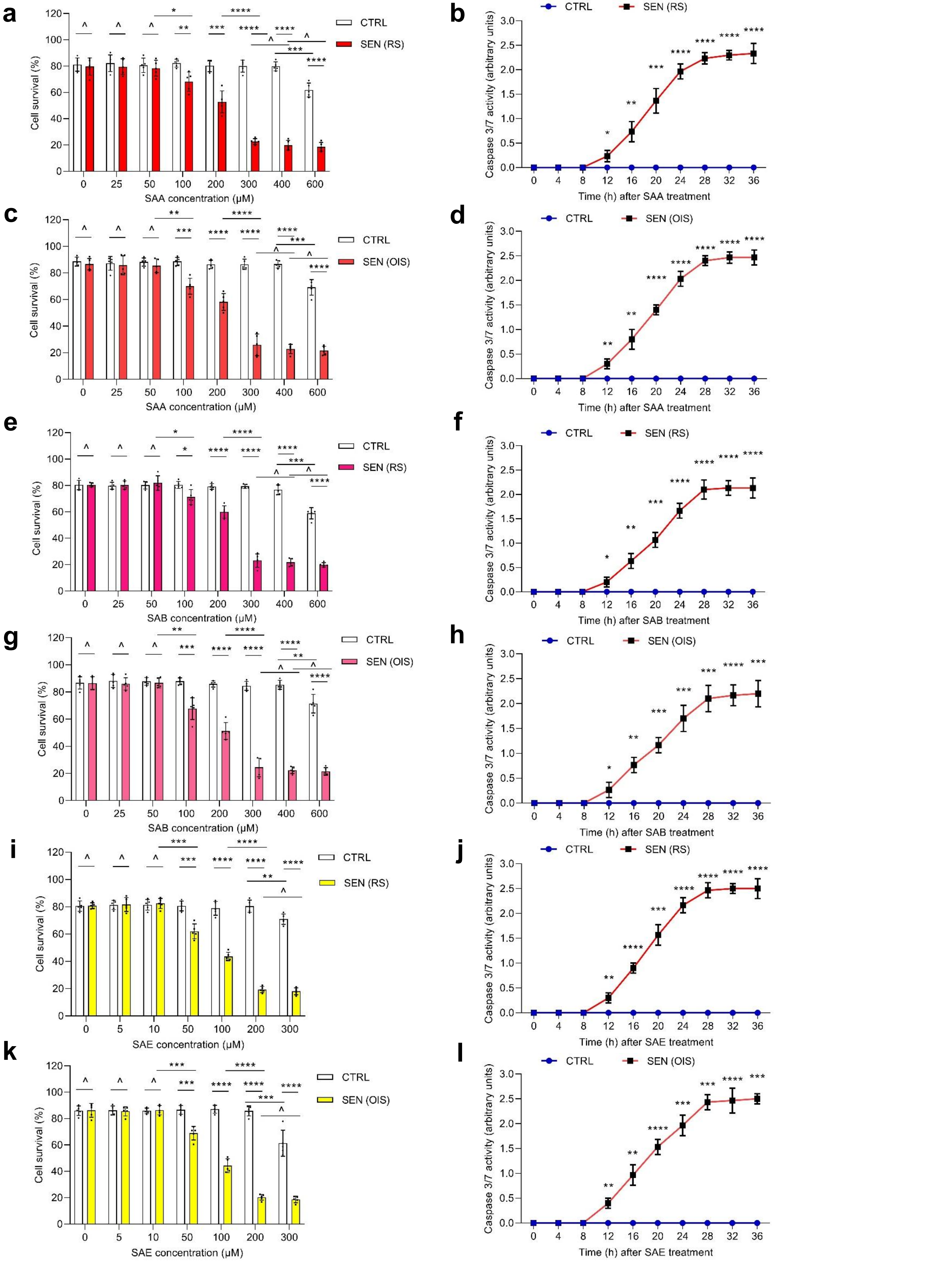

**Supplementary Fig. 3**

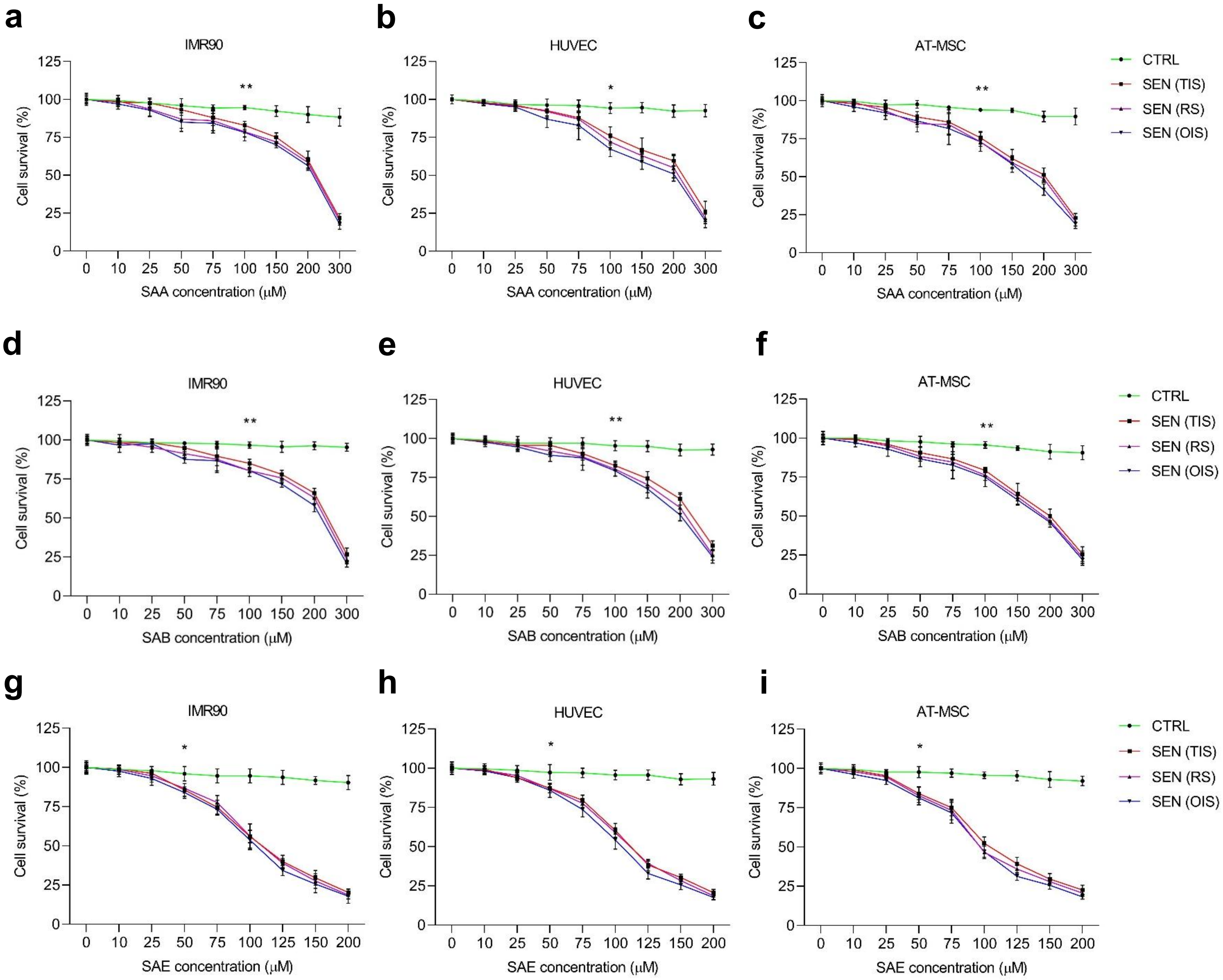

**Supplementary Fig. 4**

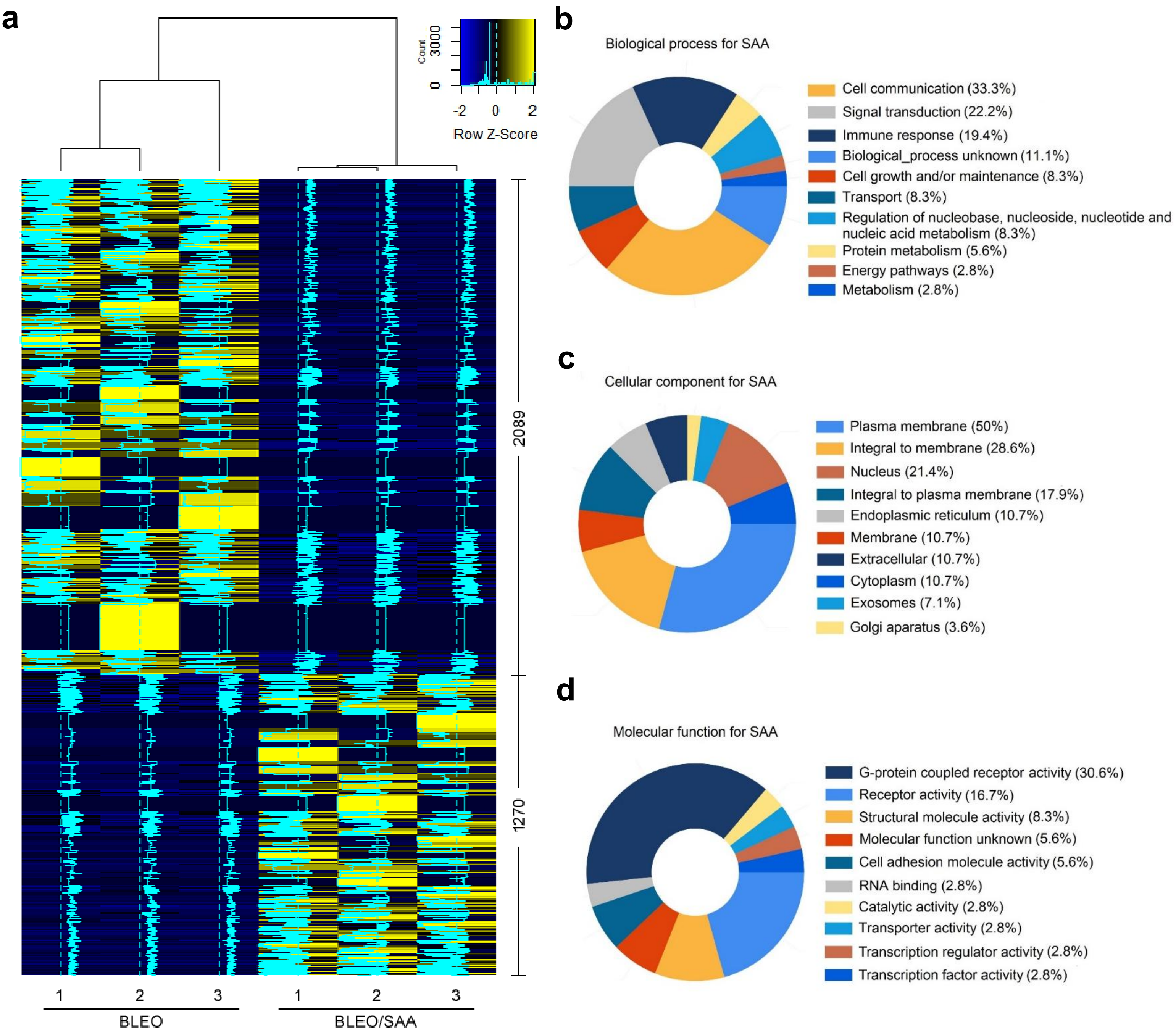

#### Supplementary Fig. 5

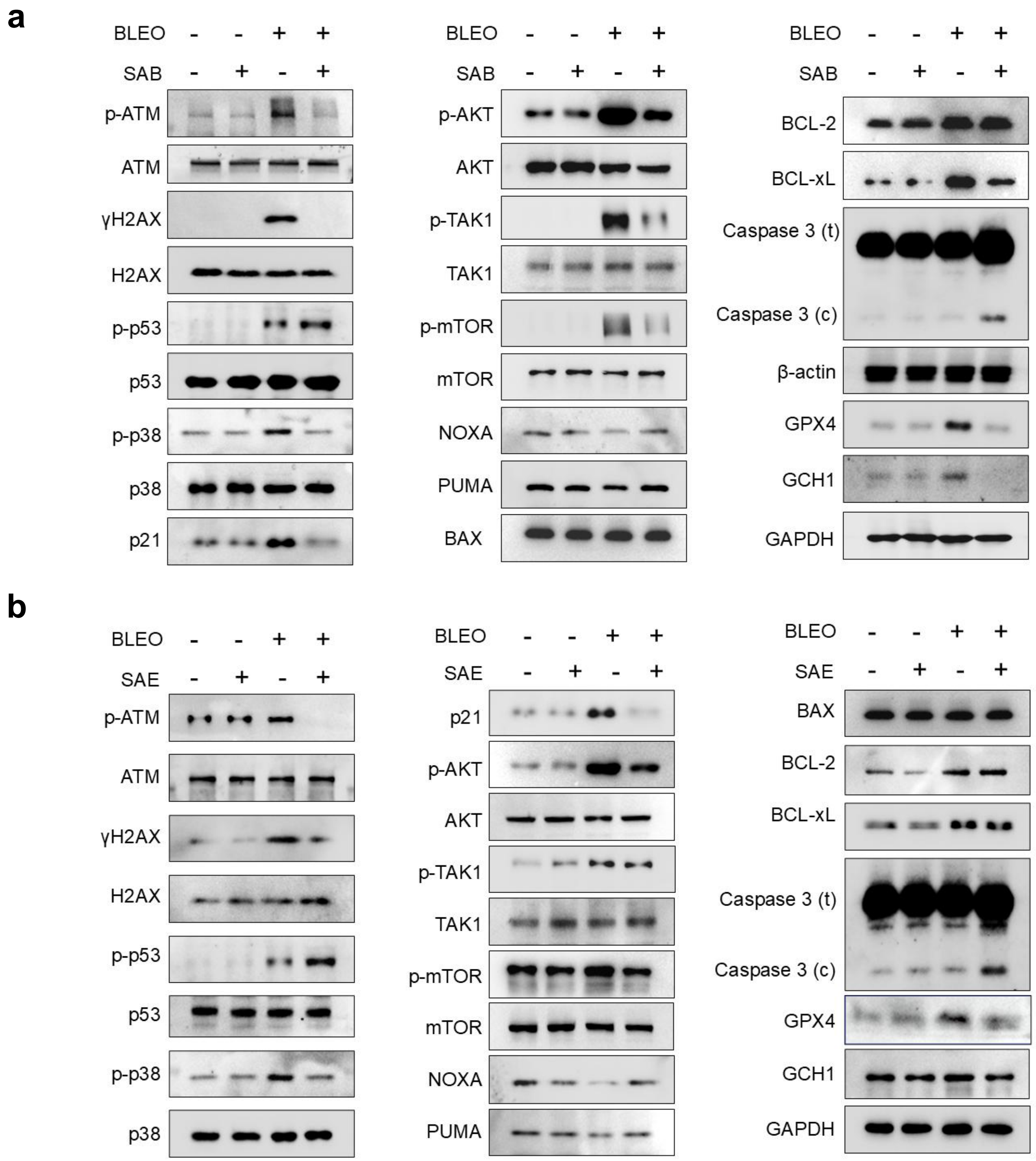

**Supplementary Fig. 6**

**a**

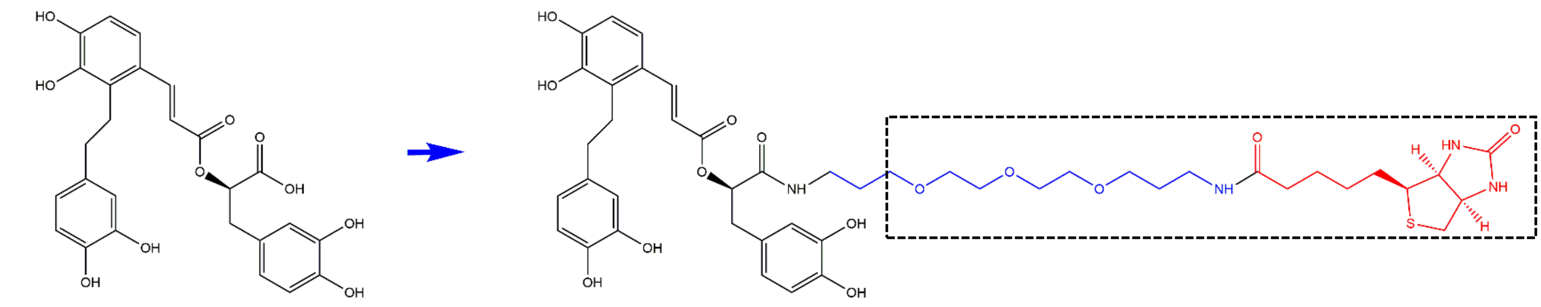

SAA (CAS: 96574-01-5)

SAA-Bio

**b**

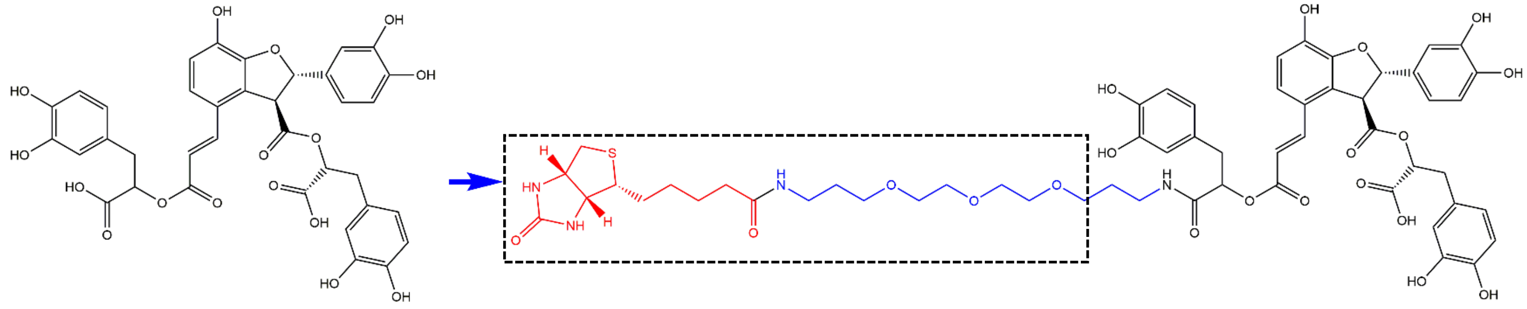

SAB (CAS:121521-90-2)

SAB-Bio

**c**

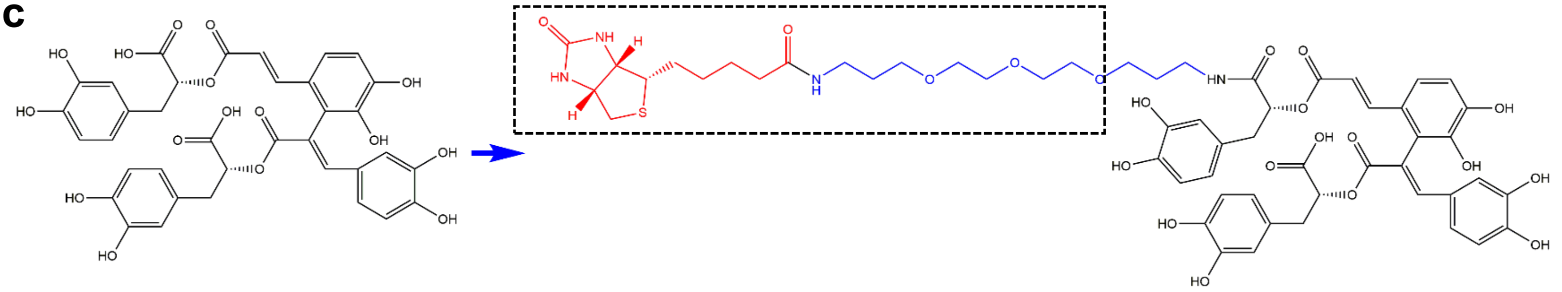

SAE (CAS:142998-46-7)

SAE-Bio

**Supplementary Fig. 7**

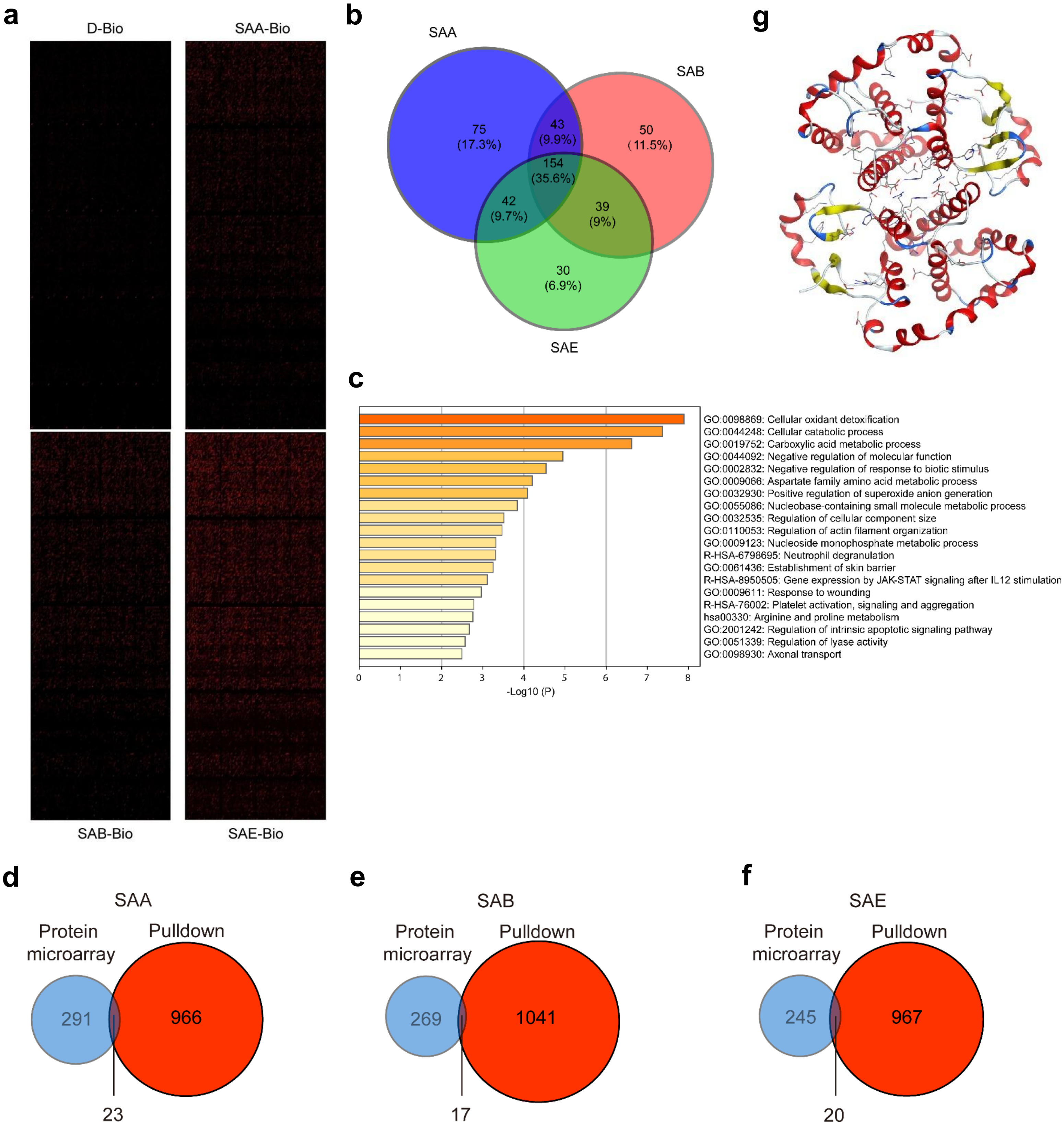

#### Supplementary Fig. 8

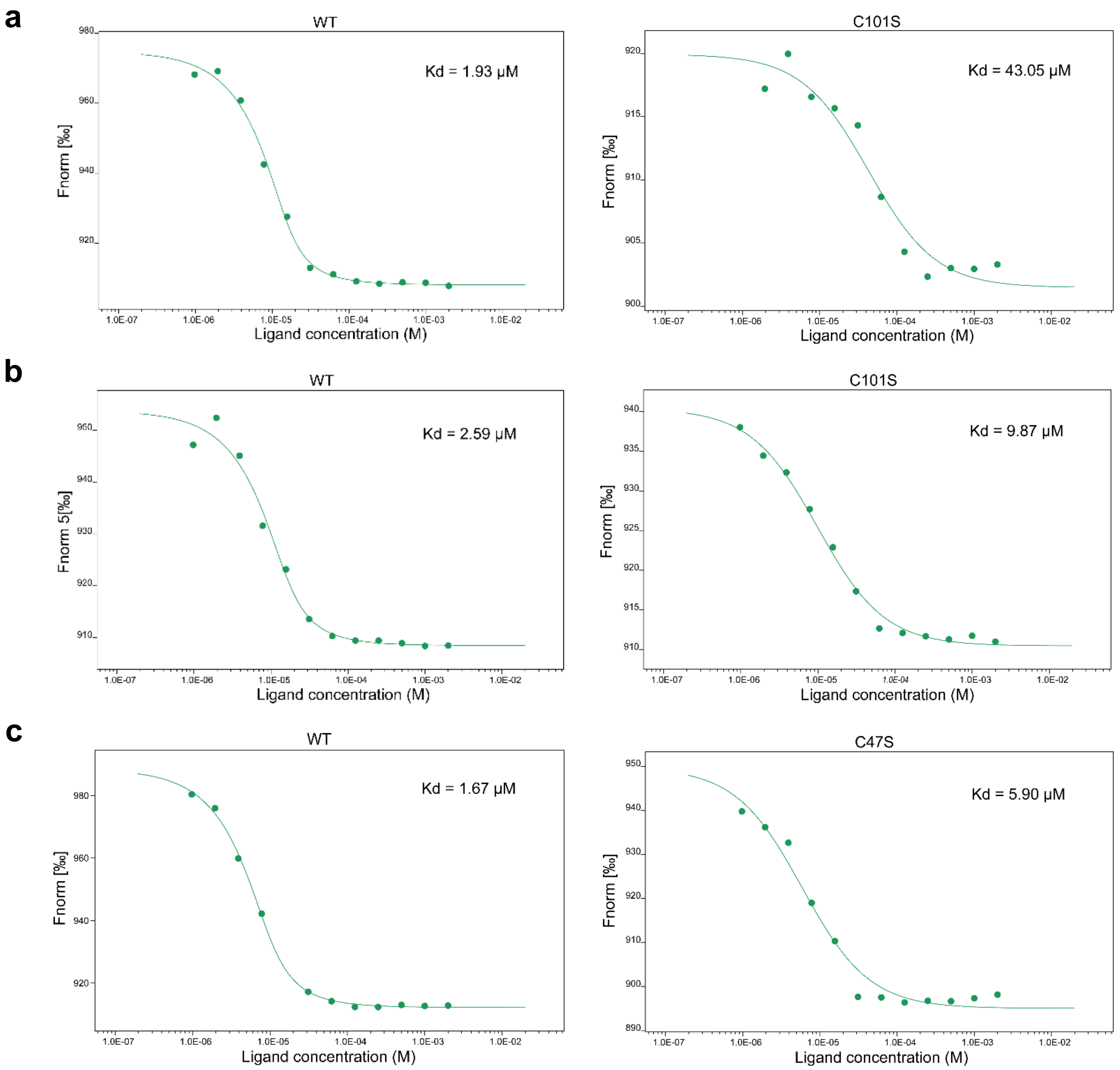

**Supplementary Fig. 9**

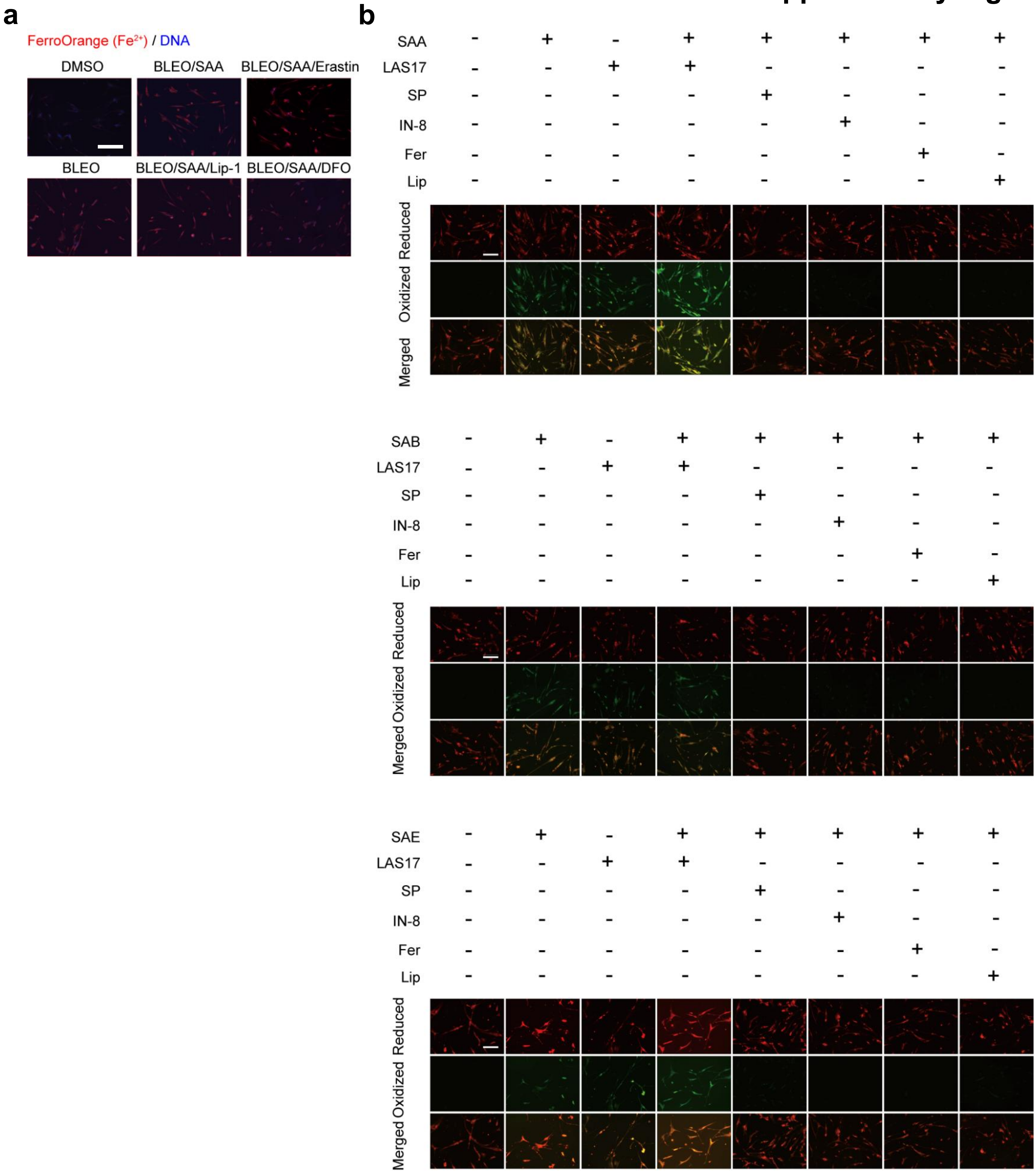

**Supplementary Fig. 10**

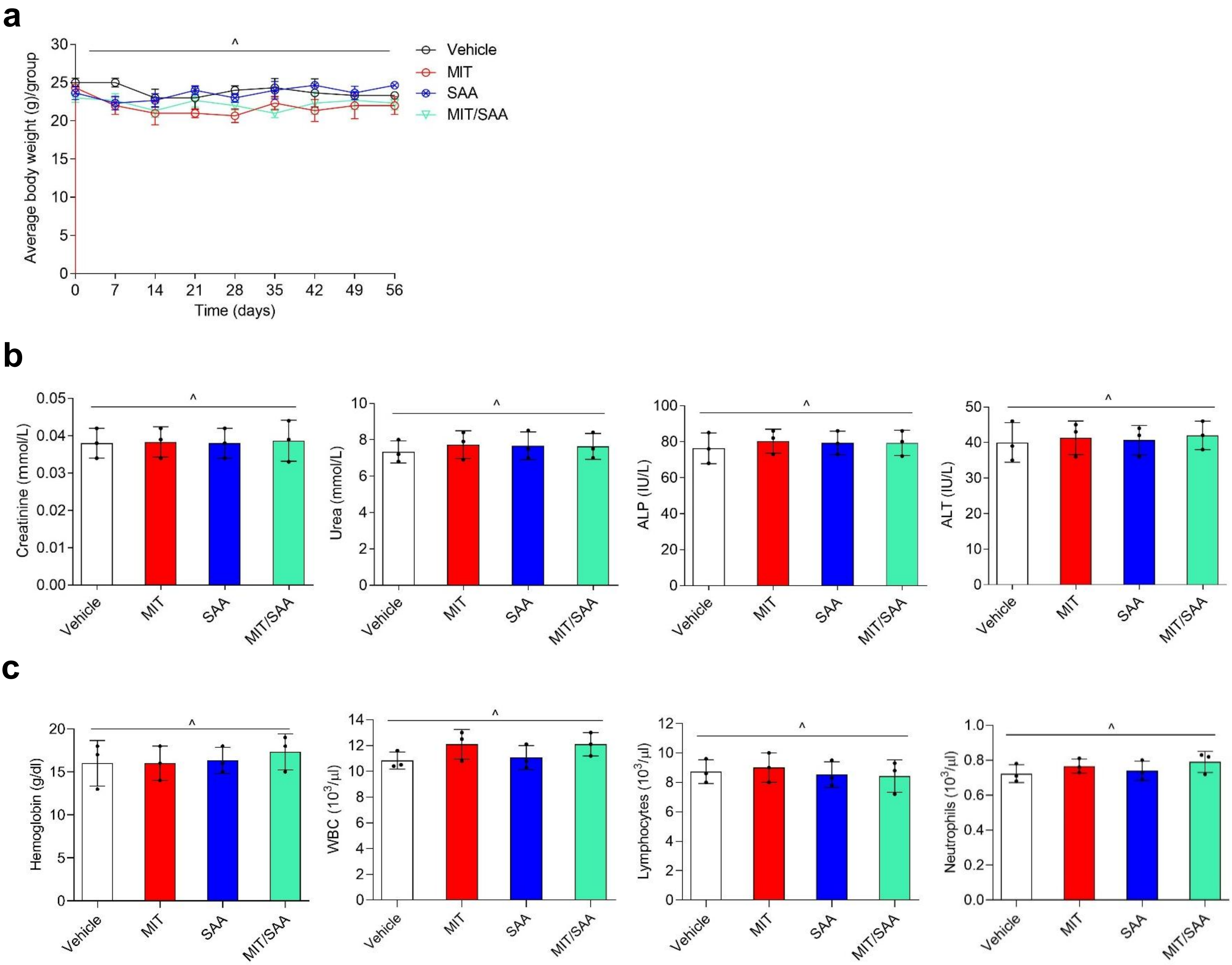

### Supplementary Fig. 11

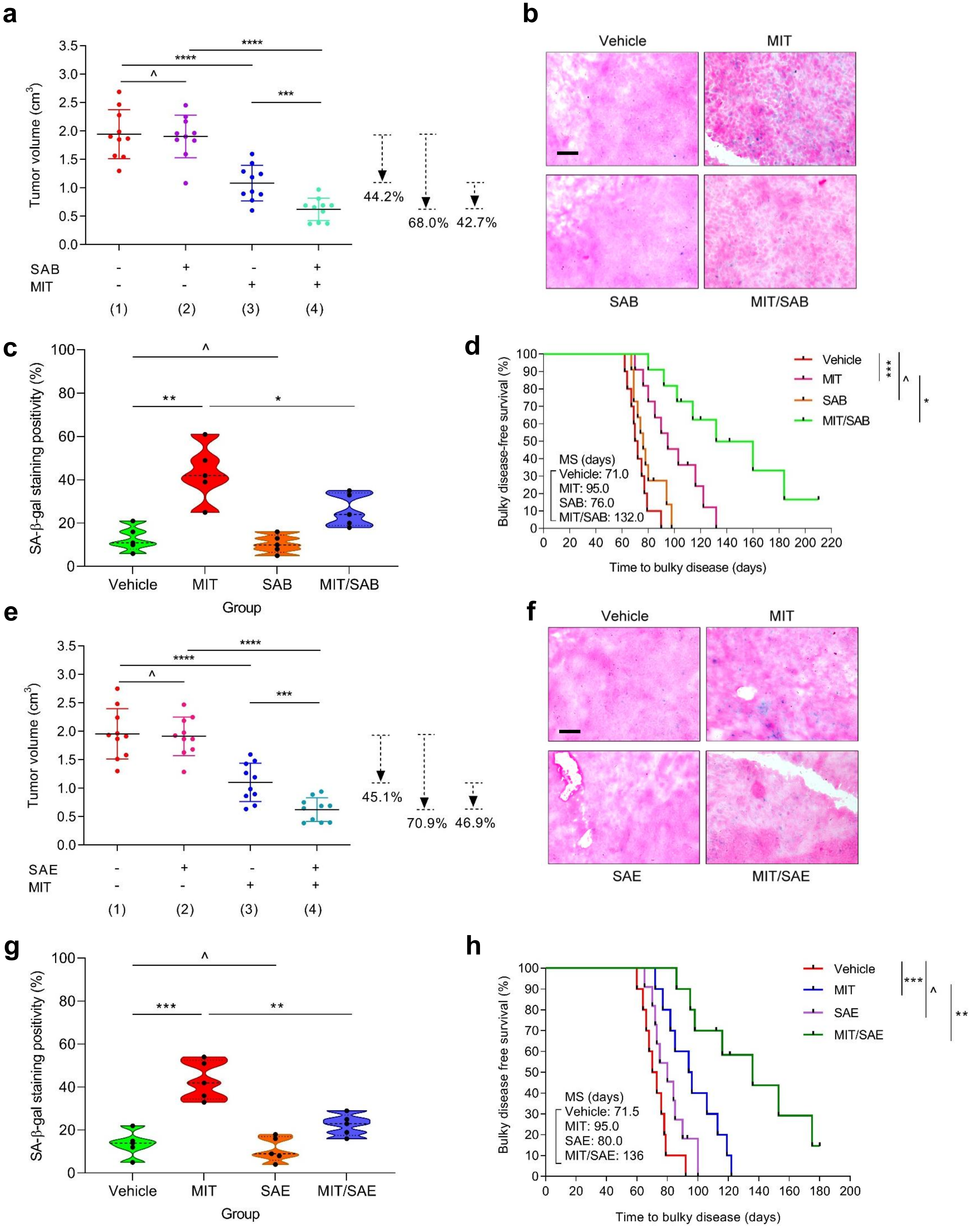

#### Supplementary Fig. 12

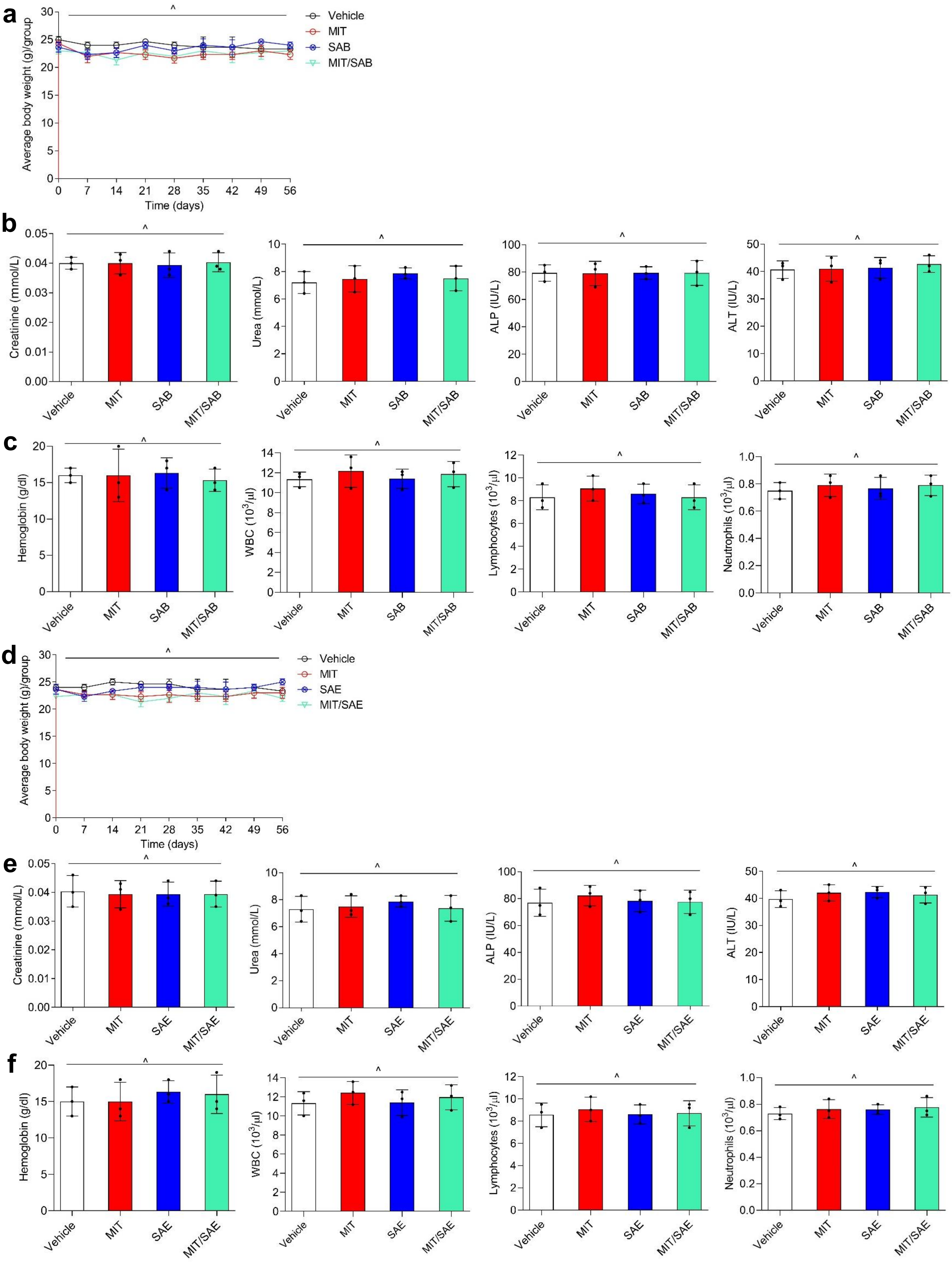

### Supplementary Fig. 13

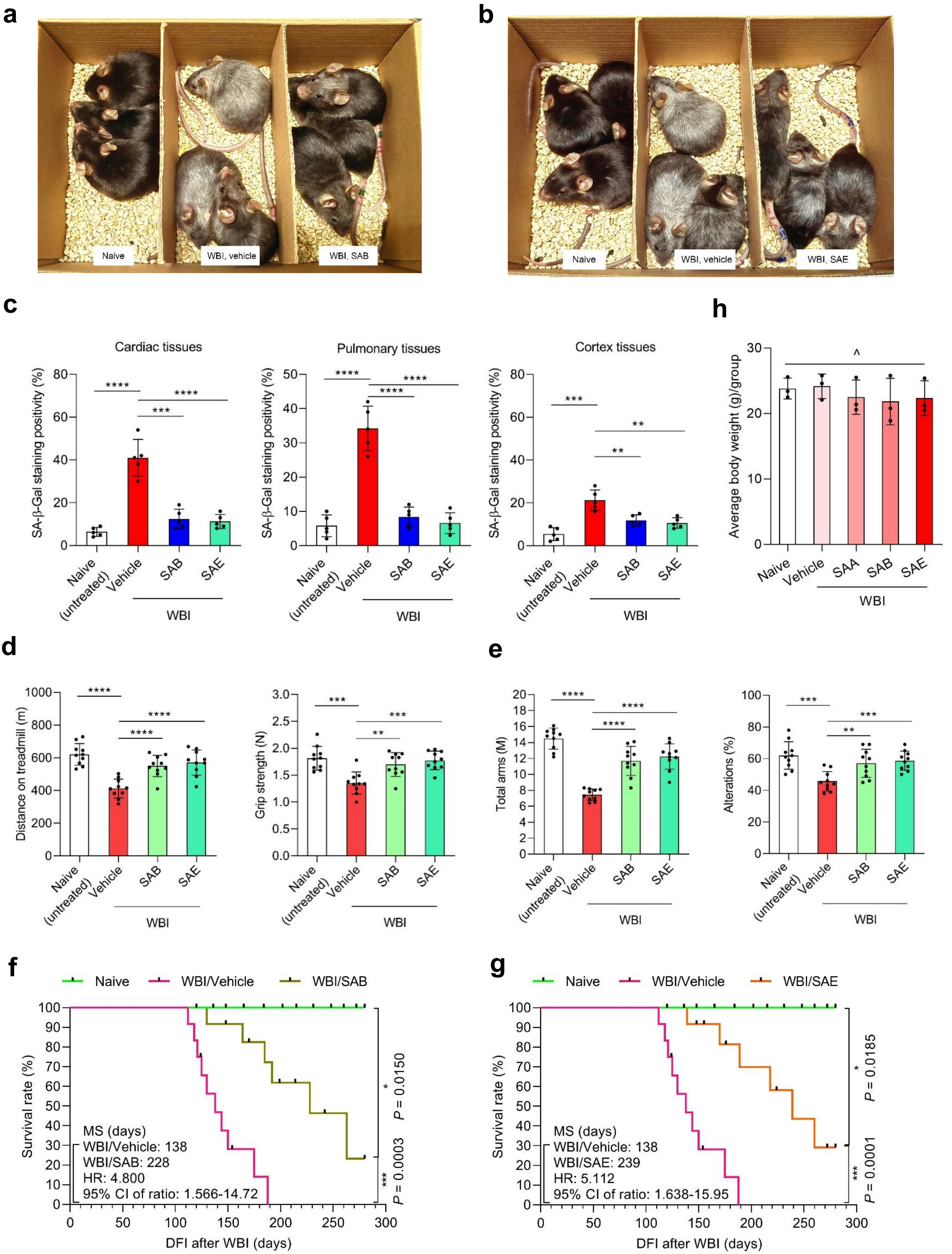

#### Supplementary Fig. 14

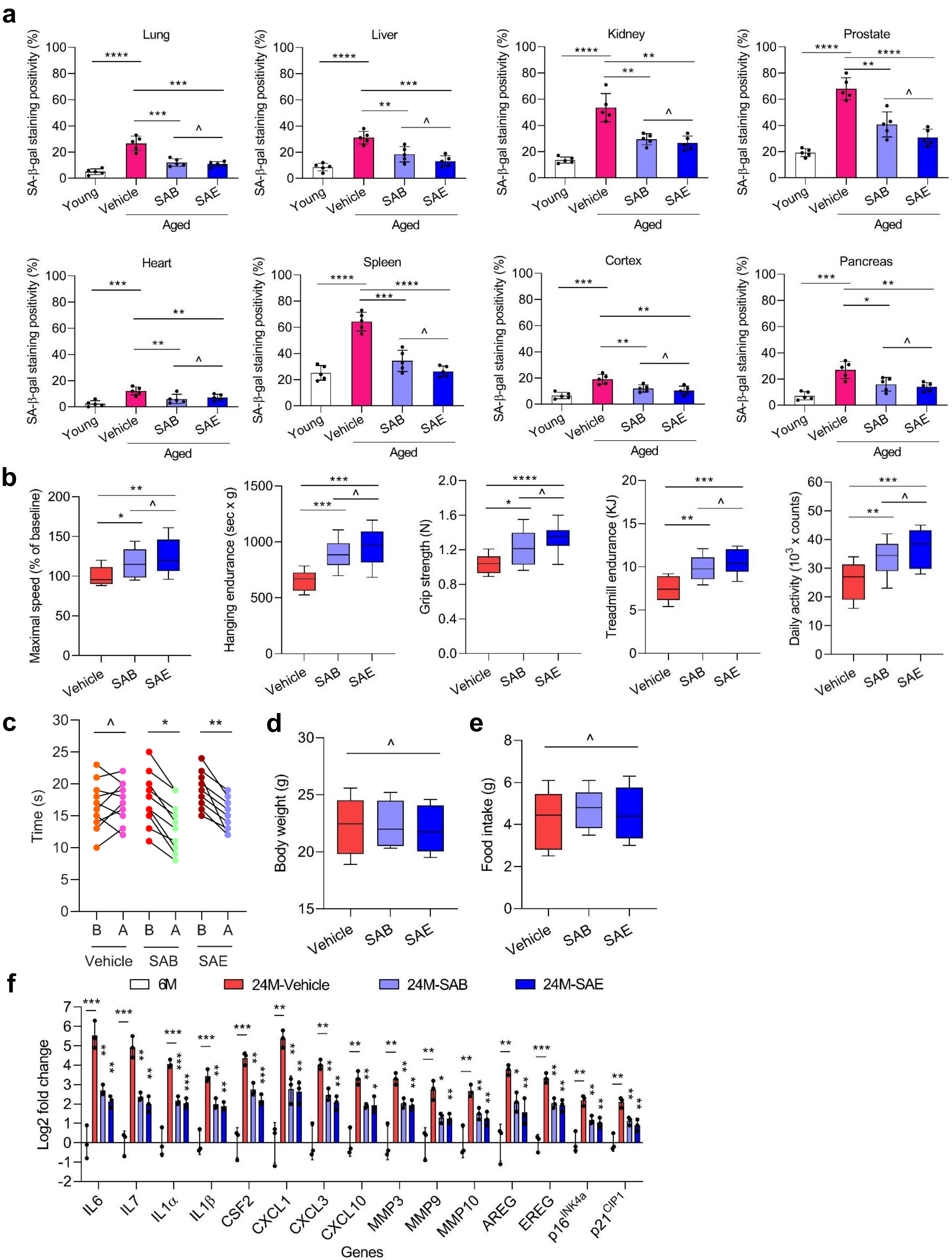

### Supplementary Fig. 15

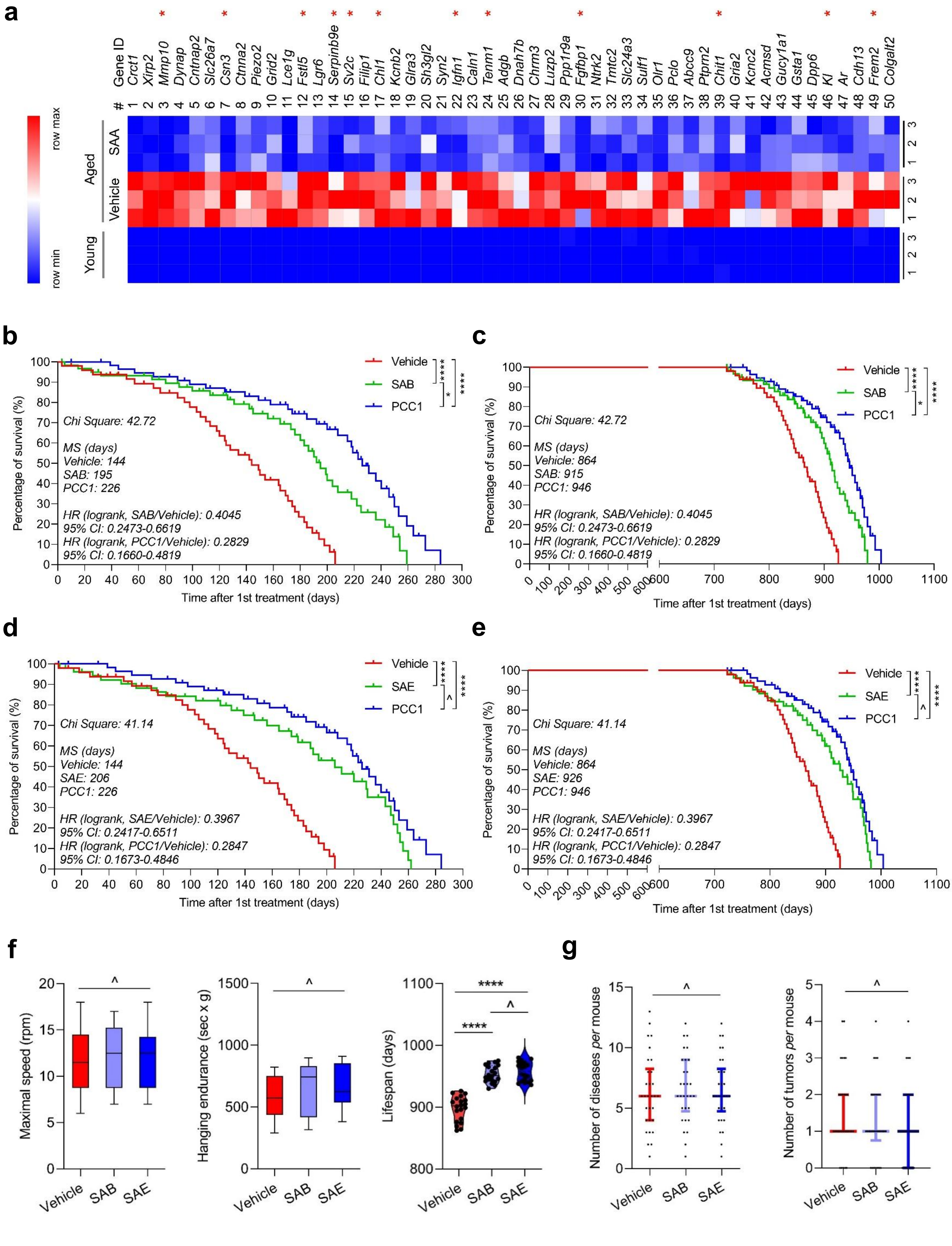

#### Supplementary Figure Legends

**Supplementary Fig. 1 Characterization of PSC27 as a cell-based model for senescence studies.** (**a**) Senescence assessment by SA-β-gal staining of PSC27 cells. Left, representative images. Scale bar, 10 μm. Right, statistics. (**b**) DNA synthesis appraisal by BrdU incorporation. Left, representative images, Scale bar, 10 μm. Right, statistics. (**c**) DNA damage response (DDR) profiling of cells. Left, representative images. Scale bar, 5 μm. Right, statistics. Immunofluorescence staining of γh2AX was performed with PSC27 cells, with the DDR profile categorized into 4 subgroups including 0 foci, 1-3 foci, 4-10 foci and >10 foci *per* cell. (**d**) Analysis of the senolytic potential of several agents, including ABT-263, ABT-737, 'D+Q' and PCC1. 'D+Q', dasatinib and quercetin. (**e**) Representative results from assessment of the senomorphic efficacy of natural compounds by analyzing the expression of a canonical SASP soluble factor CXCL8. (**f**) The efficacy of 56 compounds in the NMA library in killing senescent cells. Note, SAA, SAB, SAE, SAF and PCC1 showed the highest capacity as senolytics. However, SAF was subsequently excluded mainly due to its cytotoxicity to proliferating cells. (**g**) Expression of CXCL8 at transcription level in PSC27 cells after treatment by various agents that showed senomorphic potential. For all datasets, samples were examined after treatment with individual agents in culture for 3 days. Data in **a**-**e** are shown as mean ± SD and representative of 3 independent biological replicates with $P$ values calculated by Student's $t$-tests. ^, $P > 0.05$; *, $P < 0.05$; **, $P < 0.01$; ***, $P < 0.001$; ****, $P < 0.0001$.

**Supplementary Fig. 2 Characterization of the senolytic potential of SA compounds against different types of senescence.** (**a**) Cell counting-based survival evaluation of control (CTRL) cells and their senescent (SEN) counterparts generated by consecutive passaging, or subsequently exposed to increasing concentrations of SAA. (**b**) Cellular apoptosis examination by caspase 3/7 activity measurement upon exposure of SEN cells to SAA at 100 μM as described in (**a**). (**c**) Cell counting-based survival appraisal of CTRL cells and their SEN counterparts generated by overexpression of oncogenic *HRAS*$^{G12V}$, or subsequently exposed to increasing concentrations of SAA. (**d**) Cellular apoptosis appraisal by caspase 3/7 activity measurement upon exposure of senescent cells to SAA at 100 μM as described in (**c**). (**e**) Cell counting-based survival appraisal of CTRL cells and their SEN counterparts generated by consecutive passaging and subsequently exposed to increasing concentrations of SAB. (**f**) Cellular apoptosis examination by caspase 3/7 activity measurement upon exposure of senescent cells to SAB at 100 μM as described in (**e**). (**g**) Cell counting-based survival appraisal of CTRL cells and their SEN counterparts generated by overexpression of oncogenic *HRAS*$^{G12V}$ and subsequently exposed to increasing concentrations of SAB. (**h**) Cellular apoptosis appraisal by caspase 3/7 activity measurement upon exposure of senescent cells to SAB at 100 μM as described in (**g**). (**i**) Cell counting-based survival appraisal of CTRL cells and their SEN counterparts generated by consecutive passaging and subsequently exposed to increasing concentrations of SAE. (**j**) Cellular apoptosis examination by caspase 3/7 activity measurement upon exposure of senescent cells to SAE at 50 μM as described in (**i**). (**k**) Cell counting-based survival appraisal of CTRL cells and their SEN counterparts generated by overexpression of oncogenic *HRAS*$^{G12V}$ and subsequently exposed to increasing concentrations of SAE. (**l**) Cellular apoptosis appraisal by caspase 3/7 activity measurement upon exposure of senescent cells to SAE at 50 μM as described in (**k**). Data in **a**-**l** are shown as mean ± SD and representative of 3 independent biological replicates with $P$ values calculated by Student's $t$-tests. ^, $P > 0.05$; *, $P < 0.05$; **, $P < 0.01$; ***, $P < 0.001$; ****, $P < 0.0001$.

**Supplementary Fig. 3 Stromal cell-based *in vitro* assays identify SA compounds as broad-spectrum senolytics.** (**a**-**c**) Quantitative measurements of the viability of IMR90 (**a**), HUVEC (**b**) and AT-MSC (**c**) cells examined in CTRL, TIS, RS and OIS groups treated by 100 μM SAA for 3 d. (**d**-**f**) Quantitative measurements of the viability of IMR90 (**d**), HUVEC (**e**) and AT-MSC (**f**) cells examined in CTRL, TIS, RS and OIS groups treated by 100 μM SAB for 3 d. (**g**-**i**) Quantitative measurements of the viability of IMR90 (**g**), HUVEC (**h**) and AT-MSC (**i**) cells examined in CTRL, TIS, RS and OIS groups treated by 50 μM SAE for 3 d. CTRL, control. TIS, therapy-induced senescence. RS, replicative senescence. OIS, oncogene-induced senescence. Data in **a**-**i** are shown as mean ± SD and representative of 3 independent biological replicates with *P* values calculated by Student's *t*-tests. ^, $P > 0.05$; *, $P < 0.05$; **, $P < 0.01$.

**Supplementary Fig. 4 Bioinformatics profiling the expression pattern of senescent cells exposed to a sublethal dose of SAA.** (**a**) Heatmap showing the expression of human genes, which were significantly upregulated (1270) or downregulated (2089) upon treatment of senescent PSC27 cells by SAA (50 μM). Data derived from triplicate samples *per* group. (**b**-**d**) Pie charts depicting the biological process (BP) (**b**), cellular component (CC) (**c**) and molecular function (MF) (**d**) associated with transcripts upregulated by BLEO but downregulated by SAA as disclosed by GO analysis.

**Supplementary Fig. 5 Immunoblot analysis of PSC27 cells treated under different conditions involving BLEO and/or SAB (or SAE).** Expression of DDR signaling molecules, SASP regulatory factors, pro-apoptotic and anti-apoptotic proteins, and anti-ferroptotic factors was examined. Caspase 3 (t), total caspase 3; caspase 3 (c), cleaved caspase 3; p, phosphorylated. GAPDH, loading control.

**Supplementary Fig. 6 Illustration of chemical synthesis to conjugate biotin with SA compounds and venn diagram profiling potential targets of these SA compounds.** (**a**-**c**) A schematic workflow of chemical synthesis for SAA-Biotin (**a**), SAB-Biotin (**b**) and SAC-Biotin (**c**), respectively. The chemical structure of biotin is highlighted by the dashed black rectangle in **a**-**c**.

**Supplementary Fig. 7 Generation of biotin-conjugated complexes to probe potential targets of SA compounds and illustration of target-specific GO profiling.** (**a**) Representative presentation of human protein microarrays (20K) applied to probe the potential targets of SA compounds, with DMSO-treated biotin (D-Biotin) employed as negative control. (**b**) Venn diagram outlining all the potential co-targets (154) of SAA, SAB and SAE compounds as identified by pulldown assays. (**c**) GO profiling of the top 100 molecules out of the 154 co-targets of SA compounds identified by protein microarrays, with all relevant GO items presented for a quick view. (**d-f**) Venn diagrams showing the potential targets of SAA (**d**), SAB (**e**) and SAE (**f**), respectively, as co-identified by protein microarrays and pulldown assays. (**g**) Crystal structure of human dimeric GSTP1 showing experimentally-determined 3D architectures derived from the Protein Data Bank (PDB) (ID: 2A2R) archive (method, X-Ray diffraction; resolution, 1.40 Å)[1]. The asymmetric unit of the crystal contains 2 GSTP1 monomers. Red, α-helix; yellow, β-pleated sheets; blue, β-bend; grey, random coil.

**Supplementary Fig. 8 MST-based profiling of the interaction between SA compounds and GSTP1 or its functionally disruptive mutants and performance of cell death assessment.** (**a**) Interaction between SAA and GSTP1 overexpressed in cells with a GFP tag was assessed by the cellular microscale thermophoresis analysis (MST). Left, WT. Right, the C101S mutant. (**b**) Interaction between SAB and GSTP1 overexpressed in cells with a GFP tag was assessed by the cellular MST. Left, WT. Right, the C101S mutant. (**c**) Interaction between SAE and GSTP1 overexpressed in cells with a GFP tag was assessed by the cellular MST. Left, WT. Right, the C47S mutant.

**Supplementary Fig. 9. Determination of the potential of SA compounds in inducing ferroptosis in senescent cells.** (**a**) PSC27 cells were induced to senesce for 7days and stained with FerroOrange to quantify intracellular $Fe^{2+}$. Scale bar, 10 μm. (**b**) SAA, SAB or SAE was added to cell culture media, either alone or together with LAS17, SP, IN-8, Fer, Lip. The chemical agent 4,4-difluoro-4-bora-3a,4a-diaza-s-indacene (BODIPY-C11), was used to assess the levels of ferroptosis in cells. Note, BODIPY-C11 is a BODIPY-based, red fluorescent dye with a fatty acid modification. Upon oxidation by free radicals, its emission fluorescence shifts towards shorter wavelengths, resulting in green fluorescence. LAS17, L-buthionine sulfoximine analogue 17. SP, SP600125. IN-8, JNK-IN-8. Fer, Ferrostatin-1. Lip, Liproxstatin-1. SP and IN-8 are chemical inhibitors of JNK. Fer and Lip are chemical inhibitors of ferroptosis. Scale bar, 10 μm.

**Supplementary Fig. 10 Pathophysiological assessment of SAA treatment effects on immunocompetent mice (males).** (**a**) Animal body weight determination performed on a weekly basis for immunocompetent C57BL/6J mice. (**b**) Serum measurement of creatinine, urea, alkaline phosphatase (ALP), and alanine aminotransferase (ALT) with terminal bleeds (cardiac punctures) taken at the end of therapeutic regimens. (**c**) Routine analysis of peripheral blood. The circulating levels of hemoglobin, white blood cells, lymphocytes and platelets at the end of each therapeutic regimen were assessed. Data are shown as mean ± SD and representative of 3 independent experiments. MIT, mitoxantrone. WBC, white blood count. For all datasets, n = 3 *per* treatment arm. For all datasets, *P* values were calculated by one-way ANOVA with Tukey's multiple-comparison test. ^, $P > 0.05$.

**Supplementary Fig. 11 Senolysis by SAB/SAE in the treatment-damaged tumor microenvironment restrains SASP-conferred cancer resistance. (a)** Comparative statistics of tumor end volumes. PC3 cancer cells were inoculated either alone or combined with PSC27 stromal cells before implanted subcutaneously to the hind flank of NOD/SCID animals. The mice were then administered with MIT and SAB, either alone or in combination. (**b**) Representative images of *in vivo* senescence in tumors by SA-β-gal staining. Scale bar, 50 μm. (**c**) Comparative statistics of tumor senescence as described in (**b**). (**d**) Comparative survival of animals (MIT/SAB) sacrificed upon development of advanced bulky diseases. **(e)** Comparative statistics of tumor end volumes. PC3 cancer cells were inoculated either alone or combined with PSC27 stromal cells before implanted subcutaneously to the hind flank of NOD/SCID animals, which were administered with MIT and SAE, either alone or in combination. (**f**) Representative images of *in vivo* senescence in tumors by SA-β-gal staining. Scale bar, 50 μm. (**g**) Comparative statistics of tumor senescence as described in (**f**). (**h**) Comparative survival of animals (MIT/SAE) sacrificed upon development of advanced bulky diseases. Data in **a**, **c**, **e** and **g** are shown as mean ± SD and representative of 3 independent experiments. In **d** and **h,** survival duration was calculated from tissue recombinant injection until death. *P* values were calculated by a two-sided log-rank (Mantel-Cox) test. MIT, mitoxantrone. For each dataset, $n$ = 10 *per* treatment arm. ^, $P > 0.05$; *, $P < 0.05$; **, $P < 0.01$; ***, $P < 0.001$; ****, $P < 0.0001$.

**Supplementary Fig. 12 Pathophysiological assessment of SAB/SAE treatment effects on immunocompetent mice (males).** (**a**) Animal body weight determination performed on a weekly basis for immunocompetent C57BL/6J mice treated with MIT/SAB. (**b**) Serum measurement of creatinine, urea, ALP and ALT with terminal bleeds (cardiac punctures) taken at the end of therapeutic regimens of animals treated with MIT/SAB. (**c**) Routine analysis of peripheral blood from animals treated with MIT/SAB. The circulating levels of hemoglobin, white blood cells, lymphocytes and platelets at the end of each therapeutic regimen were assessed. (**d**) Animal body weight determination performed on a weekly basis for immunocompetent C57BL/6J mice treated with MIT/SAE. (**e**) Serum measurement of creatinine, urea, ALP and ALT with terminal bleeds (cardiac punctures) taken at the end of therapeutic regimens of animals treated with MIT/SAE. (**f**) Routine analysis of peripheral blood from animals treated with MIT/SAE. The circulating levels of hemoglobin, white blood cells, lymphocytes and platelets at the end of each therapeutic regimen were assessed. Data are shown as mean ± SD and representative of 3 independent experiments. MIT, mitoxantrone. ALP, alkaline phosphatase. ALT, alanine aminotransferase. WBC, white blood count. For all datasets, n = 3 *per* treatment arm. For all datasets, *P* values were calculated by one-way ANOVA with Tukey's multiple-comparison test. ^, $P > 0.05$.

**Supplementary Fig. 13 SAB/SAE treatment mitigates physical dysfunction of animals exposed to WBI.** (**a**) Whole body snapshoot comparison of C57BL/6J males that were naïve, WBI-exposed followed by vehicle-treatment or WBI-exposed followed by SAB (**a**) or SAE (**b**) treatment, respectively. (**c**) Comparative statistics of SA-β-gal staining positivity of cardiac (left), pulmonary (middle) and cortex (right) tissues of animals either naïve, WBI followed by treatment with SAB or SAE, respectively. (**d**) Measurement of running distance on treadmill (left) and grip strength (right) for animals in each group as described in (**c**). (**e**) Y-maze test for experimental mice. Left, comparative total arms. Right, comparative alterations. (**f**, **g**) Comparative survival of naïve, WBI mice followed by treatment with either SAB (**f**) or SAE (**g**). (**h**) Quantitative comparison of body weight of animals treated as indicated. *P* values were calculated by a two-sided log-rank (Mantel-Cox) test. Data in **c**-**e** are shown as mean ± SD and representative of 3 independent biological replicates, with *P* values calculated by Student's *t*-tests. ^, $P > 0.05$; *, $P < 0.05$; **, $P < 0.01$; ***, $P < 0.001$; ****, $P < 0.0001$.

**Supplementary Fig. 14 SAB/SAE treatment eliminates senescent cells in multiple organs.** (**a**) Comparative statistics of SA-β-gal staining positivity in the lung, liver, kidney, prostate, heart, spleen, cortex and pancreas tissues of young, aged mice treated with vehicle, SAB or SAE. (**b**) Quantification of maximal walking speed (relative to baseline), hanging endurance, grip strength, treadmill endurance and daily activity of 20-month-old C57BL/6J males after 4-month intervention with SAB or SAE. (**c**) Quantification of the time to cross the balance beam. Data before and after treatment of each animal are connected to allow direct comparison of treatment effects. (**d**) Comparison of body weight of animals as described in **Fig. 8a**. (**e**) Measurement of food intake of animals as described in **Fig. 8a**. (**f**) Quantitative transcript profiling of SASP expression in lung tissues collected from 6-month-old untreated (6M), 24-month-old vehicle-treated (24M-Vehicle), 24-month-old SAB-treated mice (24M-SAB) and 24-month-old SAE-treated mice (24M-SAE), respectively. Data are shown as mean ± SD and derive from 3 biological replicates (*n* = 3 independent assays). *P* values were calculated by Student's *t*-tests. ^, $P > 0.05$; *, $P < 0.05$; **, $P < 0.01$; ***, $P < 0.001$; ****, $P < 0.0001$.

**Supplementary Fig. 15 SAA suppresses the SASP expression *in vivo* and intervention with SAB/SAE prolongs survival of aged mice.** (**a**) Heatmap depicting top genes (50) significantly upregulated in lung tissues of mice but subject to reversal by SAA intervention. (**b**-**e**) Post-treatment survival (**b**,**d**) and whole-life survival (**c**,**e**) curves of C57BL/6J animals treated biweekly with SAB (*n* = 35; 18 males, 17 females) or SAE (*n* = 33; 18 males, 15 females), or vehicle (*n* = 39; 20 males, 19 females) starting at 24-27 months of age. PCC1 was administered as a positive control (*n* = 32; 17 males, 15 females). (**f**) Maximal walking speed and hanging endurance averaged over the last 2 months of life for mice treated with SAB/SAE (*n* = 10/group), and lifespan for the longest living mice (top 20) in both groups. (**g**) Disease burden and tumor burden at death of mice treated with SAB/SAE. For both sexes, *n* (animal number) = 30 *per* arm in each case. Cox proportional hazard regression model (**b**-**e**) and one-way ANOVA with Tukey's multiple-comparison test (**f**-**g**). ^, $P > 0.05$; *, $P < 0.05$; **, $P < 0.01$; ***, $P < 0.001$; ****, $P < 0.0001$.

#### Supplementary Table 1. A list of naturally derived agents in the NMA library

| No. | ID | Name | Synonyms | CAS | SMILES | Formula | Molecular weight |
|---|---|---|---|---|---|---|---|
| 1 | Flavin | Flavin mononucleotide | Riboflavin 5'-phosphate sodium;Vitamin B2 Phosphate Sodium Salt;FMN-Na;Riboflavin phosphate sodium;riboflavin-5'-phosphate;FMN | 130-40-5 | [Na+].c12c(nc3c(n1C[C@@H]([C@@H]([C@@H](COP(=O)([O-])O)O)O)O)cc(c(C)c3)C)c(=O)[nH]c(=O)n2 | C17H20N4NaO9P | 478.33 |
| 2 | Teniposide | Teniposide | VM26;NSC 122819 | 29767-20-2 | O1COc2c1cc1[C@H]([C@@H]3[C@@H]([C@@H](c1c2)c1cc(c(c(c1)OC)O)OC)C(=O)OC3)O[C@H]1[C@@H]([C@H]([C@@H]2O[C@@H](OC[C@H]2O1)c1cccs1)O)O | C32H32O13S | 656.66 |
| 3 | 7-Hydroxycoumarin | 7-Hydroxycoumarin | Hydrangin;Hydrangine;NSC 19790;Skimmetine;Umbelliferone | 93-35-6 | Oc1cc2c(cc1)ccc(=O)o2 | C9H6O3 | 162.14 |
| 4 | Methyl protocatechuate | Methyl protocatechuate | Methyl 3,4-dihydroxybenzoate;Protocatechuic acid methyl ester;3,4-Dihydroxybenzoic acid methyl ester | 2150-43-8 | COC(=O)c1cc(O)c(O)cc1 | C8H8O4 | 168.15 |

| | | | | | | | |
|---|---|---|---|---|---|---|---|
| 5 | Echinacoside | Echinacoside | | 82854-37-3 | c1(ccc(cc1O)/C=C/C(=O)O[C@H]1[C@@H]([C@H]([C@@H](O[C@@H]1CO[C@@H]1O[C@@H]([C@H]([C@@H]([C@H]1O)O)O)CO)OCCc1ccc(c(c1)O)O)O)O[C@@H]1O[C@H]([C@@H]([C@H]([C@H]1O)O)O)C)O | C35H46O20 | 786.73 |
| 6 | Genistein | Genistein | NPI 031L | 446-72-0 | c12c(c(=O)c(co1)c1ccc(cc1)O)c(cc(c2)O)O | C15H10O5 | 270.24 |
| 7 | Taxifolin | Taxifolin | (+)-Taxifolin;(+)-Dihydroquercetin;Dihydroquercetin | 480-18-2 | Oc1ccc(cc1O)[C@@H]1[C@@H](O)C(=O)c2c(O)cc(O)cc2O1 | C15H12O7 | 304.25 |
| 8 | Kaempferol | Kaempferol | Kempferol;Robigenin | 520-18-3 | Oc1ccc(cc1)c1c(O)c(=O)c2c(O)cc(O)cc2o1 | C15H10O6 | 286.23 |
| 9 | Theophylline-7-acetic acid | Theophylline-7-acetic acid | Acefylline;Theophyllineacetic acid;acetyloxytheophylline;Carboxymethyltheophylline | 652-37-9 | Cn1c2c(c(=O)n(c1=O)C)n(cn2)CC(=O)O | C9H10N4O4 | 238.2 |
| 10 | Salidroside | Salidroside | Rhodioloside | 10338-51-9 | c1cc(ccc1CCO[C@H]1[C@@H]([C@H]([C@@H]([C@H](O1)CO)O)O)O)O | C14H20O7 | 300.3 |
| 11 | Ketoisophorone | Ketoisophorone | 4-Oxoisophorone;2,6,6-Trimethyl-2-cyclohexene-1,4-dione | 1125-21-9 | CC1=CC(=O)CC(C)(C)C1=O | C9H12O2 | 152.19 |

| | | | | | | | |
|---|---|---|---|---|---|---|---|
| 12 | Rhoifolin | Rhoifolin | Rhoifoloside;Apigenin 7-O-neohesperidoside;Apigenin-7-O-rhamnoglucoside | 17306-46-6 | c1(cc(=O)c2c(cc(cc2o1)O[C@H]1[C@@H]([C@H]([C@@H]([C@H](O1)CO)O)O)O[C@H]1[C@@H]([C@@H]([C@H]([C@@H](O1)C)O)O)O)O)c1ccc(cc1)O | C27H30O14 | 578.53 |
| 13 | Swertiamarin | Swertiamarin | Swertiamaroside | 17388-39-5 | OC[C@H]1O[C@@H](O[C@@H]2OC=C3C(=O)OCC[C@@]3(O)[C@H]2C=C)[C@H](O)[C@@H](O)[C@@H]1O | C16H22O10 | 374.34 |
| 14 | Loganin | Loganin | Loganoside | 18524-94-2 | C[C@H]1[C@H](C[C@H]2[C@@H]1[C@@H](OC=C2C(=O)OC)O[C@H]1[C@@H]([C@H]([C@@H]([C@H](O1)CO)O)O)O)O | C17H26O10 | 390.38 |
| 15 | 10-Hydroxycamptothecin | 10-Hydroxycamptothecin | (S)-10-Hydroxycamptothecin;10-HCPT | 19685-09-7 | CC[C@@]1(O)C(=O)OCc2c1cc1-c3nc4ccc(O)cc4cc3Cn1c2=O | C20H16N2O5 | 364.36 |
| 16 | Neohesperidin Dihydrochalco | Neohesperidin Dihydrochalcone | NHDC;Neohesperidin DC;NCI-c60764 | 20702-77-6 | C[C@H]1[C@@H]([C@H]([C@H]([C@@H](O1)O[C@@H]1[C@H]([C@@H]([C@H](O[C@H]1Oc1cc(c(c(c1)O)C(=O)CCc1cc(c(cc1)OC)O)O)CO)O)O)O)O)O | C28H36O15 | 612.58 |

| | | | | | | | |
|---|---|---|---|---|---|---|---|
| 17 | Gentiopicrin | Gentiopicrin | Gentiopicroside | 20831-76-9 | C=C[C@H]1[C@@H](OC=C2C1=CCOC2=O)O[C@H]1[C@@H]([C@H]([C@@H]([C@H](O1)CO)O)O)O | C16H20O9 | 356.33 |
| 18 | Orcinol glucoside | Orcinol glucoside | Sakakin | 21082-33-7 | Cc1cc(O)cc(O[C@@H]2O[C@H](CO)[C@@H](O)[C@H](O)[C@H]2O)c1 | C13H18O7 | 286.28 |
| 19 | Baicalin | Baicalin | Baicalein 7-O-β-D-glucuronide | 21967-41-9 | c1ccc(cc1)c1cc(=O)c2c(c(c(cc2o1)O[C@H]1[C@@H]([C@H]([C@@H]([C@H](O1)C(=O)O)O)O)O)O)O | C21H18O11 | 446.37 |
| 20 | Catalpol | Catalpol | Catalpinoside;Digitalis purpurea L | 2415-24-9 | OC[C@H]1O[C@@H](O[C@@H]2OC=C[C@H]3[C@H](O)[C@@H]4O[C@]4(CO)[C@@H]23)[C@H](O)[C@@H](O)[C@@H]1O | C15H22O10 | 362.33 |
| 21 | Sanguinarine | Sanguinarine | Sanguinarin;Pseudochelerythrine | 2447-54-3 | C[n+]1c2c(c3ccc4c(c3c1)OCO4)ccc1cc3c(cc21)OCO3 | C20H14NO4 | 332.33 |
| 22 | Picroside I | Picroside I | 6'-Cinnamoylcatalpol | 27409-30-9 | OC[C@]12O[C@H]1[C@@H](O)[C@@H]1C=CO[C@@H](O[C@@H]3O[C@H](COC(=O)\C=C\c4ccccc4)[C@@H](O)[C@H](O)[C@H]3O)[C@H]21 | C24H28O11 | 492.47 |
| 23 | Scutellarin | Scutellarin | Breviscapinun;Breviscapine;Scutellarein-7-glucuronide;Breviscapin | 27740-01-8 | c1cc(ccc1c1cc(=O)c2c(c(c(cc2o1)O[C@H]1[C@@H]([C@H]([C@@H]([C@H](O1)C(=O)O)O)O)O)O)O)O | C21H18O12 | 462.37 |

| | | | | | | | |
|---|---|---|---|---|---|---|---|
| 24 | Oridonin | Oridonin | Isodonol;NSC-250682;Rubescenin;Rubescensin A | 28957-04-2 | CC1(C)CC[C@H](O)[C@]23CO[C@](O)([C@@H](O)[C@H]12)[C@@]12[C@H](O)[C@@H](CC[C@@H]31)C(=C)C2=O | C20H28O6 | 364.44 |
| 25 | Pectolinarin | Pectolinarin | | 28978-02-1 | COc1ccc(cc1)-c1cc(=O)c2c(O)c(OC)c(O[C@@H]3O[C@H](CO[C@@H]4O[C@@H](C)[C@H](O)[C@@H](O)[C@H]4O)[C@@H](O)[C@H](O)[C@H]3O)cc2o1 | C29H34O15 | 622.59 |
| 26 | Caffeic acid | Caffeic Acid Phenethyl Ester | CAPE;Phenylethyl Caffeate | 104594-70-9 | c1ccc(cc1)CCOC(=O)/C=C/c1ccc(c(c1)O)O | C17H16O4 | 284.31 |
| 27 | Madecassoside | Madecassoside | Asiaticoside A | 34540-22-2 | C[C@@H]1CC[C@@]2(CC[C@]3(C)C(=CC[C@@H]4[C@@]5(C)C[C@@H](O)[C@H](O)[C@@](C)(CO)[C@H]5[C@H](O)C[C@@]34C)[C@@H]2[C@H]1C)C(=O)O[C@@H]1O[C@H](CO[C@@H]2O[C@H](CO)[C@@H](O[C@@H]3O[C@@H](C)[C@H](O)[C@@H](O)[C@H]3O)[C@H](O)[C@H]2O)[C@@H](O)[C@H](O)[C@H]1O | C48H78O20 | 975.12 |

| | | | | | | | |
|---|---|---|---|---|---|---|---|
| 28 | Picroside II | Picroside II | 6-Vanilloylcatalpol;Vanilloyl catalpol | 39012-20-9 | [C@@H]1(OC(=O)c2cc(c(cc2)O)OC)[C@H]2[C@@](O2)(CO)[C@@H]2[C@H]1C=CO[C@H]2O[C@H]1[C@@H]([C@H]([C@@H]([C@H](O1)CO)O)O)O | C23H28O13 | 512.46 |
| 29 | Methyl Vanillate | Methyl Vanillate | | 3943-74-6 | COC(=O)c1cc(OC)c(O)cc1 | C9H10O4 | 182.17 |
| 30 | Cryptotanshinone | Cryptotanshinone | Tanshinone c;Cryptotanshinon | 35825-57-1 | c12c(C(CCC1)(C)C)ccc1c2C(=O)C(=O)C2=C1OC[C@@H]2C | C19H20O3 | 296.37 |
| 31 | 7,8-Dihydroxyflavone | 7,8-Dihydroxyflavone | 7,8-Dihydroxyflavone;7,8-DHF | 38183-03-8 | c1ccc(cc1)c1cc(=O)c2c(o1)c(c(cc2)O)O | C15H10O4 | 254.24 |
| 32 | Ethyl ferulate | Ethyl ferulate | Ferulic acid ethyl ester;Ethyl 4'-hydroxy-3'-methoxycinnamate;Ethyl 3-(4-hydroxy-3-methoxyphenyl)acrylate | 4046-02-0 | CCOC(=O)/C=C/c1cc(c(cc1)O)OC | C12H14O4 | 222.24 |
| 33 | Asiatic acid | Asiatic acid | Dammarolic acid;Asiantic acid | 464-92-6 | C[C@@H]1CC[C@@]2(CC[C@@]3(C(=CC[C@H]4[C@]3(CC[C@@H]3[C@@]4(C[C@H]([C@@H]([C@@]3(C)CO)O)O)C)C)[C@@H]2[C@H]1C)C)C(=O)O | C30H48O5 | 488.7 |
| 34 | Betulinic acid | Betulinic acid | Betulic acid;Lupatic acid;ALS-357 | 472-15-1 | [C@H]12[C@@](CC[C@]3([C@@]4(CC[C@H]5C([C@H](CC[C@@]5([C@H]4CC[C@H]13)C)O)(C)C)C)C)(CC[C@H]2C(=C)C)C(=O)O | C30H48O3 | 456.71 |
| 35 | Dehydrocostus Lactone | Dehydrocostus Lactone | Epiligulyl oxide;(-)-Dehydrocostus lactone | 477-43-0 | C=C1CC[C@@H]2[C@H]1[C@H]1OC(=O)C(=C)[C@@H]1CCC2=C | C15H18O2 | 230.3 |

| | | | | | | | |
|---|---|---|---|---|---|---|---|
| 36 | Nobiletin | Nobiletin | Hexamethoxyflavone | 478-01-3 | COc1c(cc(cc1)c1cc(=O)c2c(o1)c(c(c(c2OC)OC)OC)OC)OC | C21H22O8 | 402.4 |
| 37 | Morin | Morin | Al-Morin;Aurantica;Calico Yellow;Toxylon pomiferum | 480-16-0 | Oc1cc(O)c2c(=O)c(O)c(c3c(O)cc(O)cc3)oc2c1 | C15H10O7 | 302.23 |
| 38 | Chrysin | Chrysin | 5,7-Dihydroxyflavone;NSC 407436;5, 7-Dihydroxyflavone | 480-40-0 | Oc1cc(O)c2c(=O)cc(oc2c1)c1ccccc1 | C15H10O4 | 254.24 |
| 39 | Naringenin | Naringenin | NSC 34875;S-Dihydrogenistein;NSC 11855;Salipurol;Naringetol;Pelargidanon | 480-41-1 | C1[C@H](Oc2cc(cc(c2C1=O)O)O)c1ccc(cc1)O | C15H12O5 | 272.26 |
| 40 | Tangeretin | Tangeretin | NSC53909;NSC618905;Tangeritin | 481-53-8 | COC1=CC=C(C=C1)C1=CC(=O)C2=C(O1)C(OC)=C(OC)C(OC)=C2OC | C20H20O7 | 372.38 |
| 41 | Hyperoside | Hyperoside | Hyperin;Quercetin 3-galactoside | 482-36-0 | c1cc(c(cc1c1c(c(=O)c2c(cc(cc2o1)O)O)O[C@H]1[C@@H]([C@H]([C@H]([C@H](O1)CO)O)O)O)O)O | C21H20O12 | 464.38 |
| 42 | Isoimperatorin | Isoimperatorin | | 482-45-1 | CC(=CCOc1c2ccc(=O)oc2cc2c1cco2)C | C16H14O4 | 270.28 |
| 43 | Daphnetin | Daphnetin | 7,8-Dihydroxycoumarin;Daphnetol | 486-35-1 | c1cc(c(c2c1ccc(=O)o2)O)O | C9H6O4 | 178.14 |
| 44 | Curcumol | Curcumol | (-)-Curcumol | 4871-97-0 | C1C[C@@H]([C@@]23C[C@H]([C@@](CC(=C)[C@H]12)(O3)O)C(C)C)C | C15H24O2 | 236.35 |

| | | | | | | | |
|---|---|---|---|---|---|---|---|
| 45 | Forsythin | Forsythin | Phillyrin | 487-41-2 | [C@@H]1([C@@H]([C@H]([C@@H]([C@H](O1)CO)O)O)O)Oc1c(cc(cc1)[C@H]1OC[C@H]2[C@@H]1CO[C@H]2c1cc(c(cc1)OC)OC)OC | C27H34O11 | 534.56 |
| 46 | Icariin | Icariin |Ieariline | 489-32-7 | COc1ccc(cc1)-c1oc2c(C\C=C(\C)C)c(O[C@@H]3O[C@H](CO)[C@@H](O)[C@H](O)[C@H]3O)cc(O)c2c(=O)c1O[C@@H]1O[C@@H](C)[C@H](O)[C@@H](O)[C@H]1O | C33H40O15 | 676.68 |
| 47 | Epicatechin | Epicatechin | (-)-Epicatechin;(-)-Epicatechol;L-Epicatechin | 490-46-0 | C1[C@H]([C@H](Oc2cc(cc(c12)O)O)c1cc(c(cc1)O)O)O | C15H14O6 | 290.28 |
| 48 | Gentisic acid | Gentisic acid | 2,5-Dihydroxybenzoic acid;Phloretate;5-Hydroxysalicylic acid;Gentianic acid;Carboxyhydroquinone;Hydroquinonecarboxylic acid | 490-79-9 | c1cc(c(cc1O)C(=O)O)O | C7H6O4 | 154.12 |
| 49 | Salvianolic acid A | Salvianolic acid A | Dan Phenolic Acid A | 96574-01-5 | OC(=O)[C@@H](Cc1ccc(O)c(O)c1)OC(=O)\C=C\c1ccc(O)c(O)c1\C=C\c1ccc(O)c(O)c1 | C26H22O10 | 494.45 |

| | | | | | | | |
|---|---|---|---|---|---|---|---|
| 50 | Salvianolic acid B | Salvianolic acid B | Dan Phenolic Acid B | 115939-25-8 | OC(=O)[C@@H](Cc1ccc(O)c(O)c1)OC(=O)\C=C\c1ccc(O)c2O[C@@H]([C@@H](C(=O)O[C@H](Cc3ccc(O)c(O)c3)C(O)=O)c12)c1ccc(O)c(O)c1 | C36H30O16 | 718.59 |
| 51 | Salvianolic acid C | Salvianolic Acid C | Dan Phenolic Acid C | 115841-09-3 | OC(=O)[C@@H](Cc1ccc | C26H20O10 | 492.43 |
| 52 | Salvianolic acid D | Salvianolic acid D | Dan Phenolic Acid D | 142998-47-8 | OC(=O)Cc1c(O)c(O)ccc1 | C20H18O10 | 418.35 |
| 53 | Salvianolic acid E | Salvianolic acid E | Dan Phenolic Acid E | 142998-46-7 | OC(=O)Cc1c(O)c(O)ccc1\C=C\C(=O)O[C@H](Cc1ccc(O)c(O)c1)C(O)=O1OC(=O)Cc1c(O)c(O)ccc1\C=C\C(=O)O[C@H](Cc1ccc(O)c(O)c1)C(O)=O<br>OC(=O)Cc1c(O)c(O)ccc1\C=C\C(=O)O[C@H](Cc1ccc(O)c(O)c1)C(O)=O1 | C36H30O16 | 718.61 |
| 54 | Salvianolic acid F | Salvianolic acid F | Dan Phenolic Acid F | 158732-59-3 | OC1=C(C=CC(/C=C/C(O | C17H14O6 | 314.29 |

| | | | | | | | |
|---|---|---|---|---|---|---|---|
| 55 | Salvianolic acid Y | Salvianolic acid Y | Dan Phenolic Acid Y | 1638738-76-7 | O=C(O)[C@@H](OC([C@ | $C_{36}H_{30}O_{16}$ | 718.61 |

Abbreviations: SMILES, simplified molecular input line entry system; CAS, chemical abstract service.

**Supplementary Table 2. Significance analysis related to data in Fig. 2c, Extended Data Fig. 1c, Extended Data Fig. 2c and Supplementary Fig. 2a,c,e,g,i,k.**

***SAA-related dataset***

| Student's *t*-test (unpaired two-tailed) | Summary | Adjusted *P* Value |
|---|---|---|
| SAA - 0 µM | | |
| CTRL *vs.* Bleomycin-induced senescence (TIS) | ^ | 0.3218 |
| CTRL *vs.* Replication-induced senescence (RS) | ^ | 0.6873 |
| CTRL *vs.* Oncogene-induced senescence (OIS) | ^ | 0.4380 |
| SAA - 25 µM | | |
| CTRL *vs.* Bleomycin-induced senescence (TIS) | ^ | 0.1345 |
| CTRL *vs.* Replication-induced senescence (RS) | ^ | 0.5195 |
| CTRL *vs.* Oncogene-induced senescence (OIS) | ^ | 0.7290 |
| SAA - 50 µM | | |
| CTRL *vs.* Bleomycin-induced senescence (TIS) | ^ | 0.0522 |
| CTRL *vs.* Replication-induced senescence (RS) | ^ | 0.5631 |
| CTRL *vs.* Oncogene-induced senescence (OIS) | ^ | 0.3560 |
| SAA - 100 µM | | |
| CTRL *vs.* Bleomycin-induced senescence (TIS) | * | 0.0121 |
| CTRL *vs.* Replication-induced senescence (RS) | ** | 0.0041 |
| CTRL *vs.* Oncogene-induced senescence (OIS) | *** | 0.0003 |
| SAA - 200 µM | | |
| CTRL *vs.* Bleomycin-induced senescence (TIS) | *** | 0.0004 |
| CTRL *vs.* Replication-induced senescence (RS) | *** | 0.0002 |
| CTRL *vs.* Oncogene-induced senescence (OIS) | **** | < 0.0001 |
| SAA - 300 µM | | |
| CTRL *vs.* Bleomycin-induced senescence (TIS) | **** | < 0.0001 |
| CTRL *vs.* Replication-induced senescence (RS) | **** | < 0.0001 |
| CTRL *vs.* Oncogene-induced senescence (OIS) | **** | < 0.0001 |
| SAA - 400 µM | | |
| CTRL *vs.* Bleomycin-induced senescence (TIS) | **** | < 0.0001 |
| CTRL *vs.* Replication-induced senescence (RS) | **** | < 0.0001 |
| CTRL *vs.* Oncogene-induced senescence (OIS) | **** | < 0.0001 |
| SAA - 600 µM | | |
| CTRL *vs.* Bleomycin-induced senescence (TIS) | **** | < 0.0001 |
| CTRL *vs.* Replication-induced senescence (RS) | **** | < 0.0001 |
| CTRL *vs.* Oncogene-induced senescence (OIS) | **** | < 0.0001 |

***SAB-related dataset***

| Student's *t*-test (unpaired two-tailed) | Summary | Adjusted *P* Value |
|---|---|---|
| SAB - 0 µM | | |
| CTRL *vs.* Bleomycin-induced senescence (TIS) | ^ | 0.6559 |
| CTRL *vs.* Replication-induced senescence (RS) | ^ | 0.9375 |
| CTRL *vs.* Oncogene-induced senescence (OIS) | ^ | 0.9529 |
| SAB - 25 µM | | |
| CTRL *vs.* Bleomycin-induced senescence (TIS) | ^ | 0.5313 |
| CTRL *vs.* Replication-induced senescence (RS) | ^ | 0.7679 |
| CTRL *vs.* Oncogene-induced senescence (OIS) | ^ | 0.4971 |
| SAB - 50 µM | | |
| CTRL *vs.* Bleomycin-induced senescence (TIS) | ^ | 0.0621 |
| CTRL *vs.* Replication-induced senescence (RS) | ^ | 0.5432 |
| CTRL *vs.* Oncogene-induced senescence (OIS) | ^ | 0.6250 |
| SAB - 100 µM | | |
| CTRL *vs.* Bleomycin-induced senescence (TIS) | * | 0.0157 |
| CTRL *vs.* Replication-induced senescence (RS) | * | 0.0116 |
| CTRL *vs.* Oncogene-induced senescence (OIS) | *** | 0.0007 |
| SAB - 200 µM | | |
| CTRL *vs.* Bleomycin-induced senescence (TIS) | *** | 0.0002 |

| | | |
|---|---|---|
| CTRL *vs.* Replication-induced senescence (RS) | **** | < 0.0001 |
| CTRL *vs.* Oncogene-induced senescence (OIS) | **** | < 0.0001 |
| SAB - 300 µM | | |
| CTRL *vs.* Bleomycin-induced senescence (TIS) | **** | < 0.0001 |
| CTRL *vs.* Replication-induced senescence (RS) | **** | < 0.0001 |
| CTRL *vs.* Oncogene-induced senescence (OIS) | **** | < 0.0001 |
| SAB - 400 µM | | |
| CTRL *vs.* Bleomycin-induced senescence (TIS) | **** | < 0.0001 |
| CTRL *vs.* Replication-induced senescence (RS) | **** | < 0.0001 |
| CTRL *vs.* Oncogene-induced senescence (OIS) | **** | < 0.0001 |
| SAB - 600 µM | | |
| CTRL *vs.* Bleomycin-induced senescence (TIS) | **** | < 0.0001 |
| CTRL *vs.* Replication-induced senescence (RS) | **** | < 0.0001 |
| CTRL *vs.* Oncogene-induced senescence (OIS) | **** | < 0.0001 |

***SAE-related dataset***

| **Student's *t*-test (unpaired two-tailed)** | **Summary** | **Adjusted *P* Value** |
|---|---|---|
| SAE - 0 µM | | |
| CTRL *vs.* Bleomycin-induced senescence (TIS) | ^ | 0.3874 |
| CTRL *vs.* Replication-induced senescence (RS) | ^ | 0.9141 |
| CTRL *vs.* Oncogene-induced senescence (OIS) | ^ | 0.9182 |
| SAE - 5 µM | | |
| CTRL *vs.* Bleomycin-induced senescence (TIS) | ^ | 0.4587 |
| CTRL *vs.* Replication-induced senescence (RS) | ^ | 0.9036 |
| CTRL *vs.* Oncogene-induced senescence (OIS) | ^ | 0.7831 |
| SAE - 10 µM | | |
| CTRL *vs.* Bleomycin-induced senescence (TIS) | ^ | 0.0723 |
| CTRL *vs.* Replication-induced senescence (RS) | ^ | 0.7073 |
| CTRL *vs.* Oncogene-induced senescence (OIS) | ^ | 0.7607 |
| SAE - 50 µM | | |
| CTRL *vs.* Bleomycin-induced senescence (TIS) | ** | 0.0016 |
| CTRL *vs.* Replication-induced senescence (RS) | *** | 0.0002 |
| CTRL *vs.* Oncogene-induced senescence (OIS) | *** | 0.0002 |
| SAE - 100 µM | | |
| CTRL *vs.* Bleomycin-induced senescence (TIS) | **** | < 0.0001 |
| CTRL *vs.* Replication-induced senescence (RS) | **** | < 0.0001 |
| CTRL *vs.* Oncogene-induced senescence (OIS) | **** | < 0.0001 |
| SAE - 200 µM | | |
| CTRL *vs.* Bleomycin-induced senescence (TIS) | **** | < 0.0001 |
| CTRL *vs.* Replication-induced senescence (RS) | **** | < 0.0001 |
| CTRL *vs.* Oncogene-induced senescence (OIS) | **** | < 0.0001 |
| SAE - 300 µM | | |
| CTRL *vs.* Bleomycin-induced senescence (TIS) | **** | < 0.0001 |
| CTRL *vs.* Replication-induced senescence (RS) | **** | < 0.0001 |
| CTRL *vs.* Oncogene-induced senescence (OIS) | **** | < 0.0001 |

**Supplementary Table 3. Significance analysis by Student's *t*-test related to data of Supplementary Fig. 3g-i.**

**Supplementary Table 3. IMR90 + SAE**

| | |
|---|---|
| Student's *t*-test | Ordinary |
| Alpha | 0.05 |

| Tukey's multiple comparisons test | Summary | Adjusted P Value |
|---|---|---|
| PCC1 - 0 µM | | |
| CTRL *vs.* TIS | ^ | > 0.9999 |
| CTRL *vs.* RS | ^ | > 0.9999 |
| CTRL *vs.* OIS | ^ | > 0.9999 |
| PCC1 - 10 µM | | |
| CTRL *vs.* TIS | ^ | 0.8820 |
| CTRL *vs.* RS | ^ | > 0.9999 |
| CTRL *vs.* OIS | ^ | 0.7287 |
| PCC1 - 25 µM | | |
| CTRL *vs.* TIS | ^ | 0.5898 |
| CTRL *vs.* RS | ^ | 0.2980 |
| CTRL *vs.* OIS | ^ | 0.1647 |
| PCC1 - 50 µM | | |
| CTRL *vs.* TIS | * | 0.0428 |
| CTRL *vs.* RS | * | 0.0426 |
| CTRL *vs.* OIS | * | 0.0235 |
| PCC1 - 75 µM | | |
| CTRL *vs.* TIS | ** | 0.0056 |
| CTRL *vs.* RS | ** | 0.0099 |
| CTRL *vs.* OIS | ** | 0.0033 |
| PCC1 - 100 µM | | |
| CTRL *vs.* TIS | *** | 0.0004 |
| CTRL *vs.* RS | ** | 0.0018 |
| CTRL *vs.* OIS | *** | 0.0007 |
| PCC1 - 125 µM | | |
| CTRL *vs.* TIS | *** | 0.0004 |
| CTRL *vs.* RS | **** | < 0.0001 |
| CTRL *vs.* OIS | ** | 0.0011 |
| PCC1 - 150 µM | | |
| CTRL *vs.* TIS | **** | < 0.0001 |
| CTRL *vs.* RS | **** | < 0.0001 |
| CTRL *vs.* OIS | **** | < 0.0001 |
| PCC1 - 200 µM | | |
| CTRL *vs.* TIS | **** | < 0.0001 |
| CTRL *vs.* RS | **** | < 0.0001 |
| CTRL *vs.* OIS | **** | < 0.0001 |

**Supplementary Table 3. HUVEC + SAE**

| | |
|---|---|
| Student's *t*-test | Ordinary |
| Alpha | 0.05 |

| Tukey's multiple comparisons test | Summary | Adjusted P Value |
|---|---|---|
| PCC1 - 0 µM | | |
| CTRL *vs.* TIS | ^ | > 0.9999 |
| CTRL *vs.* RS | ^ | > 0.9999 |
| CTRL *vs.* OIS | ^ | > 0.9999 |
| PCC1 - 10 µM | | |
| CTRL *vs.* TIS | ^ | 0.8944 |
| CTRL *vs.* RS | ^ | 0.6240 |
| CTRL *vs.* OIS | ^ | 0.4216 |

| | | |
|---|---|---|
| PCC1 - 25 µM | | |
| CTRL *vs.* TIS | ^ | 0.0913 |
| CTRL *vs.* RS | ^ | 0.2184 |
| CTRL *vs.* OIS | ^ | 0.1321 |
| PCC1 - 50 µM | | |
| CTRL *vs.* TIS | * | 0.0423 |
| CTRL *vs.* RS | * | 0.0346 |
| CTRL *vs.* OIS | * | 0.0448 |
| PCC1 - 75 µM | | |
| CTRL *vs.* TIS | ** | 0.0024 |
| CTRL *vs.* RS | ** | 0.0010 |
| CTRL *vs.* OIS | ** | 0.0020 |
| PCC1 - 100 µM | | |
| CTRL *vs.* TIS | *** | 0.0004 |
| CTRL *vs.* RS | *** | 0.0001 |
| CTRL *vs.* OIS | *** | 0.0004 |
| PCC1 - 125 µM | | |
| CTRL *vs.* TIS | **** | < 0.0001 |
| CTRL *vs.* RS | **** | < 0.0001 |
| CTRL *vs.* OIS | **** | < 0.0001 |
| PCC1 - 150 µM | | |
| CTRL *vs.* TIS | **** | < 0.0001 |
| CTRL *vs.* RS | **** | < 0.0001 |
| CTRL *vs.* OIS | **** | < 0.0001 |
| PCC1 - 200 µM | | |
| CTRL *vs.* TIS | **** | < 0.0001 |
| CTRL *vs.* RS | **** | < 0.0001 |
| CTRL *vs.* OIS | **** | < 0.0001 |

**Supplementary Table 3. AT-MSC + SAE**

| | |
|---|---|
| Student's *t* -test | Ordinary |
| Alpha | 0.05 |

| Tukey's multiple comparisons test | Summary | Adjusted P Value |
|---|---|---|
| PCC1 - 0 µM | | |
| CTRL *vs.* TIS | ^ | > 0.9999 |
| CTRL *vs.* RS | ^ | > 0.9999 |
| CTRL *vs.* OIS | ^ | > 0.9999 |
| PCC1 - 10 µM | | |
| CTRL vs. TIS | ^ | 0.8820 |
| CTRL vs. RS | ^ | 0.5614 |
| CTRL vs. OIS | ^ | 0.2595 |
| PCC1 - 25 µM | | |
| CTRL *vs.* TIS | ^ | 0.0686 |
| CTRL *vs.* RS | ^ | 0.1145 |
| CTRL *vs.* OIS | ^ | 0.0734 |
| PCC1 - 50 µM | | |
| CTRL *vs.* TIS | * | 0.0113 |
| CTRL *vs.* RS | * | 0.0191 |
| CTRL *vs.* OIS | ** | 0.0037 |
| PCC1 - 75 µM | | |
| CTRL *vs.* TIS | ** | 0.0030 |
| CTRL *vs.* RS | ** | 0.0035 |
| CTRL *vs.* OIS | ** | 0.0036 |
| PCC1 - 100 µM | | |
| CTRL *vs.* TIS | **** | < 0.0001 |
| CTRL *vs.* RS | **** | < 0.0001 |
| CTRL *vs.* OIS | **** | < 0.0001 |

| | | |
|---|---|---|
| PCC1 - 125 µM | | |
| CTRL *vs.* TIS | **** | < 0.0001 |
| CTRL *vs.* RS | **** | < 0.0001 |
| CTRL *vs.* OIS | **** | < 0.0001 |
| PCC1 - 150 µM | | |
| CTRL *vs.* TIS | **** | < 0.0001 |
| CTRL *vs.* RS | **** | < 0.0001 |
| CTRL *vs.* OIS | **** | < 0.0001 |
| PCC1 - 200 µM | | |
| CTRL *vs.* TIS | **** | < 0.0001 |
| CTRL *vs.* RS | **** | < 0.0001 |
| CTRL *vs.* OIS | **** | < 0.0001 |

**Supplementary Table 4. Protein targets identified by 20K arrays**

| SAA-Bio | SAB-Bio | SAE-Bio | Co-identified targets |
|---|---|---|---|
| CKS2 | COL4A3BP | GAP43 | GAP43 |
| GAP43 | GAP43 | GUK1 | AK6 |
| GTF2I | GTF2I | NME2 | TK1 |
| GUK1 | PICK1 | AK6 | GKAP1 |
| KHK | PRMT2 | TK1 | PIP4K2C |
| NME1 | AK6 | CMPK1 | ULK2 |
| PICK1 | TK1 | GKAP1 | CLIC2 |
| SGK1 | GKAP1 | MAPK12 | GNAI2 |
| SYTL2 | PIP4K2C | PIP4K2C | PRPSAP1 |
| AK6 | ULK2 | ULK2 | RAB14 |
| TK1 | ARL2 | ATP6V1C1 | RAB3B |
| CMPK1 | CLIC2 | CLIC2 | PLEKHJ1 |
| GKAP1 | GNAI2 | GNAI2 | RTN4 |
| MAPK12 | PRPSAP1 | PRPSAP1 | SAT1 |
| PIP4K2C | RAB14 | RAB14 | SFN |
| ULK2 | RAB18 | RAB3B | SOD1 |
| ARL2 | RAB3B | EMC9 | GGCT |
| ATP6V1C1 | RAB9A | NTMT1 | GLOD4 |
| CLIC2 | RCVRN | PLEKHJ1 | NDUFA8 |
| GNAI2 | SAR1B | RTN4 | RPLP1 |
| PRPSAP1 | HN1 | SAT1 | ARL5B |
| RAB14 | PLEKHJ1 | SFN | CRABP2 |
| RAB18 | RTN4 | SOD1 | PHPT1 |
| RAB3B | SAT1 | GGCT | CDC34 |
| RAB9A | SOD1 | GLOD4 | EMCN |
| RRAS2 | GGCT | NDUFA8 | USP15 |
| SAR1B | GLOD4 | RPLP1 | NEIL2 |
| EMC9 | NDUFA8 | ARL5B | HPGDS |
| NTMT1 | RPLP1 | CDC37L1 | DTNBP1 |
| HN1 | ARL5B | CRABP2 | AZIN1 |
| IFIT3 | CRABP2 | PHPT1 | C11orf49 |
| IP6K1 | PHPT1 | SAAL1 | GSTP1 |
| PANK3 | S100A6 | CALB1 | MB |
| PHF23 | SAAL1 | CDC34 | SCG3 |
| PLEKHJ1 | ARL2BP | EMCN | AP1AR |
| RTN4 | CALB1 | CYTH4_frag | METTL21B |
| S100A11 | CDC34 | SELENBP1 | FAM131C |
| SAT1 | EMCN | USP15 | LYPLA2 |
| SFN | USP15 | NEIL2 | RBBP9 |
| SOD1 | BHMT2 | HPGDS | TSPAN1 |
| GGCT | NEIL2 | GTF3C6 | WIPI2 |
| DYNLT3 | HPGDS | DTNBP1 | APIP |
| GAPDH | GTF3C6 | PPCDC | ASNS |
| GLOD4 | DTNBP1 | AZIN1 | BLVRA |
| LGALS1 | AZIN1 | GSTP1 | BTF3 |
| NDUFA8 | BBOX1 | MB | SMAP |
| RPLP1 | DCPS | SCG3 | MAD2L1 |
| ARL5B | GSTP1 | AP1AR | PRDX3 |
| CDC37L1 | MB | CMBL | TBCB |
| CRABP2 | SCG3 | METTL21B | ETS1 |
| PHPT1 | AP1AR | FAM131C | TSR2 |
| S100A6 | METTL21B | OLA1 | EIF4EBP3 |
| CDC34 | FAM131C | MAGEA4 | HBG1 |
| EMCN | OLA1 | LYPLA2 | HBG2 |
| CYTH4_frag | HNRNPA0 | GLO1 | HRSP12 |

| | | | |
|---|---|---|---|
| USP15 | MAGEA4 | NOSIP | ISG15 |
| NEIL2 | ECE2 | RBBP9 | CA1 |
| HPGDS | LYPLA2 | TSPAN1 | GRAP2 |
| DTNBP1 | RBBP9 | WIPI2 | NOTCH2NL |
| PPCDC | TSPAN1 | ZG16B | ARG1 |
| AZIN1 | WIPI2 | APIP | GIMAP7 |
| BBOX1 | ZG16B | ASNS | 45353 |
| C11orf49 | APIP | BLVRA | COTL1 |
| DCPS | ASNS | BTF3 | HSPBAP1 |
| GSTP1 | BCL2L13 | SMAP | PLEK |
| MB | BLVRA | MAD2L1 | SNAPC5 |
| MRPL44 | BTF3 | PBLD | DEF6_frag |
| SCG3 | SMAP | PRDX3 | TGM4 |
| AP1AR | FXYD5 | TBCB | ARPC1B |
| METTL21B | MAD2L1 | GDPD5 | SHD |
| FAM131C | PRDX3 | PIM1 | AAR2 |
| HNRNPA0 | TBCB | ETS1 | ECHDC1 |
| ECE2 | ETS1 | TSR2 | HM13 |
| LYPLA2 | HAUS8 | EIF4EBP3 | ANXA8 |
| GLO1 | TSR2 | HBG1 | HACL1 |
| NOSIP | AIF1 | HBG2 | ID3 |
| RBBP9 | C1orf21 | HRSP12 | METTL9 |
| TSPAN1 | EIF4EBP3 | ISG15 | NUDT2 |
| WIPI2 | HBG1 | MCM7 | PGM3 |
| APIP | HBG2 | CA1 | ARRB1 |
| ASNS | HRSP12 | GRAP2 | C6orf106 |
| BLVRA | IL32 | NOTCH2NL | C9orf16 |
| BTF3 | ISG15 | ARG1 | TRIAP1 |
| SMAP | SKP1 | DYDC1 | XM_002816682.3_frag |
| CMSS1 | CA1 | GIMAP7 | GINS2 |
| FXYD5 | GRAP2 | MAGEB2 | GPBP1_frag |
| MAD2L1 | IDO1 | 2-Mar | RAD1 |
| PBLD | NOTCH2NL | COTL1 | SPG21 |
| PRDX3 | ARG1 | HSPBAP1 | MYL6 |
| SNX3 | GIMAP7 | PLEK | KXD1 |
| TBCB | 2-Mar | SNAPC5 | HSBP1 |
| GDPD5 | NDUFA3 | CRYAB | UBQLN4_frag |
| ETS1 | SMCP | DEF6_frag | ILF2 |
| HPGD | COTL1 | PDCD2L | PSMD4 |
| HAUS8 | HSPBAP1 | TGM4 | S100A8 |
| RAB24 | PLEK | ARPC1B | OLA1 |
| TSR2 | SNAPC5 | CCT3 | KJ903456_frag |
| EIF4EBP3 | CRYAB | SHD | DACT3 |
| HBG1 | DEF6_frag | AAR2 | PVRL3_frag |
| HBG2 | PDCD2L | CALCOCO2 | NUDCD2 |
| HRSP12 | TGM4 | ECHDC1 | SCGB3A2 |
| IL32 | ANP32A | HM13 | LYPLAL1 |
| ISG15 | ARPC1B | ANXA8 | ANAPC15 |
| MCM7 | CAPRIN2_frag | HACL1 | S100B |
| SETMAR | CCT3 | ID3 | TMSB10 |
| C11orf16 | PNMA6A | METTL9 | LRAT |
| CA1 | SHD | CPTP_frag | NME5 |
| GRAP2 | TUBB2B | NUDT2 | ARL1 |
| IDO1 | AAR2 | PGM3 | ZNF471_frag |
| NOTCH2NL | ECHDC1 | TSTA3 | ATG4C |
| ARG1 | HM13 | ARRB1 | BC047307_frag |
| DYDC1 | LCP1 | C11orf49 | COMT |
| GIMAP7 | ANXA8 | C6orf106 | DAPP1 |
| MAGEB2 | ADIRF | C9orf16 | CCDC53 |

| | | | |
|---|---|---|---|
| 2-Mar | HACL1 | TRIAP1 | C3orf52 |
| SMCP | ID3 | XM_002816682.3_frag | NANP |
| COTL1 | METTL9 | GINS2 | TXNDC17 |
| HSPBAP1 | NUDT2 | GPBP1_frag | ATG12 |
| PLEK | PGM3 | RAD1 | CSTA |
| SNAPC5 | ARRB1 | SPG21 | FNDC3B |
| DEF6_frag | C11orf49 | MYL6 | S100P |
| FABP4 | C6orf106 | KXD1 | BC033035.2_frag |
| TGM4 | C9orf16 | HSBP1 | URM1 |
| ARPC1B | CA3 | UBQLN4_frag | C15orf41 |
| CAPRIN2_frag | DHFR | ILF2 | DEPP |
| MAN1B1 | PLP1 | PSMD4 | CBX1 |
| NABP2 | THAP4 | S100A8 | GUCA1A |
| PNMA6A | TRIAP1 | ZCCHC7 | NDUFAF2 |
| SHD | UBE2R2 | TRMT12 | SDSL |
| AAR2 | XM_002816682.3_frag | IFIT5 | C4orf19 |
| ECHDC1 | GINS2 | KJ903456_frag | FTSJ1 |
| HM13 | GPBP1_frag | DACT3 | CD53 |
| LCP1 | NASP | KLHL3_frag | REEP6 |
| ZCCHC17 | RAD1 | PVRL3_frag | GDA |
| ANXA8 | RUVBL2 | C1QTNF7 | UBE2H |
| ADIRF | SERPINB6 | NUDCD2 | THOC1 |
| HACL1 | SPG21 | SCGB3A2 | ABTB1 |
| ID3 | DBNDD1 | TTR | IGFBP6 |
| METTL9 | MYL6 | DPY30 | STARD7 |
| CPTP_frag | PEX19 | LYPLAL1 | XAGE3 |
| MPST | KXD1 | ANAPC15 | BMF |
| NUDT2 | HSBP1 | S100B | THEM4 |
| PGM3 | SNRPF | TMSB10 | EXOSC7 |
| PSAT1 | SNX1 | GID8 | DMRTC1 |
| SNCB | UBQLN4_frag | FAM131A | FAM84A |
| ARRB1 | ILF2 | LRAT | RNH1 |
| C6orf106 | PSMD4 | NME5 | L3HYPDH |
| C9orf16 | S100A8 | BCL2A1 | CYP4F11 |
| CA3 | ZCCHC7 | TEX2_frag | MIEN1 |
| IFT20 | NFKBIB | ARL1 | PITPNA |
| RNF181 | KJ903456_frag | LGALS2 | TPM1 |
| DHFR | DACT3 | THRSP | ALOXE3 |
| THAP4 | KLHL3_frag | LSM8 | IL18 |
| TRIAP1 | PLEK2 | SHFM1 | CREG1 |
| XM_002816682.3_frag | PVRL3_frag | ZNF471_frag | |
| DCXR | NUDCD2 | ATG4C | |
| GINS2 | SCGB3A2 | BC047307_frag | |
| GPBP1_frag | TTR | PAPSS1 | |
| NXT1 | CFP | FGFR1OP | |
| RAD1 | LYPLAL1 | COMT | |
| SERPINB6 | ANAPC15 | DAPP1 | |
| SPG21 | FANCI_frag | CCDC53 | |
| DBNDD1 | S100B | HMGB2 | |
| IST1 | TMSB10 | PPP1R2 | |
| MYL6 | CLDN3 | UCHL1 | |
| KXD1 | FAM131A | C3orf52 | |
| HSBP1 | LRAT | NANP | |
| UBQLN4_frag | NME5 | PHLDA3 | |
| ILF2 | SERPINA1 | TXNDC17 | |
| ODAM_frag | BCL2A1 | ATG12 | |
| PSMD4 | ARHGDIB | CSTA | |
| RPLP2 | ARL1 | FNDC3B | |
| S100A8 | SENP8 | S100P | |

| | | |
|---|---|---|
| OLA1 | THRSP | TIMM13 |
| TRMT12 | SHFM1 | BC033035.2_frag |
| IFIT5 | ZNF471_frag | URM1 |
| KJ903456_frag | ATG4C | PAGE4 |
| DACT3 | BC047307_frag | C15orf41 |
| STAM | COMT | RABGGTB |
| PVRL3_frag | DAPP1 | DEPP |
| S100A3 | HAO2 | CBX1 |
| NUDCD2 | ANXA5 | EFCAB1 |
| SCGB3A2 | CCDC53 | EIF3K |
| CFP | HDGFRP3 | SPAG7 |
| GAGE2D | PPP1R2 | MTUS2 |
| LYPLAL1 | C3orf52 | WDR61 |
| ANAPC15 | CALM1 | GUCA1A |
| PSMG3 | DNAL1 | NDUFAF2 |
| FANCI_frag | NANP | BHMT |
| S100B | TXNDC17 | SCIN |
| TMSB10 | ATG12 | SDSL |
| ATF2 | CSTA | TDO2 |
| CLDN3 | FNDC3B | UROD |
| LRAT | S100P | C4orf19 |
| NME5 | TIMM13 | PRKAR1A |
| TEX2_frag | ARRB2 | IFIH1 |
| ARL1 | BC033035.2_frag | FTSJ1 |
| C12orf10 | INPP1 | MAGEC2 |
| LGALS2 | SDCCAG3 | PKIB |
| MMP28 | URM1 | CD53 |
| SENP8 | SP140L | REEP6 |
| CRABP1 | ANXA10 | AAGAB |
| LSM8 | C15orf41 | SART3 |
| ZNF471_frag | NUDC | STK16 |
| ATG4C | RABGGTB | NDUFAB1 |
| TPD52L1 | DEPP | RABEPK |
| BC047307_frag | CBX1 | RMI1_frag |
| PAPSS1 | KIAA0513 | GDA |
| FGFR1OP | MTUS2 | CNOT7 |
| COMT | CALML5 | GRB2 |
| DAPP1 | PMP2 | UBE2H |
| CCDC53 | SFN | ADSS |
| EIF2S1 | GUCA1A | KJ902886 |
| HDGFRP3 | NDUFAF2 | CMIP |
| HMGB2 | RAB37 | THOC1 |
| C3orf52 | TPT1 | GSKIP |
| NANP | BHMT | ABTB1 |
| TXNDC17 | SDSL | IGFBP6 |
| ATG12 | C4orf19 | DRICH1 |
| CSTA | PRKAR1A | STARD7 |
| FNDC3B | IFIH1 | CHCHD4 |
| S100P | ALOX15B | EXTL3 |
| TPRKB | FTSJ1 | CCNDBP1 |
| ARRB2 | MAGEC2 | PTGES3 |
| BC033035.2_frag | PKIB | XAGE3 |
| SDCCAG3 | CD53 | BMF |
| URM1 | REEP6 | THEM4 |
| SP140L | AAGAB | PCP2 |
| PAGE4 | SART3 | BSND |
| POP7 | RMI1_frag | C1orf174 |
| ANXA10 | GDA | PMAIP1 |
| C15orf41 | PAGE2 | EXOSC7 |

| | | |
|---|---|---|
| NUDC | VCY | KRT18P55 |
| 2-Sep | TNIK_frag | ATG4A |
| DEPP | SESTD1 | PTMS |
| CBX1 | UBE2H | UBA3 |
| IL6 | S100A1 | DMRTC1 |
| MARVELD2 | ADSS | HBM |
| GUCA1A | SYAP1 | S100A2 |
| NDUFAF2 | KJ902886 | FAM84A |
| RAB37 | CMIP | TMSB4X |
| SDSL | UBALD1 | TRAPPC3 |
| TDO2 | FBXO2 | N6AMT2 |
| UROD | SERPINB4 | RNH1 |
| C4orf19 | RGS10 | TRIM13 |
| CROCCP2 | THOC1 | L3HYPDH |
| ALOX15B | GSKIP | CYP4F11 |
| FTSJ1 | ABTB1 | MIEN1 |
| CD53 | IGFBP6 | MAPK8 |
| REEP6 | PRUNE | PITPNA |
| STK16 | DRICH1 | TCL6 |
| NDUFAB1 | STARD7 | TPM1 |
| GDA | CHCHD4 | SET |
| DCUN1D2 | EXTL3 | THOC7 |
| HTATIP2 | GAB1 | MGEA5 |
| RAB1B | XAGE3 | TYW3 |
| UBE2H | BMF | LETM2 |
| AIPL1 | THEM4 | ALOXE3 |
| SAR1A | OCM | HNRNPC |
| SYAP1 | ALOX12B | PVALB |
| DRAP1 | EXOSC7 | ATXN10 |
| SERPINB4 | IFNA6 | POLR3H |
| THOC1 | NACA | IL18 |
| ABTB1 | ANKRD26P1 | CREG1 |
| ACTN1 | DMRTC1 | PRMT3 |
| IGFBP6 | SH3BGR | |
| STARD7 | FAM84A | |
| RLBP1 | MRI1 | |
| PTGES3 | RNH1 | |
| RBP2 | GLMN | |
| XAGE3 | L3HYPDH | |
| BMF | CYP4F11 | |
| THEM4 | MIEN1 | |
| PCP2 | PITPNA | |
| C1orf174 | TCL6 | |
| PMAIP1 | TPM1 | |
| EXOSC7 | SET | |
| ATG4A | THOC7 | |
| CSK | MUM1L1 | |
| PTMS | ALOXE3 | |
| TSR1 | ATXN10 | |
| ANKRD26P1 | POLR3H | |
| DMRTC1 | ENDOU | |
| HBM | IL18 | |
| S100A2 | GCSH | |
| FAM84A | CREG1 | |
| G6PD | | |
| MRI1 | | |
| RNH1 | | |
| PDE9A | | |
| TRIM13 | | |

| |
|---|
| L3HYPDH |
| CYP4F11 |
| MIEN1 |
| MAPK8 |
| PITPNA |
| RMND5B |
| TPM1 |
| PDE6D |
| GPN1 |
| ZCRB1 |
| NUDT10 |
| TYW3 |
| LETM2 |
| IFNA14 |
| IFNA4 |
| ALOXE3 |
| ME3 |
| DEFA1 |
| ENDOU |
| IL18 |
| C14orf166 |
| CREG1 |
| TUBGCP4 |

Note: GSTP1 is hypothesized to be a putative co-target that mediates the senolytic activity of SA-compounds.

**Supplementary Table 5. Pathway and process enrichment analysis outputs**

| GO | Category | Description | Count | Percentage (%) | Log10(P) | Log10(q) |
|---|---|---|---|---|---|---|
| GO:0098869 | GO Biological Processes | Cellular oxidant detoxification | 8 | 5.56 | -7.89 | -3.55 |
| GO:0044248 | GO Biological Processes | Cellular catabolic process | 18 | 12.5 | -7.37 | -3.33 |
| GO:0019752 | GO Biological Processes | Carboxylic acid metabolic process | 17 | 11.81 | -6.62 | -2.97 |
| GO:0044092 | GO Biological Processes | Negative regulation of molecular function | 14 | 9.72 | -4.95 | -1.56 |
| GO:0002832 | GO Biological Processes | Negative regulation of response to biotic stimulus | 6 | 4.17 | -4.54 | -1.31 |
| GO:0009066 | GO Biological Processes | Aspartate family amino acid metabolic process | 4 | 2.78 | -4.21 | -1.16 |
| GO:0032930 | GO Biological Processes | Positive regulation of superoxide anion generation | 3 | 2.08 | -4.09 | -1.15 |
| GO:0055086 | GO Biological Processes | Nucleobase-containing small molecule metabolic process | 11 | 7.64 | -3.84 | -0.96 |
| GO:0032535 | GO Biological Processes | Regulation of cellular component size | 8 | 5.56 | -3.51 | -0.78 |
| GO:0110053 | GO Biological Processes | Regulation of actin filament organization | 7 | 4.86 | -3.47 | -0.75 |
| GO:0009123 | GO Biological Processes | Nucleoside monophosphate metabolic process | 4 | 2.78 | -3.32 | -0.66 |
| R-HSA-6798695 | Reactome Gene Sets | Neutrophil degranulation | 9 | 6.25 | -3.31 | -0.66 |
| GO:0061436 | GO Biological Processes | Establishment of skin barrier | 3 | 2.08 | -3.25 | -0.63 |
| R-HSA-8950505 | Reactome Gene Sets | Gene and protein expression by JAK-STAT signaling after interleukin-12 stimulation | 3 | 2.08 | -3.11 | -0.53 |
| GO:0009611 | GO Biological Processes | Response to wounding | 8 | 5.56 | -2.96 | -0.44 |
| R-HSA-76002 | Reactome Gene Sets | Platelet activation, signaling and aggregation | 6 | 4.17 | -2.78 | -0.32 |
| hsa00330 | KEGG Pathway | Arginine and proline metabolism | 3 | 2.08 | -2.76 | -0.31 |
| GO:2001242 | GO Biological Processes | Regulation of intrinsic apoptotic signaling pathway | 5 | 3.47 | -2.67 | -0.25 |
| GO:0051339 | GO Biological Processes | Regulation of lyase activity | 3 | 2.08 | -2.57 | -0.21 |
| GO:0098930 | GO Biological Processes | Axonal transport | 3 | 2.08 | -2.49 | -0.16 |

Note: Above are top 20 clusters with their representative enriched terms (one *per* cluster).
"Count" is the number of genes in the user-provided lists with membership in the given ontology term.
"%" is the percentage of all of the user-provided genes that are found in the given ontology term (only input genes with at least one ontology term annotation are included in the calculation).
"Log10(P)" is the p-value in log base 10. "Log10(q)" is the multi-test adjusted p-value in log base 10.

**Supplementary Table 6. Protein targets co-identified by pulldown assays**

| ***No.*** | ***Gene Symbol*** |
|---|---|
| 1 | TRIO |
| 2 | FILIP1L |
| 3 | ANXA2 |
| 4 | APEX1 |
| 5 | HNRNPUL1 |
| 6 | ACO2 |
| 7 | TJP1 |
| 8 | NIBAN2 |
| 9 | EIF2S3 |
| 10 | PPA2 |
| 11 | TXNRD1 |
| 12 | PABPC1 |
| 13 | CAST |
| 14 | FUBP1 |
| 15 | SF3B1 |
| 16 | TWF1 |
| 17 | PAICS |
| 18 | CDC37 |
| 19 | LRPAP1 |
| 20 | ARHGAP17 |
| 21 | DNM1L |
| 22 | HSPA2 |
| 23 | MAPK1 |
| 24 | NT5C2 |
| 25 | SELENBP1 |
| 26 | MAP1A |
| 27 | ERC1 |
| 28 | DDX46 |
| 29 | CAPZA2 |
| 30 | UBE2L3 |
| 31 | KHDRBS1 |
| 32 | FUBP3 |
| 33 | SNX27 |
| 34 | HSPH1 |
| 35 | KYNU |
| 36 | TXLNA |
| 37 | GSTP1 |
| 38 | HDGF |
| 39 | BAG3 |
| 40 | SF3A3 |
| 41 | EIF4B |
| 42 | DNAJB1 |
| 43 | PCYT1A |
| 44 | NPLOC4 |
| 45 | VASP |
| 46 | PARP14 |
| 47 | SEC24D |
| 48 | TRIOBP |
| 49 | ECHS1 |

| | |
|---|---|
| 50 | SVIL |
| 51 | DBNL |
| 52 | INPPL1 |
| 53 | TBK1 |
| 54 | GLRX3 |
| 55 | SPTBN1 |
| 56 | STXBP3 |
| 57 | HNRNPH3 |
| 58 | CKAP5 |
| 59 | TLN2 |
| 60 | TWF2 |
| 61 | TIPRL |
| 62 | MTPN |
| 63 | TACO1 |
| 64 | MCTS1 |
| 65 | SF1 |
| 66 | GSR |
| 67 | SRRT |
| 68 | HNRNPDL |
| 69 | GPX4 |
| 70 | SNRNP70 |
| 71 | DDX42 |
| 72 | AKR7A2 |
| 73 | STAMBPL1 |
| 74 | RANGAP1 |
| 75 | SF3B2 |
| 76 | CHORDC1 |
| 77 | ECPAS |
| 78 | NOMO2 |
| 79 | CCDC50 |
| 80 | COLGALT1 |
| 81 | IGFBP5 |
| 82 | GATD3B |
| 83 | SNRPD1 |
| 84 | PSPC1 |
| 85 | PRPF8 |
| 86 | DAZAP1 |
| 87 | NIT1 |
| 88 | FAM98B |
| 89 | UBE2I |
| 90 | BLVRA |
| 91 | CSK |
| 92 | CARS1 |
| 93 | GSTO1 |
| 94 | SH3GL1 |
| 95 | MAPKAPK2 |
| 96 | ABCF2 |
| 97 | CRKL |
| 98 | CTTNBP2NL |
| 99 | GRHPR |
| 100 | HK2 |

| | |
|---|---|
| 101 | SCRIB |
| 102 | XRN2 |
| 103 | ST13P4 |
| 104 | PDAP1 |
| 105 | SELENOH |
| 106 | AK2 |
| 107 | ATG7 |
| 108 | SEC24A |
| 109 | ACSF2 |
| 110 | AKT1 |
| 111 | SCLY |
| 112 | RNF214 |
| 113 | HMGCL |
| 114 | LAP3 |
| 115 | CHID1 |
| 116 | H2BC21 |
| 117 | AMPD2 |
| 118 | SCP2 |
| 119 | GSK3B |
| 120 | PHLDB1 |
| 121 | COA3 |
| 122 | RBM25 |
| 123 | PPP5C |
| 124 | CIRBP |
| 125 | RBPMS |
| 126 | NMT1 |
| 127 | HNRNPAB |
| 128 | SF3A2 |
| 129 | DENR |
| 130 | PIP4K2B |
| 131 | VPS11 |
| 132 | PPP1R12C |
| 133 | EGLN1 |
| 134 | TSN |
| 135 | TRIM3 |
| 136 | PLCD1 |
| 137 | MT2A |
| 138 | NCKIPSD |
| 139 | ASAP1 |
| 140 | RRAGA |
| 141 | TXNDC12 |
| 142 | FYCO1 |
| 143 | CPSF7 |
| 144 | RCC2 |
| 145 | SEH1L |
| 146 | CAMSAP2 |
| 147 | THYN1 |
| 148 | ELAVL1 |
| 149 | BTF3 |
| 150 | EWSR1 |
| 151 | RASA1 |

| | |
|---|---|
| 152 | EIF4H |
| 153 | EFTUD2 |
| 154 | SF3B4 |
| 155 | SF3B3 |
| 156 | RAB11FIP5 |
| 157 | STK24 |
| 158 | HBS1L |
| 159 | DRG1 |
| 160 | MINK1 |
| 161 | CLASP1 |
| 162 | ATXN2 |
| 163 | DDX23 |
| 164 | HOOK3 |
| 165 | EIF2S2 |
| 166 | ACTR10 |
| 167 | OLA1 |
| 168 | BPNT1 |
| 169 | PLPBP |
| 170 | CMPK1 |
| 171 | EXOC6B |
| 172 | C3 |
| 173 | SMAP |
| 174 | IVD |
| 175 | NDUFS5 |
| 176 | CHMP1B |
| 177 | ARIH1 |
| 178 | PRPF6 |
| 179 | ATXN2L |
| 180 | ATAD3A |
| 181 | HSPB8 |
| 182 | EEFSEC |
| 183 | CDC42BPA |
| 184 | RNF213 |
| 185 | CCDC6 |
| 186 | MAP2K1 |
| 187 | SEC23B |
| 188 | GLRX |
| 189 | RBM3 |
| 190 | TIMP1 |
| 191 | PAFAH1B2 |
| 192 | H2AZ2 |
| 193 | MXRA8 |
| 194 | CARS2 |
| 195 | FAM98A |
| 196 | IGFBP7 |
| 197 | HPCAL1 |
| 198 | MTSS2 |
| 199 | FUS |
| 200 | STC1 |
| 201 | RPL26 |
| 202 | NUMB |

| | |
|---|---|
| 203 | ATP5IF1 |
| 204 | TMEM132A |
| 205 | MAPRE2 |
| 206 | AIDA |
| 207 | PDLIM2 |
| 208 | TBCD |
| 209 | IRF2BP2 |
| 210 | ABHD12 |
| 211 | HOMER3 |
| 212 | EIF1 |
| 213 | NUBP2 |
| 214 | USP28 |
| 215 | IGFBP4 |
| 216 | VBP1 |
| 217 | ZFP36L1 |
| 218 | WDR26 |
| 219 | OGFR |
| 220 | SBF1 |
| 221 | IKBKG |
| 222 | ADAM10 |
| 223 | PPP1R18 |
| 224 | FAM91A1 |
| 225 | WNK1 |
| 226 | MPRIP |
| 227 | ECM1 |
| 228 | DLD |
| 229 | USP39 |
| 230 | PRKRA |
| 231 | PRPF31 |
| 232 | ALDH3A2 |
| 233 | SWAP70 |
| 234 | AAK1 |
| 235 | FAM98C |
| 236 | RELA |
| 237 | ARFGAP1 |
| 238 | UAP1L1 |
| 239 | SNRPG |
| 240 | CC2D1A |
| 241 | SLC9A3R2 |
| 242 | PFDN5 |
| 243 | PTK2 |
| 244 | TRIM21 |
| 245 | NCBP1 |
| 246 | ARIH2 |
| 247 | PANX1 |
| 248 | SRRM1 |
| 249 | DAPK3 |
| 250 | HNRNPLL |
| 251 | SNRPC |
| 252 | HDAC2 |
| 253 | GTPBP1 |

| | |
|---|---|
| 254 | VPS45 |
| 255 | NADK2 |
| 256 | APOOL |
| 257 | GBF1 |
| 258 | GIPC1 |
| 259 | ATE1 |
| 260 | TBC1D2 |
| 261 | RAF1 |
| 262 | EXOC5 |
| 263 | VPS39 |
| 264 | MBNL2 |
| 265 | DNAJC10 |
| 266 | TIGAR |
| 267 | SNTB2 |
| 268 | WDR82 |
| 269 | CYRIB |
| 270 | PTPN14 |
| 271 | STX5 |
| 272 | LSM12 |
| 273 | ATP6V1G1 |
| 274 | CAMK2G |
| 275 | EIF4G3 |
| 276 | PCBP3 |
| 277 | TCEAL3 |
| 278 | COL8A1 |
| 279 | KRAS |
| 280 | TUBAL3 |
| 281 | CPAMD8 |
| 282 | KPNA4 |
| 283 | GIT2 |
| 284 | DCTN6 |
| 285 | PRKAB1 |
| 286 | EML3 |
| 287 | RPS28 |
| 288 | ACOT1 |
| 289 | SRGAP1 |
| 290 | RPP30 |
| 291 | MYD88 |
| 292 | SERPINB7 |
| 293 | VPS4A |
| 294 | C1orf198 |
| 295 | LPAR1 |
| 296 | RPS21 |
| 297 | GNA13 |
| 298 | HSPA14 |
| 299 | HDHD5 |
| 300 | ABRACL |
| 301 | GOPC |
| 302 | CEP120 |
| 303 | NDUFAB1 |
| 304 | CACYBP |

| | |
|---|---|
| 305 | SH3PXD2B |
| 306 | MIA3 |
| 307 | XAGE1A |
| 308 | USP15 |
| 309 | EDEM3 |
| 310 | OSBPL8 |
| 311 | CSNK1A1 |
| 312 | SRSF2 |
| 313 | RIPK2 |
| 314 | CIAO2B |
| 315 | COMMD2 |
| 316 | CCDC22 |
| 317 | MRPL37 |
| 318 | PRPF40A |
| 319 | QKI |
| 320 | ATP5F1D |
| 321 | LARS2 |
| 322 | YARS2 |
| 323 | GHDC |
| 324 | GAK |
| 325 | PCYT2 |
| 326 | GIT1 |
| 327 | CTSA |
| 328 | LANCL1 |
| 329 | PATL1 |
| 330 | CSNK2A2 |
| 331 | LSM4 |
| 332 | MRPS35 |
| 333 | NUP133 |
| 334 | GNE |
| 335 | NRP2 |
| 336 | ACSF3 |
| 337 | SELENOM |
| 338 | ASCC3 |
| 339 | C11orf68 |
| 340 | PAIP1 |
| 341 | PDP1 |
| 342 | PUF60 |
| 343 | NDUFA8 |
| 344 | PGLS |
| 345 | PFDN2 |
| 346 | PRRC2C |
| 347 | SV2A |
| 348 | FHL3 |
| 349 | MAPKAP1 |
| 350 | DHRS4 |
| 351 | PRUNE1 |
| 352 | ARHGEF17 |
| 353 | MRPL55 |
| 354 | UTRN |
| 355 | PELO |

| | |
|---|---|
| 356 | SART3 |
| 357 | PCDH18 |
| 358 | FKBP3 |
| 359 | NUFIP2 |
| 360 | YLPM1 |
| 361 | RBBP7 |
| 362 | UBLCP1 |
| 363 | WIPI1 |
| 364 | TJP2 |
| 365 | UBE2D3 |
| 366 | PTPN12 |
| 367 | SLC16A6 |
| 368 | MTIF2 |
| 369 | POLR2B |
| 370 | EDC3 |
| 371 | CLPP |
| 372 | SRP14 |
| 373 | TP53RK |
| 374 | MIPEP |
| 375 | TMPO |
| 376 | MYO10 |
| 377 | LUZP1 |
| 378 | SCO2 |
| 379 | CNOT3 |
| 380 | NRDC |
| 381 | VPS33B |
| 382 | FRMD8P1 |
| 383 | SKP1 |
| 384 | SRM |
| 385 | TAGLN |
| 386 | TBL1XR1 |
| 387 | MMS19 |
| 388 | DDX49 |
| 389 | NDUFS7 |
| 390 | TSNAX |
| 391 | SCAF4 |
| 392 | CDK2 |
| 393 | FADS3 |
| 394 | XRN1 |
| 395 | RBM26 |
| 396 | PPP2R5D |
| 397 | CASZ1 |
| 398 | GALT |
| 399 | GSDMD |
| 400 | TRIM32 |
| 401 | PQBP1 |
| 402 | ABI1 |
| 403 | ECI2 |
| 404 | INPP4B |
| 405 | DRG2 |
| 406 | FBXL18 |

| | |
|---|---|
| 407 | CHRM5 |
| 408 | ZC3H15 |
| 409 | PRPF4 |
| 410 | SHFL |
| 411 | MTMR1 |
| 412 | ZNF207 |
| 413 | METTL13 |
| 414 | EIF2AK4 |
| 415 | LRCH4 |
| 416 | ANGPTL4 |
| 417 | NRBP1 |
| 418 | GOLGA2 |
| 419 | NOM1 |
| 420 | NEB |
| 421 | UBE3C |
| 422 | STAMBP |
| 423 | MRPL2 |
| 424 | DCAKD |
| 425 | RIC8A |
| 426 | ARL6IP1 |
| 427 | TBL2 |
| 428 | SUMF1 |
| 429 | TTN |
| 430 | COG1 |
| 431 | DECR2 |
| 432 | EIF2B2 |
| 433 | MRPL48 |
| 434 | PPFIA1 |
| 435 | DAGLB |
| 436 | ZNF827 |
